# Molecular changes in agroinfiltrated leaves of *Nicotiana benthamiana* expressing suppressor of silencing P19 and coronavirus-like particles

**DOI:** 10.1101/2024.09.23.614541

**Authors:** Louis-Philippe Hamel, Francis Poirier-Gravel, Marie-Ève Paré, Rachel Tardif, Marc-André Comeau, Pierre-Olivier Lavoie, Andréane Langlois, Marie-Claire Goulet, Dominique Michaud, Marc-André D’Aoust

## Abstract

The production of coronavirus disease 2019 vaccines can be achieved by transient expression of the Spike (S) protein of Severe Acute Respiratory Syndrome Coronavirus 2 in agroinfiltrated leaves of *Nicotiana benthamiana*. Relying on bacterial vector *Agrobacterium tumefaciens*, this process is favored by the co-expression of viral silencing suppressor P19. Upon expression, the S protein enters the cell secretory pathway, before being trafficked to the plasma membrane where formation of coronavirus-like particles (CoVLPs) occurs. We previously characterized effects of influenza virus hemagglutinin forming VLPs through similar processes. However, leaf samples were only collected after six days of expression and it remains unknown whether influenza VLPs and CoVLPs induce similar responses. Here, time course sampling was used to profile responses of *N. benthamiana* leaf cells expressing P19 only, or P19 along with the S protein. The latter triggered early, but transient activation of the unfolded protein response and waves of transcription factor genes involved in immunity. Accordingly, defense genes were induced with different expression kinetics, including those promoting lignification, terpene biosynthesis, and oxidative stress. Crosstalk between stress hormone pathways also occurred, notably leading to the repression of jasmonic acid biosynthesis genes after agroinfiltration, and dampening of salicylic acid-inducible responses upon S protein accumulation. Overall, influenza VLP- and CoVLP-induced responses broadly overlapped, suggesting nanoparticle production to have the most effects on plant immunity, regardless of the virus surface proteins expressed. Taking advantage of RNAseq inferences, we finally show the co-expression of Kunitz trypsin inhibitors to reduce CoVLP-induced defense and leaf symptoms, with no adverse effect on plant productivity.

## Introduction

In December 2019, a cluster of pneumonia cases emerged in Wuhan, a city from the province of Hubei in China. Sequencing efforts revealed these cases to be caused by a novel coronavirus termed 2019-nCoV (Zhou *et al*., 2020; Zhu *et al*., 2020). Later renamed Severe Acute Respiratory Syndrome Coronavirus 2 (SARS-CoV-2), this pathogen turned out to be highly contagious and the associated coronavirus disease 2019 (COVID-19) rapidly spread throughout the world (see Appendix S1 for a definition of all abbreviations). In March 2020, COVID-19 was declared a pandemic by the World Health Organization (WHO). In the absence of an efficient vaccine or treatment, health authorities from many countries introduced measures to limit spreading of the disease, including the use of surgical masks, social distancing, and the closing of non-essential institutions. While these measures reduced virus transmission, they had serious societal and economic burdens that led to their uneven acceptation and application. In turn, COVID-19 kept spreading and as of September 2024, the original SARS-CoV-2 strain Wuhan-Hu-1 and its variants had caused over 776 million of confirmed COVID-19 cases and more than 7 million deaths worldwide (https://covid19.who.int).

Vaccination is broadly recognized as one of the most effective methods to prevent viral infections and the public health emergency raised by COVID-19 sparked unprecedented efforts to develop protective vaccines, including through traditional approaches such as live attenuated or inactivated viruses, peptides, or adjuvanted recombinant proteins. Novel manufacturing platforms also emerged, including those based on mRNA, DNA, as well as replicating and non-replicating viral vectors (Funk *et al*., 2021). Clinical trials conducted early during the pandemic confirmed the efficacy of mRNA-based vaccines (Polack *et al*., 2020; Baden *et al*., 2021). Since then, additional vaccines with suitable safety and efficacy profiles were successfully deployed (Rotshild *et al*., 2021).

Thus far, the development of COVID-19 vaccines has focused on primary target of the host humoral response, the SARS-CoV-2 Spike (S) protein. Anchored in the viral membrane through a transmembrane domain (TMD), this glycoprotein forms homotrimers at the surface of virions (Wrapp *et al*., 2020). As it interacts with the host cell-surface receptor Angiotensin-Converting Enzyme 2 (ACE2) (Zhou *et al*., 2020), the S protein plays an essential role in virus attachment and entry into host mammalian cells (Jackson *et al*., 2022). Serological analyses indicate that most neutralizing antibodies produced by infected hosts target the receptor-binding domain (RBD) of the S protein, blocking its interaction with ACE2 (Piccoli *et al*., 2020; Li *et al*., 2022). As cellular immunity facilitates patient recovery in addition of maximizing long-term protection against onward infections (Sette and Crotty, 2021), an ideal COVID-19 vaccine should induce robust humoral and T cell responses in immunized hosts.

Transient protein expression mediated by *Rhizobium radiobacter* (commonly known and hereafter referred to as *Agrobacterium tumefaciens*) is a powerful approach to express recombinant proteins in plants (Sainsbury and Lomonossoff, 2014). Typically performed in leaves of the model plant *Nicotiana benthamiana* (Bally *et al*., 2018), *Agrobacterium*-mediated expression makes use of plant cells as ‘protein biofactories’, including for the manufacturing of biopharmaceutical products (Chung *et al*., 2022). Easily scalable, leaf agroinfiltration is an alternative to classical protein expression systems that is now seen as a helpful and viable option in the fight against global health issues, including endemic and new infectious diseases (Anon., 2022). Using agroinfiltration in *N. benthamiana* leaves, the biopharmaceutical company Medicago for instance developed influenza vaccine candidates by transiently expressing recombinant hemagglutinins (HAs) that induce the formation of virus-like particles (VLPs) in host plant cells. Devoid of other viral proteins or genetic components required for replication (Landry *et al*., 2010), VLPs accumulate in the virtual space located between the plant cell plasma membrane (PM) and the cell wall (D’Aoust *et al*., 2008). In clinical trials, vaccine candidates against pandemic (monovalent) and seasonal (quadrivalent) influenza virus strains were shown to induce balanced humoral and T cell responses against influenza viruses in humans (Landry *et al*., 2014; Pillet *et al*., 2016; 2019; Ward *et al*., 2020).

In response to the rapid and global spreading of COVID-19, Medicago employed a similar approach to develop a vaccine candidate made of coronavirus-like particles (CoVLPs). These self-assembling VLPs consist of a lipid envelope derived from the PM of plant cells and trimer clusters of recombinant S protein in a stabilized, pre-fusion conformation. Derived from the original SARS-CoV-2 strain Wuhan-Hu-1, the recombinant S protein was further engineered to enter the secretory pathway of plant cells more efficiently and to maximize budding of CoVLPs. Transmission electron microscopy (TEM) revealed CoVLPs to be nanoparticles similar in size and shape to SARS-CoV-2 virions, including a lipid envelope in which S protein trimers were anchored at high density (Ward *et al*., 2021). In clinical trials, the vaccine induced strong and durable neutralizing antibody and T-cell responses against SARS-CoV-2 (Gobeil *et al*., 2021; Ward *et al*., 2021). Vaccine efficacy ranged between 69.5% for any symptomatic COVID- 19 and 78.8% for moderate to severe disease cases (Hager *et al*., 2022). This vaccine, named Covifenz^®^, was authorized for human use in Canada on February 24, 2022, under Food and Drug Regulations of Health Canada.

To prevent silencing of the *S* transgene delivered by *Agrobacterium*, the S protein was co-expressed with P19, a viral suppressor of RNA silencing (Silhavy *et al*., 2002). In host plants, this artificial expression system unavoidably causes an intricate array of stresses, including mechanical stimulation caused by the agroinfiltration process in itself, bacterium perception by leaf cells, and temporal accumulation of the foreign proteins. When expressing virus surface proteins such as recombinant HAs or the S protein, nanoparticle budding also affects the PM, which supplies lipids for VLP (or CoVLP) envelope makeup. We recently reported that expression of influenza HA induces a unique molecular signature in plant cells at six days post-infiltration (DPI), including the activation of immune responses such as oxidative stress and oxylipin-mediated signalling (Hamel *et al*., 2024a). Time course sampling further revealed these immune responses to be preceded by activation of the unfolded protein response (UPR), a transient phenomenon suggested to favor HA expression in *N. benthamiana* leaves (Hamel *et al*., 2024b).

In this study, we used time course sampling over several days to assess the molecular responses of *N. benthamiana* leaf cells solely expressing silencing suppressor P19, or co-expressing P19 in combination with the S protein. Results showed the latter to trigger a myriad of immune responses with different kinetics and crosstalk between the salicylic acid (SA) and the jasmonic acid (JA) signalling pathways. Most importantly, molecular responses induced by influenza VLPs and CoVLPs largely overlapped, suggesting nanoparticle production to have the most effects on plant immunity, regardless of the virus surface protein expressed. This data significantly expands our understanding of plant molecular responses to foreign protein expression and it will be helpful in coming years to improve molecular farming practices, as here exemplified by the co-expression of Kunitz trypsin inhibitors (KTIs) that reduced CoVLP-induced defenses and leaf symptoms, with no adverse effect on S protein accumulation *in planta*.

## Results

### Stress symptoms and recombinant protein expression

To define molecular responses induced by the heterologous production of CoVLPs in *N. benthamiana*, recombinant S protein was co-expressed along with P19 in leaf tissues (a condition hereafter referred to as CoVLP plants, or CoVLP samples). At 6 DPI, the most obvious stress symptoms consisted of greyish necrotic flecking of the leaf blade and curling of leaf blade edges (Figure 1a). Within the canopy, these symptoms were observed on specific leaves of the main stem (referred to as primary, or P, leaves), although the leaves of some axillary stems (referred to as secondary, or S, leaves) were also affected (see asterisks in Figure S1a).

**Figure 1.**
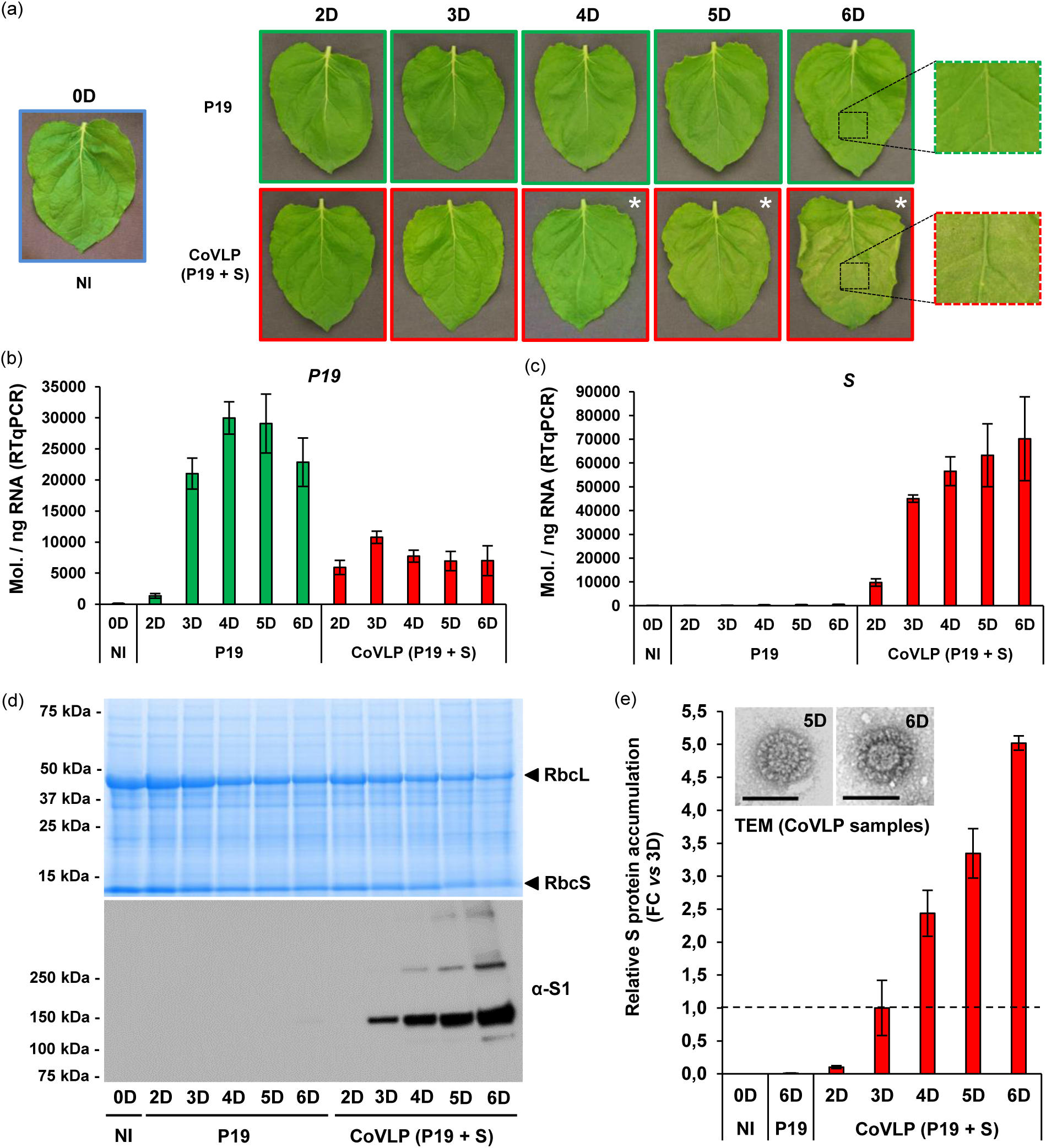
Stress symptoms and S protein expression. (a) Stress symptoms observed on representative leaves of each condition at various time points in days (D) post-infiltration. Based on primary leaf numbering (Figure S1a), pictures either highlight P9 or P10 leaves and asterisks denote visible necrotic lesions. On the right, magnified leaf sections highlight differences in leaf symptoms at the end of sampling. Expression of recombinant genes *P19* (b) and *S* (c) as measured by RTqPCR. Results are expressed in numbers of transcript molecules per ng of RNA ± se. (d) Total protein extracts after SDS-PAGE and Coomassie blue-staining (upper panel). Arrows highlight RuBisCO small (RbcS) and large (RbcL) subunits. The western blot (lower panel) depicts S protein accumulation. (e) Relative S protein accumulation as measured by standardized capillary western immunoassays. S protein accumulation in CoVLP samples at 3 DPI was arbitrarily set at one-fold (dashed line). At the top, TEM imaging confirms the production of CoVLPs at 5 and 6 DPI. Scale bars equal 100 nm. Conditions are as follows: NI: non-infiltrated samples; P19: agroinfiltrated samples expressing P19 only; CoVLP: agroinfiltrated samples co-expressing P19 and recombinant S protein.

Importantly, symptom distribution was strictly conserved when multiple CoVLP plants were compared. To better characterize the distribution of leaf symptoms, P leaves were numbered according to their respective position along the main stem, starting from the bottom and moving up towards the apex (P1 to P12; left panels of Figure S1a). Similarly, S leaf bundles were numbered based on the position of their respective axillary stem along the plant main stem (S1 to S10; right panel of Figure S1a). Based on this numbering, most symptoms were scored on leaves P8, P9, and P10 (Figure S1a), suggesting uneven stress levels due to differential accumulation of the S protein in the plant.

To confirm this, standardized capillary western immunoassays were conducted to quantify the S protein in five P and five S leaf sections (Figure S1b). Based on S protein level per weight unit of leaf biomass, canopy section P9/P10 was the most efficient at producing the viral surface protein (left panel of Figure S1b). After weighing the biomass of each leaf section, their net contribution to the overall S protein yield was also estimated. In brief, P leaves accounted for ∼80% of the total S protein yield, with sections P7/P8 and P9/P10 each accounting for more than 30% of the whole production (right panel of Figure S1b). As the uneven S protein distribution at the plant scale matched the distribution of leaf symptoms (Figure S1a), we selectively harvested leaves P9 and P10 to further assess the plant’s responses to CoVLP expression.

The leaves of CoVLP plants were harvested after 2, 3, 4, 5, and 6 days of expression. As the effects of *Agrobacterium* infiltration (with no recombinant protein expressed) were previously shown to largely overlap with the effects of *Agrobacterium*-mediated expression of P19 only (Hamel *et al*., 2024a), P9 and P10 leaves of agroinfiltrated plants solely expressing P19 were used as controls at each time point (P19 samples). The P9 and P10 leaves of non-infiltrated (NI) plants harvested prior to infiltration (0 DPI) were also analysed (NI samples), as an initial reference point.

The effects of foreign protein accumulation were first characterized by a macroscopic evaluation of the leaf symptoms at each time point. Using NI leaves at 0 DPI as a baseline, no obvious effect was visible on agroinfiltrated leaves harvested at 2 or 3 DPI, regardless of the recombinant protein expressed (Figure 1a). At 4 DPI, P19 leaves had started to show faint chlorosis, as revealed by yellowish discoloration of the leaf blade. At later time points, leaf yellowing had expanded, however no evidence of plant cell death (PCD) was denoted. In comparison, leaves co-expressing P19 and the S protein showed more pronounced chlorosis, accompanied by greyish necrotic flecking that became visible between 3 and 4 DPI (see asterisks in Figure 1a). Cell death then evenly expanded throughout the leaf blade and curling of leaf blade edges was clearly visible at 6 DPI (Figure 1a).

Real time quantitative polymerase chain reaction (RTqPCR) was next performed to confirm expression of the *P19* and *S* transgenes in P9 and P10 leaves. As expected, no transcripts of these genes were detected in NI samples at 0 DPI (*P19* in Figure 1b and *S* in Figure 1c). For *P19*, low transcript levels were detected in P19 samples at 2 DPI and expression gradually increased up to 4 DPI, reaching a maximum level of ∼30,000 transcript molecules per ng of RNA (Figure 1b). *P19* expression was pretty much maintained up to 5 DPI, before going down slightly at 6 DPI. *P19* expression was also detected in CoVLP samples, with a maximum level of transcripts at 3 DPI and reaching a little less than 11,000 transcript molecules per ng of RNA (Figure 1b). *P19* expression then slightly decreased and remained stable until the end of sampling at 6 DPI. As inferred from P19 samples, co-expression of the *S* gene reduced the expression of *P19* (Figure 1b), similar to the previously reported negative impact of influenza *HA* on expression of this silencing suppressor gene (Hamel *et al*., 2024a). As expected, no *S* gene transcript was detected in P19 samples (Figure 1c). By contrast, low expression of the *S* gene was detected at 2 DPI in CoVLP samples. *S* gene expression then notably increased at 3 DPI and kept doing so until the end of sampling to reach a maximum level, just over 70,000 transcript molecules per ng of RNA. When comparing expression levels of the *P19* and *S* genes, the latter was expressed at higher levels (Figure 1b,c), consistent with the relative strength of promoters used to drive transgene expression, namely a *plastocyanin* promoter for *P19* and a *2X35S* promoter for *S*.

To monitor S protein accumulation, total protein extracts were prepared from NI, P19, and CoVLP leaf samples. SDS-PAGE and Coomassie blue-staining confirmed integrity of the proteins in crude extracts (upper panel of Figure 1d), including those retrieved from the most symptomatic leaves (Figure 1a). As the expression phase progressed, a gradual reduction in levels of the ribulose-1,5- bisphosphate carboxylase/oxygenase (RuBisCO) large and small subunits was observed, with more pronounced effects in CoVLP samples than in P19 samples (upper panel of Figure 1d). From the same protein extracts, a western blot was performed in non-reducing conditions, using an antibody specific to the S1 domain of the SARS-CoV-2 S protein. For all time points, no signal was detected in NI samples, nor in P19 samples (lower panel of Figure 1d). By contrast, a clear signal was detectable in CoVLP samples at 3 DPI and the signal later increased until the end of sampling at 6 DPI.

Based on its polypeptide sequence, the predicted molecular weight of full-length recombinant S protein is ∼140 kDa for its non-mature form and ∼137 kDa after removal of the signal peptide (SP). As the strongest immunoblot signals were detected at ∼165 kDa (lower panel of Figure 1d), the main recombinant product found in leaf sample extracts presumably corresponded to the complete, monomeric and glycosylated form of the recombinant S protein. Under nonreducing conditions, larger bands corresponding to dimeric and trimeric forms of the S protein were also detected. A smaller band of ∼135 kDa, mostly detected at 6 DPI, was also visible (lower panel of Figure 1d), likely corresponding to a stable proteolytic fragment generated after clipping of the S protein by one or several host plant cell endoprotease(s).

Using standardized capillary western immunoassays, the relative accumulation of the S protein was also determined at each time point. To this end, S protein level of CoVLP samples at 3 DPI was arbitrarily set at one-fold (see dashed line in Figure 1e). Consistent with the immunoblotting signals (lower panel of Figure 1d), no accumulation of the S protein was detected in NI samples at 0 DPI or in P19 samples at 6 DPI. By contrast, barely detectable level of the S protein was detected in CoVLP samples at 2 DPI and S protein accumulation quickly picked-up between 2 and 3 DPI. Past that point, S protein accumulation kept increasing until the end of sampling, eventually increasing by more than five-fold compared to the baseline level at 3 DPI (Figure 1e). CoVLPs were next partially purified using biomass from CoVLP samples at 5 and 6 DPI. TEM imaging of the resulting products revealed nanoparticles to resemble SARS-CoV-2 virions both in size and morphology (see pictures; Figure 1e). Overall, our preliminary analyses confirmed that harvested leaf biomass was suitable for downstream analyses of the leaf transcriptomes.

### Effects of foreign protein expression on the leaf transcriptome

To define transcriptional changes in P19- and COVLP-expressing leaves, a systematic RNAseq survey was performed with RNA material derived from the NI, P19, and CoVLP samples described above. A principal component analysis (PCA) revealed a tight clustering of biological replicates from a defined condition at a given time point (Figure 2a). On the other hand, transcriptional profiles markedly differed depending on the recombinant proteins expressed and the timing at which leaf sampling was performed. Using NI leaves at 0 DPI as a control, pairwise comparisons were performed for P19 and CoVLP samples harvested at 2, 3, and 6 DPI. These time points were selected based on the S protein accumulation pattern (Figure 1d,e), to highlight transcriptional changes in the presence of a barely detectable S protein level (2 DPI), presence of a low S protein level (3 DPI), or presence of a high S protein level at the end of the sampling period (6 DPI). To be considered as significantly regulated, a given gene had to fulfill the following criteria: a Log_2_ of the expression fold change value (Log_2_FC) ≥ 2 or ≤ -2, and an adjusted p-value (padj) < 0.1 indicating a false discovery rate (fdr) below 10%. For each selected time point, unsorted lists of significantly up- and downregulated genes are available in the Supporting Information online (Tables S1, S2, and S3, respectively). Based on the established expression thresholds, volcano plots highlighting all sequenced genes were created (Figure 2b), while pairwise comparisons were exploited to create Venn diagrams for the up- and downregulated genes (Figure 2c). For each section of the Venn diagrams, unsorted lists of up- and downregulated genes at 2, 3, and 6 DPI are available in the Supporting Information online (Tables S4, S5, and S6, respectively). For P19 and CoVLP samples, these analyses showed the number of regulated genes to increase as the expression phase progressed (Figure 2b,c). For a defined sampling point, regulated genes were however more abundant in CoVLP samples than in P19 samples, except for the downregulated genes at 2 DPI (Figure 2c).

**Figure 2.**
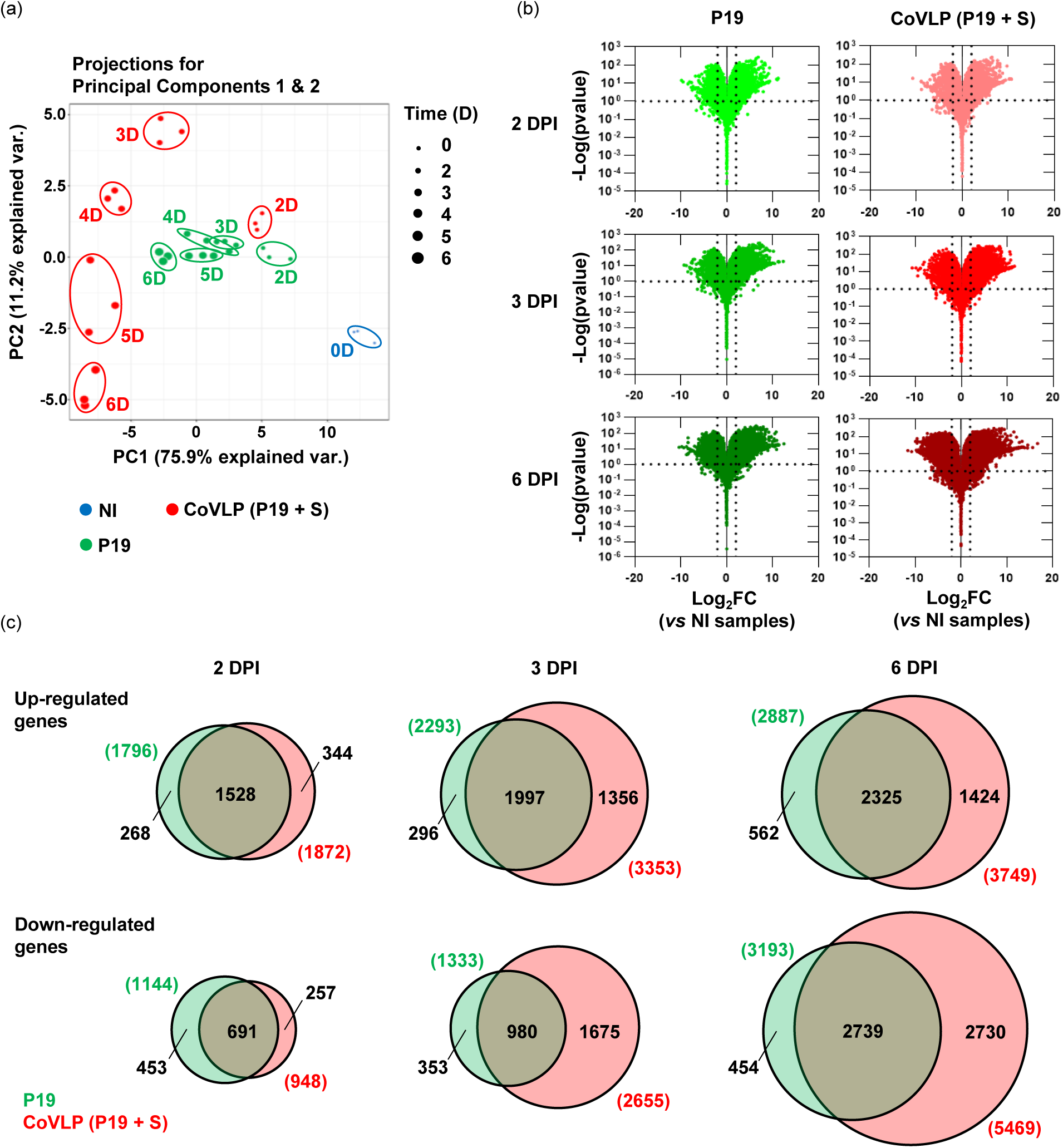
Overview of the RNAseq results. (a) PCA summarizing the effects of recombinant protein expression on the leaf transcriptomes. The 50 genes showing highest variance were employed, excluding photosynthesis-related genes. The first axis (PC1) explains 75,9% of the variance (var.), the second axis (PC2) 11,2%. NI samples are shown in blue, P19 samples in green, and CoVLP samples in red. Size of the dots reflects sampling points in days (D) post-infiltration. (b) Global transcriptional changes as depicted by volcano plots of all sequenced genes at 2 days post-infiltration (DPI; upper panels), 3 DPI (middle panels), and 6 DPI (lower panels). Dashed lines represent expression thresholds: Log_2_FC ≥ 2 or ≤ -2 and adjusted p-value (padj) < 0.1. (c) Venn diagrams depicting the numbers of up- and downregulated genes in P19 and CoVLP samples at 2, 3, and 6 DPI. Genes were extracted from pairwise comparisons between P19 and NI samples, or CoVLP and NI samples. Circle size is proportional to the number of regulated genes. Genes specific to the P19 samples are shown in green, those specific to the CoVLP samples in red. Diagram intersects show genes common to both conditions. Total numbers of regulated genes are shown in parenthesis.

For CoVLP samples at 2 DPI, a total of 1,872 genes were upregulated, including 1,528 genes (∼82%) shared with P19 samples (Figure 2c). At 2 DPI, CoVLP samples also revealed 948 downregulated genes, including 691 genes (∼73%) shared with P19 samples. At 3 DPI, CoVLP samples displayed 3,353 upregulated genes, including 1,997 genes (∼60%) shared with P19 samples (Figure 2c). On the opposite, 2,655 genes were downregulated, including 980 genes (∼37%) shared with P19 samples. Consistent with levels of S protein at 2 and 3 DPI (Figure 1d,e), a decreased proportion of shared genes at 3 DPI indicated that P19 and CoVLP samples had much closer transcriptomes at 2 DPI. This was consistent with the PCA, which showed P19 and CoVLP samples at 2 DPI to stand much closer to each other compared to the corresponding samples at 3 DPI (Figure 2a).

For CoVLP samples at 6 DPI, our RNAseq data revealed 3,749 upregulated genes, including 2,325 genes (∼62%) also upregulated in P19 samples (Figure 2c). At the end of sampling, CoVLP samples also displayed 5,469 downregulated genes, half of which (2,739 genes) were also repressed in P19 samples. Interestingly, the proportion of upregulated genes shared by both conditions was roughly the same at 3 and 6 DPI (∼60%), suggesting many of the CoVLP-induced genes to be upregulated shortly after the onset of S protein accumulation (Figure 1d,e). This idea was supported by the overall numbers of upregulated genes, which only slightly increased between 3 and 6 DPI while almost doubling in the shorter time lapse between 2 and 3 DPI (Figure 2c). Taken together, RNAseq showed expression of the S protein to result in the induction of a unique molecular signature increasing in intensity and complexity as the expression phase progressed. Whereas many differentially expressed genes were specific to CoVLP samples, others were also differentially expressed in P19 samples. For those genes shared by the two conditions, the extent of up- or downregulation was however generally much higher when the S protein was expressed (Tables S4, S5, and S6). Overall, the increased transcriptional changes were consistent with the intensity of the leaf symptoms, which were also more pronounced on the CoVLP leaves (Figure 1a).

### RTqPCR validation of RNAseq data

RNAseq on time course samples sought to profile plant gene expression as a function of sampling time. To validate our RNAseq data, RTqPCR was performed on the RNA extracts used for the RNAseq analyses. Following amplification, normalized numbers of transcript molecules per ng of RNA were equally plotted against sampling time. The expression profiles deduced from either approaches were then compared head-to-head, with linear regression (LR) and coefficient of determination (R^2^) values calculated for each tested gene (Figure S2a). Through this approach, 20 plant genes were analysed (Figure S2b), including genes with different functions, expression levels, or transcriptional behaviors in the tested conditions (Figure S2a). Overall, calculated LR and R^2^ values confirmed a strong correlation between the datasets produced from the two independent methods (Figure S2b). While confirming the robustness and validity of our RNAseq data, these results also prompted the use of RNAseq profiles to further assess specific cellular and metabolic processes affected by foreign protein expression in the host plant cells.

### Upregulation of UPR-associated genes

In eukaryotes, the UPR allows cells to cope with endoplasmic reticulum (ER) stress induced by accumulation of misfolded proteins in the ER (Read and Schröder, 2021). In plants, the UPR comprises two signalling branches, each including conserved components and activation mechanisms (Howell, 2021). We recently identified several genes involved in the UPR of *N. benthamiana*, including those encoding transcription factors (TFs) of the basic leucine zipper (bZIP) family, as well as those encoding components of the ER quality control (ERQC) and ER-associated degradation (ERAD) systems (Hamel *et al*., 2024b). Here, RNAseq revealed foreign protein expression to induce functionally diverse UPR genes, with upregulation levels higher in CoVLP samples than in P19 samples (Table S7). Upon S protein expression, UPR genes displayed remarkedly uniform expression patterns, with maximum upregulation levels reached at 3 or 4 DPI. UPR genes upregulated included *NbbZIP60* (Niben101Scf24096g00018) (Figure 3a), a TF-encoding gene that acts as a master switch for UPR activation. Also induced were ERQC-associated genes encoding protein disulfide isomerases (PDIs) and ER-resident chaperones of the Binding immunoglobulin Protein (BiP), calnexin (CNX), and calreticulin (CRT) families (Figure 3b). Similarly, early and transient upregulation was observed for ERAD-associated genes (Table S7), including those encoding the luminal lectin osteosarcoma 9 (NbOS9) (Niben101Scf01427g01001), ER-resident chaperone glucose-regulated protein 94 (NbGRP94) (Niben101Scf04331g09018), and several components of the retrotranslocation complex (Figure 3c). Taken together, these results suggest S protein expression in *N. benthamiana* leaves to result in early and transient activation of the UPR.

**Figure 3.**
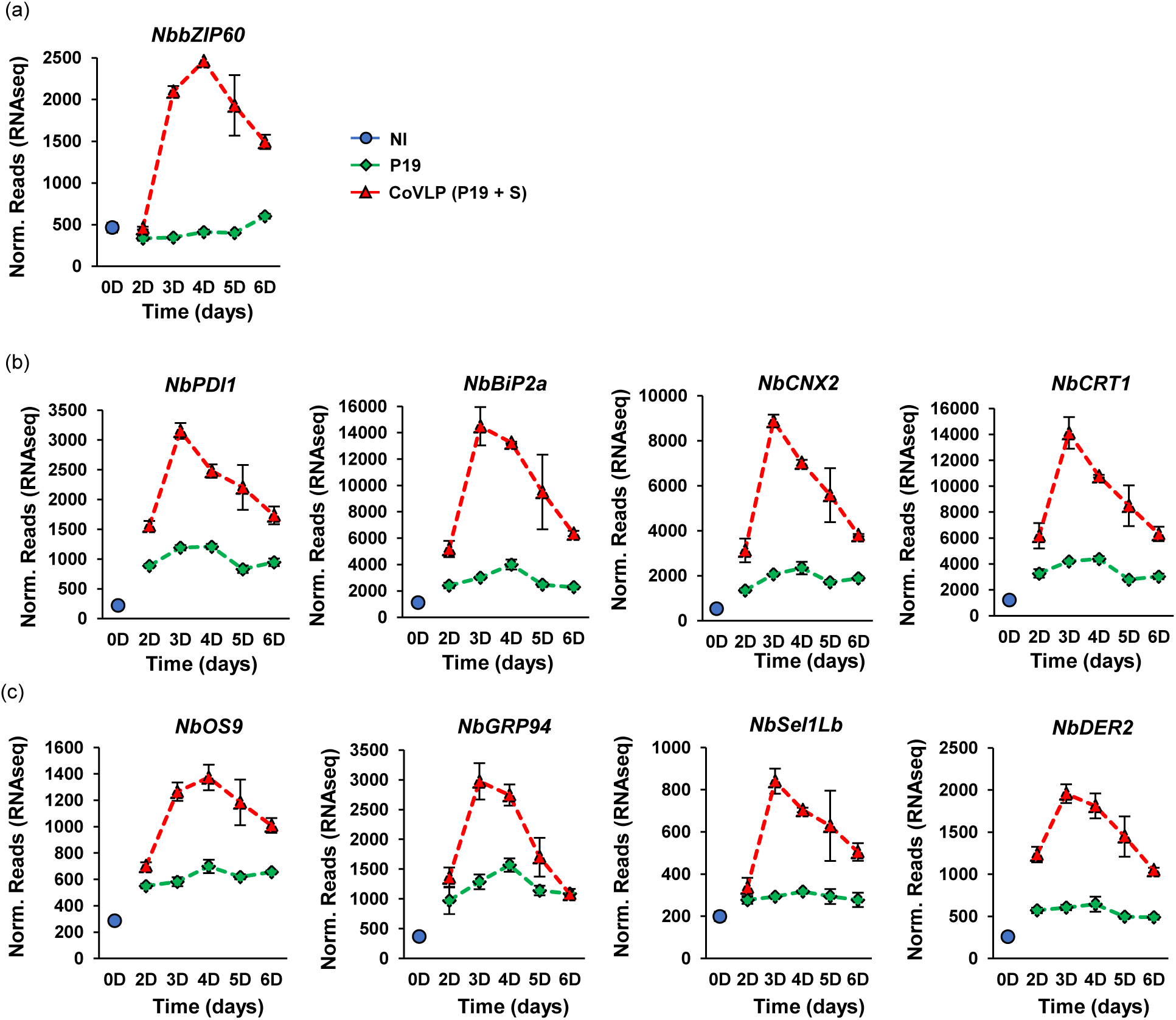
Upregulation of ER stress-related genes. Expression profiles of TF gene *NbbZIP60* (a), ERQC genes (b), and ERAD genes (c). For each time point in days (D) post-infiltration, RNAseq results are expressed in normalized (norm.) numbers of reads ± sd. Conditions are as follows: NI: non-infiltrated samples (blue); P19: agroinfiltrated samples expressing P19 only (green); CoVLP: agroinfiltrated samples co-expressing P19 and recombinant S protein (red).

### Upregulation of TF-encoding genes

To make sense of the transcriptional changes in leaves expressing foreign proteins (Figure 2), we examined the regulation of TF-encoding genes. Among the induced candidates, many belonged to TF families involved in plant immunity (Table S8). Generally, TF gene upregulation followed two predominant expression patterns. The so-called ‘early pattern’ was characterized by rapid gene upregulation peaking no later than 4 DPI, before going down at later time points. By contrast, the so-called ‘late pattern’ was characterized by gene upregulation not detected prior to 3 or 4 DPI, but rapidly increasing at later time points. In many cases, maximum expression level of late TF genes was not reached by the end of sampling at 6 DPI. The manifestation of at least two TF gene ‘waves’ implies a certain hierarchy among induced transcriptional regulators, perhaps to control early and late immune responses during foreign protein expression.

Among upregulated TF genes, many belonged to the *NO APICAL MERISTEM* (*NAM*), *ARABIDOPSIS ACTIVATION FACTOR 1* (*ATAF1*), and *CUP SHAPED COTYLEDON 2* (*CUP2*) family, more simply referred to as *NACs* (Table S8). With more than 100 members in *Arabidopsis thaliana*, NACs form one of the largest TF families in plants (Ooka *et al*., 2003). Among the early *NACs* identified, several encoded trans-membrane proteins homologous to membrane-tethered NACs in Arabidopsis (Kim *et al*., 2007). These included the *N. benthamiana* gene *NAC Targeted by Phytophthora 1* (*NbNTP1*) (Niben101Scf00308g01003), a defense-related TF whose translocation from the ER membrane to the nucleus is inhibited by a pathogen effector (McLellan *et al*., 2013). Upregulation of *NbNTP1* peaked at 3 DPI and was essentially specific to CoVLP samples (Figure 4a). Early membrane-tethered *NACs* also included the closely related genes *NbNAC62* (Niben101Scf27945g00015) and *NbNAC91* (Niben101Scf04099g08003), as well as a more distant homolog, *NbNAC103* (Niben101Scf01795g17007). Named after their closest homolog in Arabidopsis (AtNAC62, AtNAC91, and AtNAC103, respectively), these genes shared similar expression profiles following agroinfiltration, with much stronger upregulation levels in CoVLP samples (Figure 4a). Interestingly, Arabidopsis homologs of these genes were reported to be involved in activation of the UPR, and their expression to require the upstream activation of AtbZIP60 (Sun *et al*., 2013; Yang *et al*., 2014, 2023).

**Figure 4.**
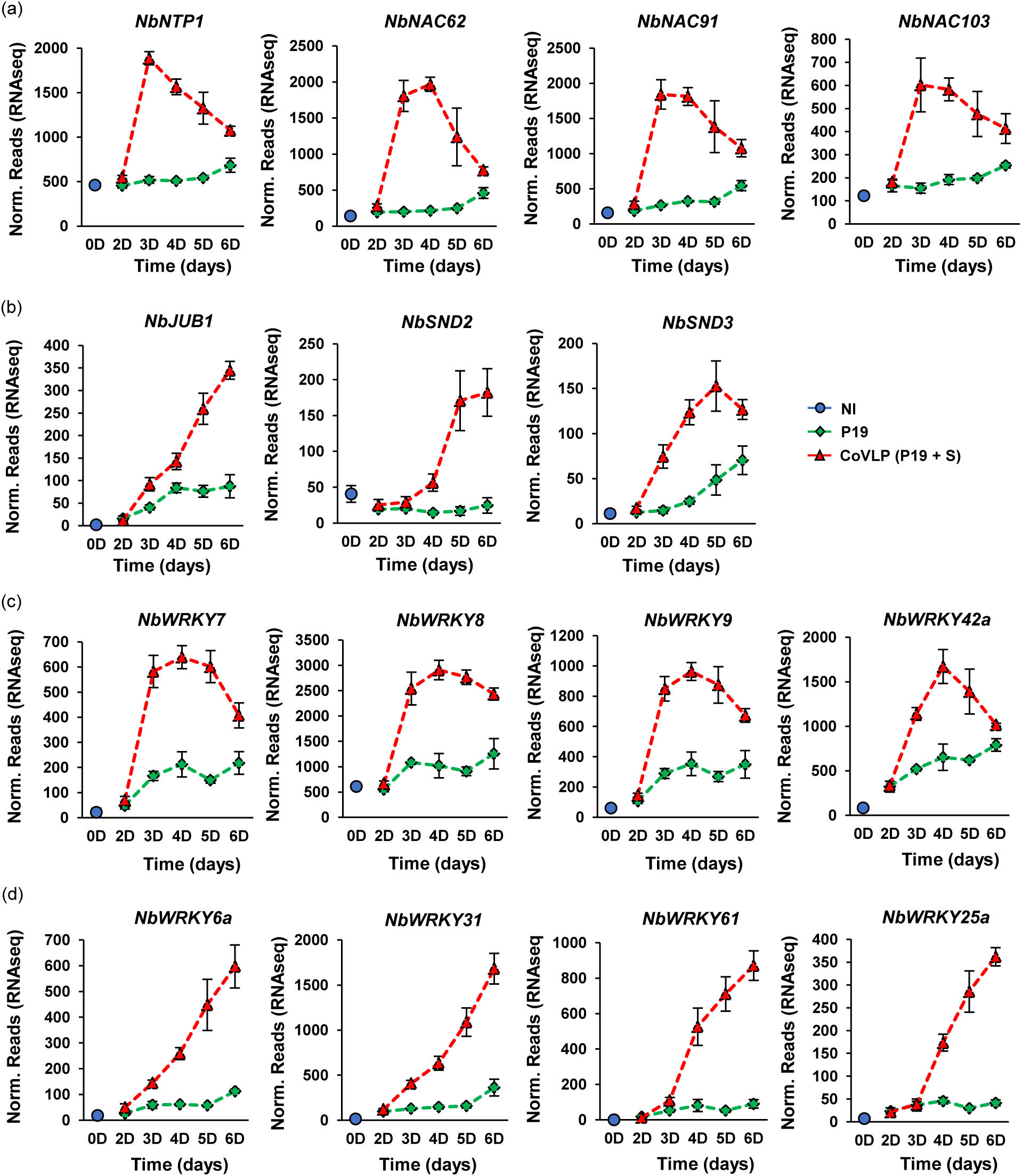
Upregulation of *NAC* and *WRKY* TF genes. Expression profiles of genes encoding early (a) and late (b) NACs. Expression profiles of genes encoding early (c) and late (d) WRKYs are also shown. For each time point in days (D) post-infiltration, RNAseq results are expressed in normalized (norm.) numbers of reads ± sd. Conditions are as follows: NI: non-infiltrated samples (blue); P19: agroinfiltrated samples expressing P19 only (green); CoVLP: agroinfiltrated samples co-expressing P19 and recombinant S protein (red).

Our RNAseq dataset also identified late *NACs*, many of which were specific to, or at least more strongly induced in CoVLP samples (Table S8). These included *NbJUB1* (Niben101Scf07673g04009) (Figure 4b), which is most similar to the Arabidopsis gene *JUNGBRUNNEN1* (*AtJUB1*). Strongly induced by reactive oxygen species (ROS), this TF favors cell longevity by controlling ROS cellular levels (Wu *et al*., 2012). Late *NACs* also included closely related genes *NbSND2* (Niben101Scf11552g02001) and *NbSND3* (Niben101Scf03169g04009) (Figure 4b), which are close homologs of Arabidopsis *SECONDARY WALL-ASSOCIATED NAC DOMAIN 2* (*AtSND2*) and *AtSND3*, respectively. As key components of a transcriptional network controlling secondary cell wall thickening, these TFs promote the expression of cellulosic, hemicellulosic, and lignin-related genes (Zhong *et al*., 2008). In many plant species, including Arabidopsis and *Nicotiana tabacum* (tobacco), NACs are also known to promote leaf senescence (Li *et al*., 2018). In a manner largely specific to CoVLP samples, homologs of senescence-promoting *NACs* were also upregulated (Table S8), including *NbNAC2* (Niben101Scf00799g00011), *NbNAC29* (Niben101Scf10322g01012), *NbNAC81* (Niben101Scf15435g00012), *NbNAC87* (Niben101Scf02762g01001), and *NbORE1* (Niben101Scf01277g00002). Respectively named after their closest homologs in Arabidopsis, those NACs thought to promote leaf senescence will be discussed in our accompanying manuscript (Hamel *et al*., 2025).

Along with NACs, proteins of the WRKY family constitute one of the largest groups of TFs in plants (Bakshi and Oelmüller, 2014). During foreign protein expression, RNAseq identified a large number of *WRKY* genes, many of which were specific to, or at least more strongly induced in CoVLP samples compared to P19 samples (Table S8). Among the early *WRKYs* identified (Figure 4c), three related candidates showed very similar expression profiles with maximum upregulation levels pretty much maintained between 3 and 5 DPI. Formerly termed *NbWRKY7* (Niben101Scf01297g04006), *NbWRKY8* (Niben101Scf02362g02014), and *NbWRKY9* (Niben101Scf02430g03006), these genes encode TFs that are phosphorylated by mitogen-activated protein kinases (MAPKs). Interestingly, phosphorylation increases target gene expression, including a 3-hydroxy-3- methylglutaryl-CoA reductase (HMGR)-encoding gene involved in terpene biosynthesis (Ishihama *et al*., 2011) and an NADPH oxidase-encoding gene involved in the activation of oxidative stress (Adachi *et al*., 2015). In Arabidopsis, functional studies also revealed a close homolog, AtWRKY33, to promote JA signalling at the expense of SA signalling (Zheng *et al*., 2006). Also upregulated early were two closely related genes, *NbWRKY42a* (Niben101Scf03786g04014) and *NbWRKY42b* (Niben101Scf04011g05006). Sharing similar expression profiles, with maximum expression levels reached at 4 DPI (Table S8; Figure 4c), these genes were named after their closest Arabidopsis homolog, AtWRKY42, a TF that promotes ROS production by favoring the expression of an NADPH oxidase-encoding gene (Niu *et al*., 2020).

Late *WRKYs* were also upregulated in our tested conditions (Table S8), including closely related genes *NbWRKY6a* (Niben101Scf02516g01004), *NbWRKY6b* (Niben101Scf02145g09012), and *NbWRKY31* (Niben101Scf02170g02011).

Named after AtWRKY6 and AtWRKY31, these genes were again induced more strongly in CoVLP samples compared to P19 samples (Figure 4d). While the biological functions of AtWRKY31 remain to be determined, AtWRKY6 was shown to activate plant immunity and leaf senescence (Robatzek and Somssich, 2001). Because they were highly induced in CoVLP samples compared to NI and even P19 controls, *NbWRKY61* (Niben101Scf09644g00009) and *NbWRKY72* (Niben101Scf24496g01001) were also of particular interest (Table S8; Figure 4d). Closely related to each other, these genes are somewhat related to *AtWRKY61* and *AtWRKY72*, although with lower homology levels compared to other *WRKYs* named after their closest homolog in Arabidopsis. Late *WRKYs* finally included a small group of closely related genes formerly termed *NbWRKY22a* (Niben101Scf09512g02006), *NbWRKY22b* (Niben101Scf03239g00001), *NbWRKY25a* (Niben101Scf04871g19004), and *NbWRKY25b* (Niben101Scf09057g01009) (Table S8; Figure 4d). Acting as positive regulators of immune responses in *N. benthamiana* (Ramos *et al*., 2021), the products of these genes are closely related to AtWRKY22 and AtWRKY29. Both of these Arabidopsis TFs are MAPKs substrates (Asai *et al*., 2002), with AtWRKY22 reported to promote leaf senescence (Zhou *et al*., 2011).

*NAC* and *WRKY* genes were the most prominent TF families upregulated upon foreign protein expression. Nevertheless, TF-encoding genes from other families with known roles in plant defense were also identified, especially in leaf samples expressing the S protein (Table S8). Among those induced genes, many encoded early or late APETALA2/ETHYLENE RESPONSIVE FACTOR (AP2/ERF) (Figure S3a), MYB (Figure S3b), bZIP (Figure S3c), or GRAS (Figure S3d) TFs.

### Upregulation of lignin-related genes

In response to pathogens, plant cells reinforce their cell wall through the coordinated deposition of polymers such as lignin (Wang *et al*., 2013). Expression of the S protein induced several genes involved in monolignol biosynthesis (Table S9), the precursors of lignin polymers. These included the *hydroxycinnamoyl-CoA transferase* gene *NbHCT1* (Niben101Scf09928g01001), the *coumarate-3- hydroxylase* gene *NbC3Ha* (Niben101Scf00077g03005), the *caffeoyl-CoA O-methyltransferase* gene *NbCCoAOMT1* (Niben101Scf14852g00006), and the *caffeic acid 3-O-methyltransferase* gene *NbCOMT1* (Niben101Scf02026g02001) (Figure 5a). The upregulation of monolignol biosynthesis genes started early, peaked at 4 DPI, and later decreased until the end of sampling at 6 DPI. Among genes that were the most strongly induced in CoVLP samples, several encoded secreted peroxidases (PRXs) and laccases (LACs) involved in the cell wall polymerization of lignin precursors (Table S9). As shown for *NbPRX5* (Niben101Scf03861g01001), *NbPRX6a* (Niben101Scf02349g03002), and *NbLAC14a* (Niben101Scf09649g06014), the upregulation of lignin polymerization genes was preferentially detected at late time points and was highly specific to CoVLP samples (Figure 5a). As the latter patterns of expression were clearly distinct from those of monolignol biosynthesis genes, these observations suggest monolignol biosynthesis and lignin polymerization genes to be regulated by distinct transcriptional networks and therefore to represent an early and a late response to S protein expression, respectively.

**Figure 5.**
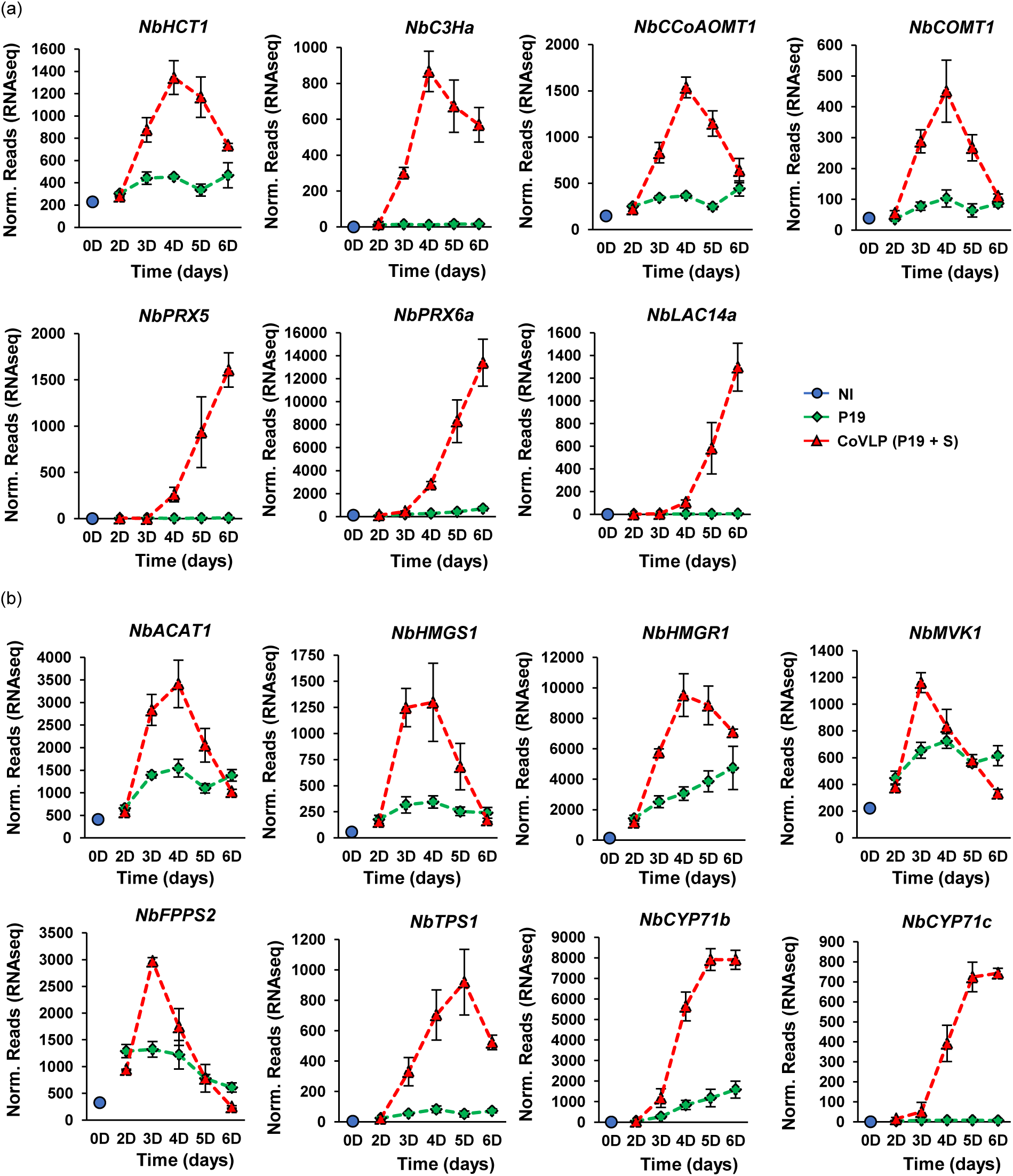
Upregulation of lignin- and terpene-related genes. (a) Expression profiles of genes involved in monolignol biosynthesis (upper panels), or cell wall lignin polymerization (lower panels). (b) Expression profiles of genes from the mevalonate pathway that promotes biosynthesis of terpene backbone molecules. As products of *TPS* and *CYP71* genes add structural diversity to terpene backbone molecules, expression profiles of *NbTPS1*, *NbCYP71b*, and *NbCYP71c* are also shown. For each time point in days (D) post-infiltration, RNAseq results are expressed in normalized (norm.) numbers of reads ± sd. Conditions are as follows: NI: non-infiltrated samples (blue); P19: agroinfiltrated samples expressing P19 only (green); CoVLP: agroinfiltrated samples co-expressing P19 and recombinant S protein (red).

### Upregulation of terpene-related genes

In plants, terpene metabolites play defensive roles, both as chemical deterrents for herbivores (Ninkuu *et al*., 2021) and as volatile organic compounds (VOCs) mediating inter-plant communications (Riedlmeier *et al*., 2017). We recently showed the expression of influenza HA to upregulate genes from the mevalonate pathway (Hamel *et al*., 2024a), one of the two terpene biosynthesis routes in plants (Ninkuu *et al*., 2021). Here, our RNAseq data revealed genes from the mevalonate pathway to be upregulated to some extent in P19 samples, but again the upregulation levels were much higher in CoVLP samples (Table S10). Generally, genes involved in the upstream steps of the mevalonate pathway were induced early after the onset of S protein accumulation, with maximal expression levels reached no later than 4 DPI. This pattern was for instance detected for acetoacetyl-CoA thiolase gene *NbACAT1* (Niben101Scf01100g01006), *3-hydroxy-3- methylglutaryl-CoA synthase* gene *NbHMGS1* (Niben101Scf01729g01015), *HMGR* gene *NbHMGR1* (Niben101Scf02203g05002), *mevalonate kinase* gene *NbMVK1* (Niben101Scf00370g03023), and *farnesyl pyrophosphate synthase* gene *NbFPPS2* (Niben101Scf00414g07005) (Figure 5b). The upregulation of genes involved in the downstream diversification of terpene molecules was also detected, including *terpene synthases* (*TPSs*) and *cytochromes P450 of family 71* (*CYP71s*) (Table S10). As depicted for *NbTPS1* (Niben101Scf00700g00005), *NbCYP71b* (Niben101Scf04869g01008), and *NbCYP71c* (Niben101Ctg08590g00001), upregulation of these genes in CoVLP samples was much stronger than in P19 samples, with maximum expression levels occurring later compared to those genes involved in the upstream steps of the mevalonate pathway (Figure 5b).

### Upregulation of genes related to oxidative stress and sugar metabolism

In plant cells, the production of ROS and consequent activation of the oxidative stress are among the most rapid and widespread responses following stress perception (Demidchik, 2015). To a major extent, ROS production is controlled by transmembrane NADPH oxidases, also known as respiratory burst oxidase homologs (RBOHs). In P19 samples and even more so in CoVLP samples, RNAseq revealed the upregulation of *NbRBOHd* (Niben101Scf02581g04013) and *NbRBOHf* (Niben101Scf02261g01002) (Table S11; Figure 6a), two close homologs of the defense-related genes *AtRBOHd* and *AtRBOHf* in Arabidopsis (Torres *et al*., 2002). During S protein expression, some of the most highly and specifically induced genes also encoded for other types of cellular oxidases that either directly or indirectly produce ROS. These included *NbPPO1* (Niben101Scf00180g08002) and *NbPPO3* (Niben101Scf04384g02014) (Table S11; Figure 6b), both of which encode polyphenol oxidases (PPOs). Typically activated in response to wounding and herbivory, PPOs oxidize phenolic compounds to generate reactive quinones that interact with oxygen and proteins to produce ROS (Boeckx *et al*., 2015). Also upregulated were genes encoding berberine-bridge enzymes (BBEs; Table S11), including highly expressed *NbBBE1b* (Niben101Scf01061g07011), *NbBBE2* (Niben101Scf00944g01001), and *NbBBE3a* (Niben101Scf01061g04027) (Figure 6c). BBEs belong to a functionally diverse family of proteins, but their catalytic mechanism always results in the production of hydrogen peroxide (Daniel *et al*., 2017). While these enzymes are involved in many cellular processes, the *BBE* genes here identified all encoded secreted proteins most similar to carbohydrate oxidases (Carter and Thornburg, 2004), thereby suggesting S protein expression to result in simple sugar oxidation in the apoplast, and in the consequent activation of oxidative stress in the extracellular space surrounding plant cells.

**Figure 6.**
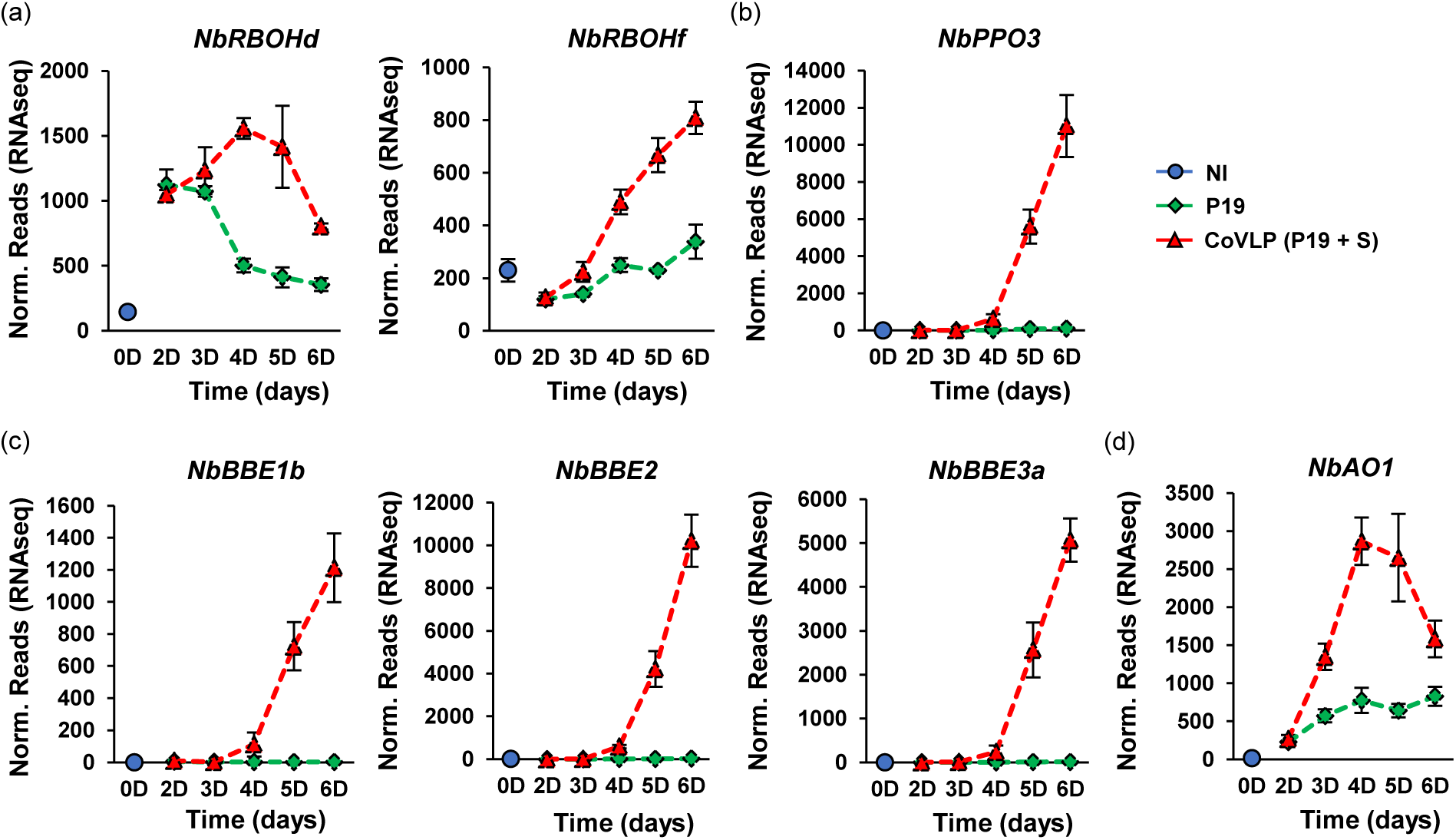
Upregulation of oxidative stress-related genes. Expression profiles of genes encoding NADPH oxidases (a), PPO (b), secreted carbohydrate oxidases of the BBE family (c), and secreted AO (d). For each time point in days (D) post-infiltration, RNAseq results are expressed in normalized (norm.) numbers of reads ± sd. Conditions are as follows: NI: non-infiltrated samples (blue); P19: agroinfiltrated samples expressing P19 only (green); CoVLP: agroinfiltrated samples co-expressing P19 and recombinant S protein (red).

Further evidence of modulated sugar signalling in the apoplast came from the altered expression of other key regulatory genes involved in this process (Table S11), including *cell wall invertase* (*CWI*) genes *NbCWI1* (Niben101Scf12270g02002) and *NbCWI2* (Niben101Scf04632g03006), *cell wall inhibitor of fructosidase* (*CIF*) gene *NbCIF1* (Niben101Scf01022g02008), and *sugar transport protein* (*STP*) genes *NbSTP1a* (Niben101Scf01529g02004), *NbSTP13a* (Niben101Scf05601g00010), *NbSTP13b* (Niben101Scf03380g03005), and *NbSTP14* (Niben101Scf01322g03005) (Figure S4). In Arabidopsis, the closest homologs of these genes were all found to be involved in immune responses (Lemonnier *et al*., 2014; Veillet *et al*., 2016). While *NbCWIs* and *NbSTP*s were all induced at much higher levels in CoVLP samples compared to P19 samples (Figure S4), *NbCIFs*, which encode protein inhibitors of CWI activity (Rausch and Greiner, 2004), were repressed in both P19 and CoVLP samples (Table S11; Figure S4). Altogether, these results suggest foreign protein expression to promote the conversion of sucrose to simple sugars in the apoplast, possibly to provide energy supporting local plant defense activation. In CoVLP leaves, the accumulation of simple sugars in the apoplast may however contribute to ROS production, considering that carbohydrate oxidases encoded by *BBE* genes are strongly induced in this condition (Figure 6c).

### Upregulation of apoplastic ascorbate oxidase genes

To prevent excessive self-damage caused by oxidative stress, plant cells produce a number of antioxidant compounds, including ascorbic acid (AsA) as the main antioxidant of the apoplast (Pignocchi *et al*., 2006). Accordingly, RNAseq revealed that the closely related *ascorbate oxidase* (*AO*) genes *NbAO1* (Niben101Scf03026g01009) and *NbAO2* (Niben101Scf22432g00001) were both induced in P19 and CoVLP samples, with much higher upregulation levels upon S protein expression (Tables S11; Figure 6d). Since secreted AOs negatively control the concentration and redox status of the AsA extracellular pool (Pignocchi and Foyer, 2003), this further pointed to an activation of oxidative stress in the apoplast, especially in S protein-expressing leaves. Overall, the observed regulation of oxidative stress-related genes in this study was strikingly similar to what was observed following the expression of influenza HA (Hamel *et al*., 2024a), a process also resulting in the production of enveloped VLPs in the apoplast.

### Regulation of SA response genes

To characterize stress hormone signalling in response to foreign protein expression, we investigated the expression of *Pathogenesis-Related 1* (*PR1*) genes. Similar to influenza HA expression (Hamel *et al*., 2024a), RNAseq revealed the upregulation of six *PR1* genes (Table S12) that could be separated in two clusters based on their distinct transcriptional behaviors. As shown for *NbPR1a* (Niben101Scf01999g07002) and *NbPR1b* (Niben101Scf00107g03008) (Figure 7a), genes from the first cluster were similarly induced, up to 4 DPI, in both P19 and CoVLP samples. At later time points, their expression kept increasing in P19 samples but started to decline in CoVLP samples. Notably, compromised induction of the *PR1* genes appeared to coincide with the high increase in S protein level (Figure 1d,e). As these *PR1* genes encode proteins that lack a C-terminal extension (CTE) (Hamel *et al*., 2024a), they are most similar to *AtPR1* (At2g14610), an Arabidopsis gene widely employed as a SA signalling marker (Durrant and Dong, 2004). Our results thus suggest that *Agrobacterium*-mediated expression of P19 in *N. benthamiana* leaves induces SA signalling and continued activation of SA-associated responses up to 6 DPI. In CoVLP samples, the situation was slightly more complex. At early time points, when no or little S protein was present, SA responses were similar to those observed in P19 samples. Past 4 DPI, when a significant amount of S protein had been produced, SA responses started to fade, most likely antagonized by S protein-induced signalling despite P19 co-expression and leaf infiltration with the *Agrobacterium* gene vector.

**Figure 7.**
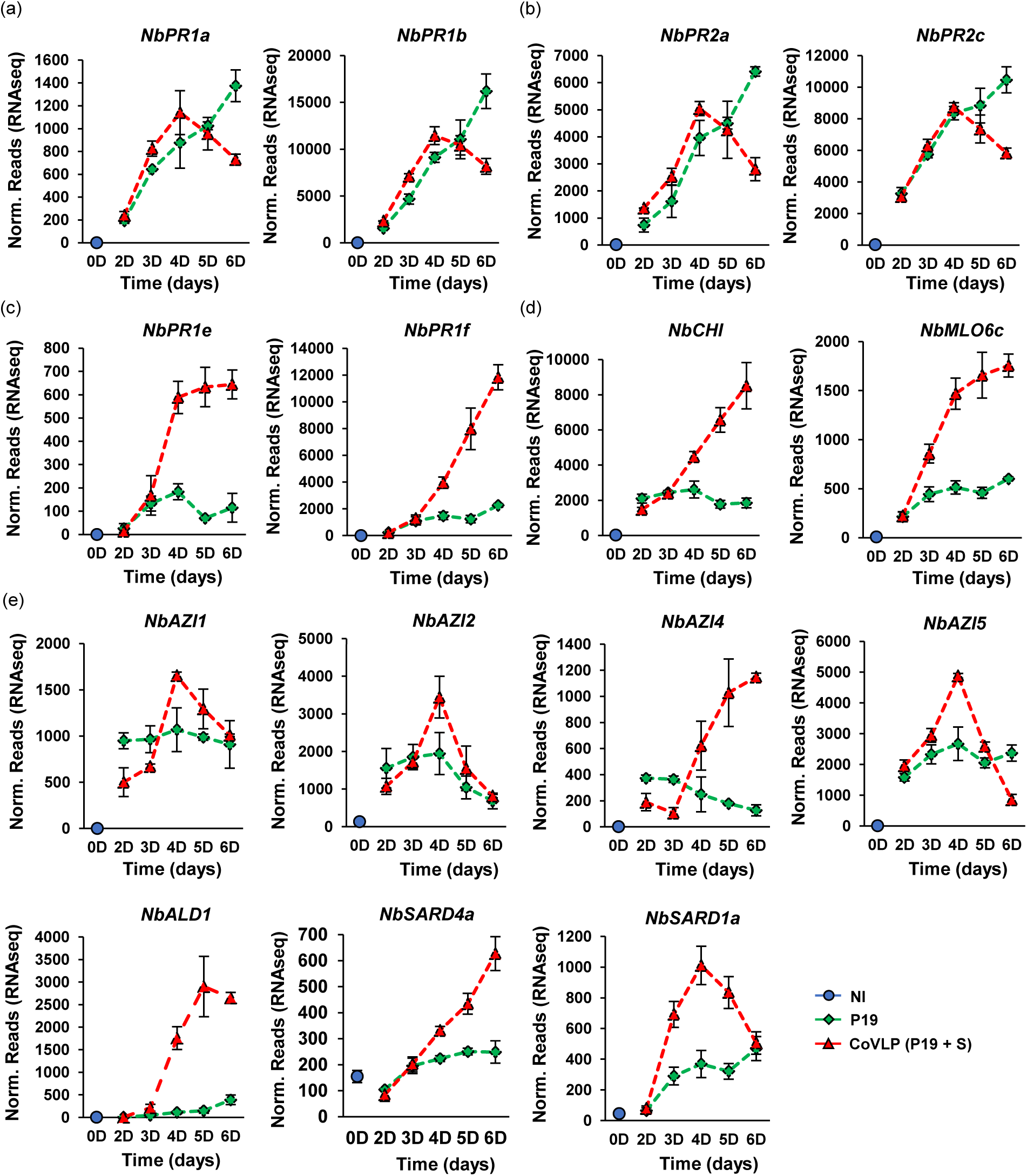
Expression of SA- and SAR-related genes. Expression profiles of SA signalling markers of the *PR1* (a) and *PR2* (b) gene families. Expression profiles of SAR-related genes is also shown, including SAR response markers of the *PR1* family (c), SAR response markers of other gene families (d), and SAR regulatory genes (e). For each time point in days (D) post-infiltration, RNAseq results are expressed in normalized (norm.) numbers of reads ± sd. Conditions are as follows: NI: non-infiltrated samples (blue); P19: agroinfiltrated samples expressing P19 only (green); CoVLP: agroinfiltrated samples co-expressing P19 and recombinant S protein (red).

Crucially, this particular expression pattern was observed not only for *PR1* genes, but also for *β-1,3-glucanase* genes of the *Pathogenesis-Related 2* (*PR2*) family (Table S12). Like *PR1* genes, *PR2* genes are often used as SA signalling markers (Durrant and Dong, 2004). As depicted for *NbPR2a* (Niben101Scf01001g00003) and *NbPR2c* (Niben101Ctg13736g00004), S protein expression resulted in compromised gene upregulation after 4 DPI, while transcript levels kept increasing when P19 only was expressed (Figure 7b). In CoVLP samples, compromised SA-mediated responses were also evidenced by the expression of genes encoding cytosolic heat shock proteins (HSPs), including small HSPs, HSP70s, and HSP90s (Table S13; Figure S5). Often associated to heat shock stress, enhanced *HSP* gene expression is also a well-known mark of SA signalling (Sangwan *et al*., 2022).

### Regulation of genes associated with systemic acquired resistance (SAR)

As introduced above, a second *PR1* gene cluster was differentially expressed during foreign protein expression (Table S12). Unlike the first cluster, genes from the second cluster, including *NbPR1e* (Niben101Scf04053g02007) and *NbPR1f* (Niben101Scf04053g02006), were similarly induced in P19 and CoVLP samples up to 3 DPI. Past that point, the upregulation levels became much higher when the S protein was expressed (Figure 7c). Due to their distinct transcriptional behavior and since they encode structurally distinct PR1 proteins harbouring a CTE, we previously proposed the expression of these genes to reflect SAR better than SA signalling in *N. benthamiana* leaves (Hamel *et al*., 2024a). Accordingly, the transcriptional profiles of *NbPR1e* and *NbPRf* matched those of several SAR response genes, including *NbCHI* (Niben101Scf02171g00007) and its close homolog *NbPR3a* (Niben101Scf07491g00003) (Table S12; Figure 7d). Encoding chitinases (CHIs) of the Pathogenesis-Related 3 (PR3) family, these genes are closely related to *AtCHI*, a well-established SAR marker in Arabidopsis (Riedlmeier *et al*., 2017). Also upregulated here were several homologs of the Arabidopsis gene *MILDEW RESISTANCE LOCUS O 6* (*AtMLO6*), another well-known marker of SAR (Table S12; Riedlmeier *et al*., 2017). Among *MLO6* genes here identified, *NbMLO6c* (Niben101Scf07792g02034) was the most strongly induced (Figure 7d). Together, these results suggest that SAR was induced in both P19 and CoVLP samples, albeit at a much higher level with the S protein expressed.

To further investigate SAR regulation upon recombinant protein expression, RNAseq data was searched for SAR regulatory genes (Table S12), including homologs of Arabidopsis *Azelaic Acid-Induced 1* (*AtAZI1*) (Cecchini *et al*., 2015), *AGD2-Like Defense 1* (*AtALD1*) (Song *et al*., 2004), *SAR-DEFICIENT 4* (*AtSARD4*) (Ding *et al*., 2016), and *AtSARD1* (Zhang *et al*., 2010). In all cases *N. benthamiana* homologs were induced to some extent in P19 samples, but with less intensity than in CoVLP samples (Figure 7e). In several cases, such as *NbAZI1* (Niben101Scf13429g02004), *NbAZI2* (Niben101Scf04779g01029), *NbAZI5* (Niben101Scf13429g03011), and *NbSARD1a* (Niben101Scf00428g09005), expression in CoVLP samples peaked at 4 DPI before going down at later time points. This indicated that upregulation of several SAR regulatory genes (Figure 7e) preceded the upregulation of SAR response genes (Figure 7c,d). Despite the late dampening of SA responses (Figure 7a,b; Figure S5), the induction of SAR-related genes in CoVLP samples suggests this pathway to be part of the host plant response to S protein expression, as also observed for the expression of influenza HA (Hamel *et al*., 2024a).

### Upregulation of oxylipin response genes

In many plant species, including Solanaceae, antagonism between the SA and JA pathways is well-established (Thaler *et al*., 2012). Based on this, the dampening of SA responses in CoVLP samples (Figure 7a,b; Figure S5) suggested such signalling crosstalk upon S protein accumulation. To further investigate this, we profiled the expression of genes commonly induced by JA or other oxylipins, including wounding- and herbivory-responsive genes. In addition to the previously described *PPO* genes (Table S11; Figure 6b), RNAseq revealed the upregulation of genes such as *plant defensins* (*PDFs*), *serine protease inhibitors* (*PIs*), *KTIs*, and *cysteine PIs* of the *Pathogenesis-Related 4* (*PR4*) family (annotated as *ʺwound-induced proteins*ʺ) upon S protein expression (Table S14). As depicted for *NbPDF1* (Niben101Scf17290g01005) and *NbPDF3* (Niben101Scf16258g02004), the upregulation *PDF* genes was strong and mostly specific to CoVLP samples (Figure 8a). For PI-encoding genes, certain candidates were also highly and specifically induced in CoVLP samples (e.g. *NbPI1;* Niben101Scf00294g00014), while others were also upregulated in P19 samples, albeit with lower intensity (e.g. *NbPR4c*; Niben101Scf01015g01002) (Figure 8b).

**Figure 8.**
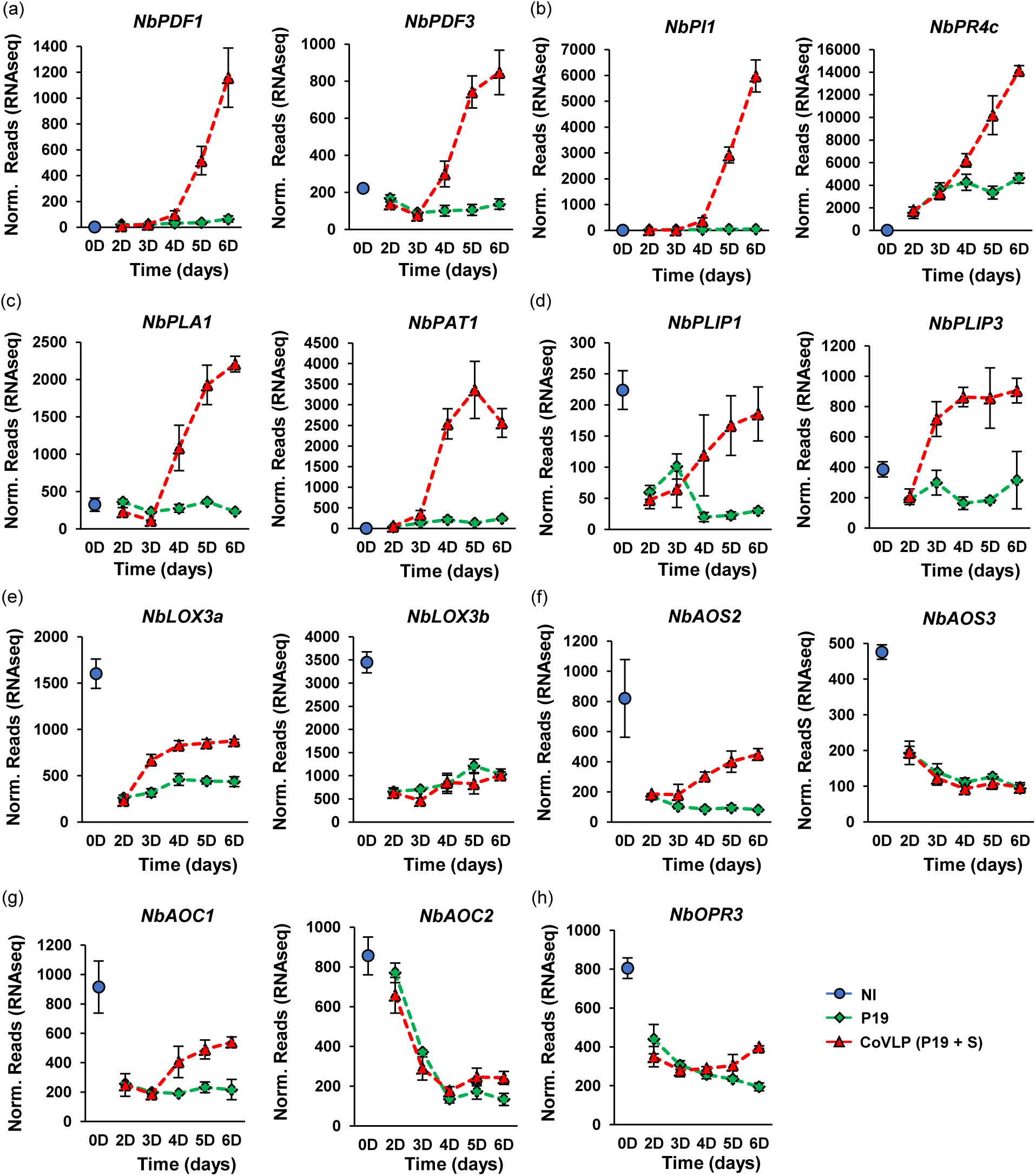
Upregulation of oxylipin response genes and expression of genes involved in JA biosynthesis. Expression profiles of oxylipin-responsive genes encoding PDFs (a) and PIs (b). Expression profiles of oxylipin regulatory genes are also shown. *PLAs* (c) and *PLIPs* (d) are involved in early steps of oxylipin biosynthesis, while *13-LOXs* (e), *AOSs* (f), *AOCs* (g), and *OPR3* (h) are more specific to JA biosynthesis. For each time point in days (D) post-infiltration, RNAseq results are expressed in normalized (norm.) numbers of reads ± sd. Conditions are as follows: NI: non-infiltrated samples (blue); P19: agroinfiltrated samples expressing P19 only (green); CoVLP: agroinfiltrated samples co-expressing P19 and recombinant S protein (red).

### Regulation of genes related to JA biosynthesis and signalling

The upregulation of genes such as *PPOs, PDFs*, and *PIs* suggested the prevalence of oxylipin signalling upon S protein accumulation, with JA presumably coordinating the observed responses. In plants, the production of JA requires the activation of phospholipases A (PLAs) and the channelling of resulting lipid intermediates through a specific branch of the oxylipin pathway (Wasternack and Feussner, 2018). Accordingly, RNAseq revealed the upregulation of several *PLA* genes (Table S14), including closely related *NbPLA1* (Niben101Scf03309g01011) and *NbPLA2* (Niben101Scf01228g02012). Also induced were several *patatin* (*PAT*) genes encoding enzymes closely related to tobacco homologs that promote the accumulation of oxylipins (Cacas *et al*., 2005). As shown for *NbPLA1* and *NbPAT1* (Niben101Scf06277g00007), enhanced *PLA* gene expression was strong and mostly specific to CoVLP samples (Figure 8c). Closely related genes *NbPLIP1* (Niben101Scf03952g00003), *NbPLIP2* (Niben101Scf01076g05026), and *NbPLIP3* (Niben101Scf01623g13002), which encode homologs of Arabidopsis PLASTID LIPASEs (AtPLIPs), were also differentially expressed (Table S14). Localized in chloroplasts, PLIPs also promote the accumulation of bioactive oxylipins, including JA (Wang *et al*., 2018). As shown for *NbPLIP1* and *NbPLIP3*, these genes were initially repressed in P19 and CoVLP samples, but specifically reinduced after 2 DPI in CoVLP samples (Figure 8d).

To produce JA, free fatty acids derived from the activity of PLAs are oxygenated by 13-lipoxygenases (13-LOXs), before their cyclization by allene oxide synthases (AOSs) and allene oxide cyclases (AOCs). The resulting 12-oxophytodienoic acid (OPDA) translocates to peroxisomes, where OPDA reductases (OPRs) and β- oxidation cycles complete the synthesis of JA (Wasternack and Feussner, 2018). In P19 and CoVLP samples, most of the JA biosynthesis genes identified were quickly repressed following agroinfiltration (Table S14). For instance, this was observed for *13-LOX* genes *NbLOX3a* (Niben101Scf02749g01001) and *NbLOX3b* (Niben101Scf07767g02004) (Figure 8e), for *AOS* genes *NbAOS2* (Niben101Scf10535g00001) and *NbAOS3* (Niben101Scf07309g00004) (Figure 8f), for *AOC* genes *NbAOC1* (Niben101Scf13816g00005) and *NbAOC2* (Niben101Scf02772g02001) (Figure 8g), and for the *OPR* gene *NbOPR3* (Niben101Scf05804g03006) (Figure 8h). As for *PLIPs* (Figure 8d), the expression of some JA biosynthesis genes was however reinduced as accumulation of the S protein started to increase (e.g. *NbLOX3a*, *NbAOS2*, and *NbAOC1*), suggesting that JA biosynthesis was repressed following *Agrobacterium* infiltration and/or expression of P19 only, perhaps due to the early induction of SA signalling (Figure 7a,b; Figure S5). As the S protein started to accumulate, the expression of some JA biosynthesis genes was however reinduced, indicating the JA pathway to be part of the host plant response to CoVLP expression.

Because JA needs to be conjugated to isoleucine (Ile) to become fully active (Staswick and Tiryaki, 2004), we also examined the expression patterns of genes involved in the conversion of JA to jasmonate-isoleucine (JA-Ile) (Table S14). In tobacco, the herbivore-induced production of JA-Ile requires products of the *Jasmonate-Resistant 4* (*JAR4*) and *JAR6* genes (Wang *et al*., 2008), a process also relying on threonine deaminases (TDs) for the biosynthesis of Ile (Kang *et al*., 2006). As here depicted for *NbTD1* (Niben101Scf02502g14001), *NbTD2* (Niben101Scf00682g04013), *NbJAR4a* (Niben101Scf05584g01007), and *NbJAR6a* (Niben101Scf00470g00001), the expression of all these genes was repressed shortly after agroinfiltration, and this phenomenon was observed in both P19 and CoVLP samples (Figure 9a,b). Based on this, we conclude that the conversion of JA to JA-Ile was not a prominent feature of the host plant response to S protein expression, similar to influenza HA expression (Hamel *et al*., 2024a). Since oxylipin-related responses were nonetheless strongly induced in CoVLP samples (Figure 6b; Figure 8a,b), we also conclude that oxylipin signalling not uniquely (or mainly) depended on the JA pathway and that other bioactive oxylipins were likely produced in leaf tissues under this condition. This possibility was further supported by the expression of close *AtMYC2* homologs *NbMYC2a* (Niben101Scf08672g02003), *NbMYC2b* (Niben101Scf06822g04004), and *NbMYC2c* (Niben101Scf15224g00002), which were all severely repressed in both P19 and CoVLP samples (Table S14; Figure 9c). As their Arabidopsis counterpart (Lorenzo *et al*., 2004), these TFs presumably act as master switches for the activation of JA signalling in *N. benthamiana*.

**Figure 9.**
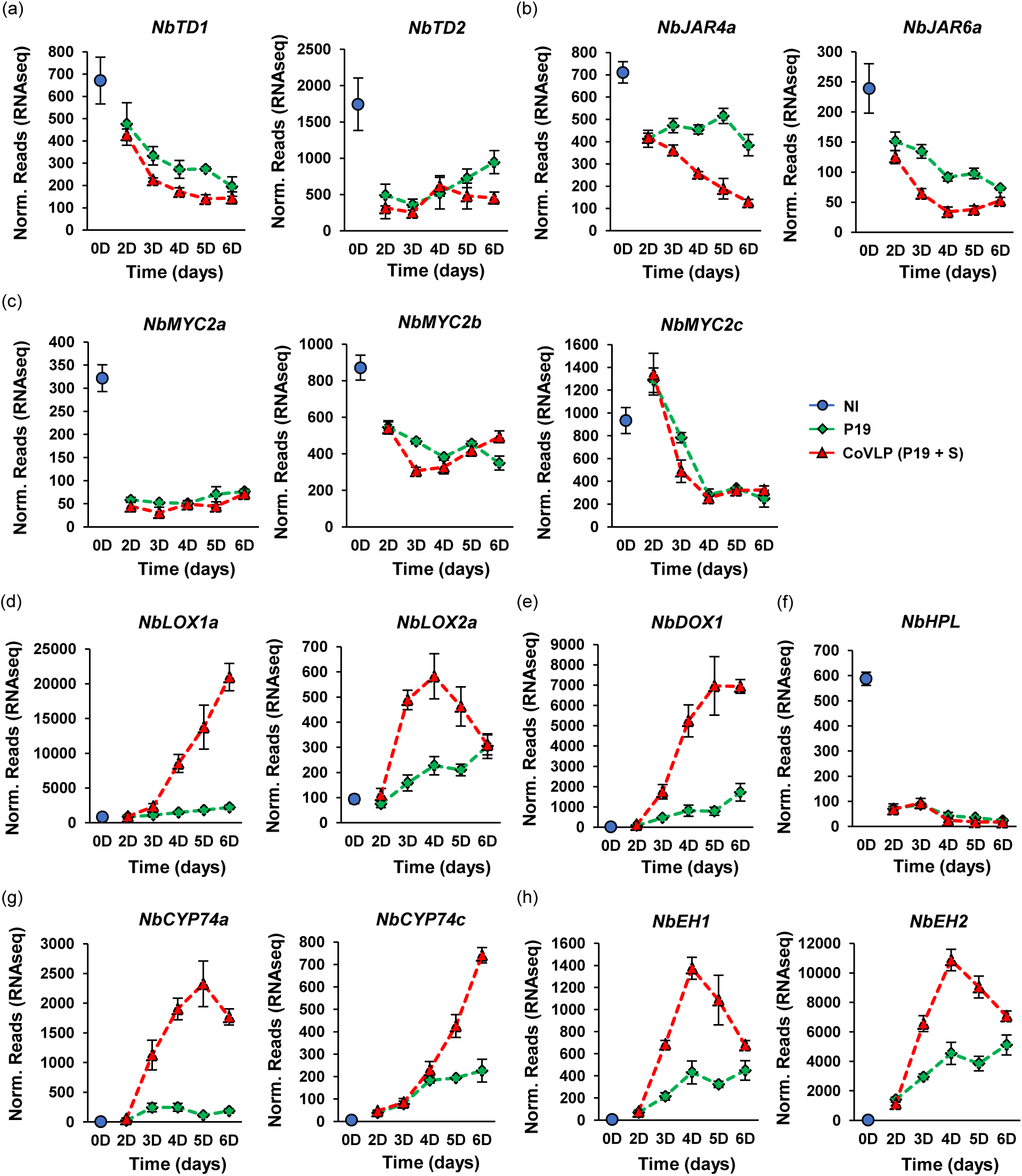
Expression of other oxylipin regulatory genes. Expression profiles of genes involved in Ile biosynthesis (a), or conjugation of Ile to JA (b). Expression profiles of *MYC2* genes, which encode TFs involved in JA signalling, are also shown (c), as well as those of genes involved in the biosynthesis of other bioactive oxylipins, namely *9-LOXs* (d), *DOX* (e), *HPL* (f), *CYP74s* (g), and *EHs* (h). For each time point in days (D) post-infiltration, RNAseq results are expressed in normalized (norm.) numbers of reads ± sd. Conditions are as follows: NI: non-infiltrated samples (blue); P19: agroinfiltrated samples expressing P19 only (green); CoVLP: agroinfiltrated samples co-expressing P19 and recombinant S protein (red).

### Expression of other oxylipin regulatory genes

Besides JA, the oxylipin pathway allows for the biosynthesis of other metabolites with roles in plant defense (Wasternack and Feussner, 2018). To shed light on which of these metabolites might be involved in S protein responses, RNAseq data was searched for more oxylipin regulatory genes. Several genes apparently involved in alternative branches of the oxylipin pathway were indeed differentially regulated (Table S14), including *9-lipoxygenase* (*9-LOX*) genes *NbLOX1a* (Niben101Scf01434g03006) and *NbLOX2a* (Niben101Scf01506g02001) (Figure 9d), as well as *α-dioxygenase* (*DOX*) gene *NbDOX1* (Niben101Scf04626g00009) (Figure 9e). These genes, induced to some extent in P19 samples, were however upregulated sooner and at much higher levels in CoVLP samples. In Arabidopsis, the closest homologs of *NbLOX1a* and *NbDOX1* encode enzymes that work in conjugation to promote the biosynthesis of oxylipins that activate local and systemic defenses (Vicente *et al*., 2012). 9-LOXs also work in combination with galactolipases of the PAT family to trigger PCD in tobacco (Cacas *et al*., 2005). The mining of our RNAseq data further identified *hydroperoxide lyase* (*HPL*) gene *NbHPL* (Niben101Scf00313g08016), which was repressed early in both P19 and CoVLP samples (Table S14; Figure 9f). In combination with 13-LOXs, HPL is a key enzyme for the production of green leaf volatiles (GLVs) (Huang *et al*., 2010). Here, *13-LOX* genes were also repressed after agroinfiltration, however *NbLOX3a* was reinduced upon S protein accumulation (Figure 8e). Reinduction of 13-LOX activity likely supported JA biosynthesis but not the production of GLVs, given the severe depletion of *HPL* transcripts after agroinfiltration (Figure 9f).

Our RNAseq dataset further revealed the upregulation of several *cytochrome P450 of family 74* (*CYP74*) genes (Table S14). Improperly annotated as *AOSs*, these genes actually encode divinyl ether synthases (DESs) or epoxyalcohol synthases (EASs) related to, but functionally distinct from, *bona fide* AOSs controlling JA biosynthesis. As depicted for *NbCYP74a* (Niben101Scf04787g02002) and *NbCYP74c* (Niben101Scf01574g01003), these genes were induced in P19 samples and even more in CoVLP samples (Figure 9g). In tobacco, NtDES1 coordinates with 9-LOXs to produce divenyl ether fatty acids involved in defense (Fammartino *et al*., 2007). As for EASs, these enzymes produce reactive epoxide compounds rapidly detoxified by epoxide hydrolases (EHs). The resulting oxylipin- diols have been suggested to have signalling as well as anti-microbial functions (Morisseau, 2013). As revealed by RNAseq (Table S14), *EH* genes *NbEH1* (Niben101Scf05133g06002) and *NbEH2* (Niben101Scf00640g04023) were also induced in both P19 and CoVLP samples, again at much higher levels upon S protein expression (Figure 9h).

In Arabidopsis and *Solanum lycopersicum* (tomato), OPDA is another oxylipin exhibiting JA-independent signalling functions (Stintzi *et al*., 2001; Bosch *et al*., 2014). Accordingly, OPDA-specific genes have been identified in Arabidopsis (Taki *et al*., 2005; Ribot *et al*., 2008). In CoVLP samples, but not P19 samples, RNAseq revealed a strong upregulation of several genes homologous to OPDA-specific genes in Arabidopsis (Table S14). These included the previously defined *BBE* genes (Figure 6c), *ribonuclease* genes *NbRNS1* (Niben101Scf04082g01004) and *NbRNS2* (Niben101Scf09597g01002), *phosphate transporter* gene *NbPHO1;H10* (Niben101Scf05712g02003), and *zinc finger protein* (*ZFP*) genes *NbZFP1* (Niben101Scf05373g01004) and *NbZFP2* (Niben101Scf10015g02003) (Figure S6). These results, which strongly suggest OPDA to be actively signalling in response to S protein expression, are consistent with the idea that oxylipin metabolites other than JA trigger plant immune responses in this condition.

### Large-scale proteomics data correlates with transcriptomics

Protein extracts from leaves P9 and P10 were prepared to investigate changes in protein abundance between P19 and CoVLP samples at 6 DPI. A large-scale proteomics survey was then conducted using isobaric tags for relative and absolute quantitation (iTRAQ) labelling. For both conditions, a total of 6,930 unique proteins were identified (Figure S7). Using P19 samples as control, pairwise comparisons were performed with CoVLP samples, so proteins specifically regulated by S protein expression could be identified. Differentially expressed proteins were defined as all protein candidates whose differential expression padj was below 0.05, corresponding to a fdr below 5%. At this expression threshold, 416 proteins were more abundant, and 418 proteins less abundant in CoVLP samples compared to P19 samples (Figure S7). Unsorted lists of up- and downregulated proteins are available in the Table S15.

Generally, proteomics data correlated with the transcriptional changes described above. For instance, CoVLP samples were found to be enriched in many UPR proteins, namely nine PDIs, six BiPs, three CRTs, and four CNXs (Table S15). This matched the expression patterns of the corresponding genes, which were induced in an early and transient fashion upon S protein expression (Table S7; Figure 3b). Likewise, proteomics revealed an enhanced accumulation of lignification-associated proteins in CoVLP samples, including six secreted PRXs (Table S15) whose corresponding genes were strongly and specifically induced in CoVLP samples (Table S9; Figure 5a). Enzymes involved in monolignol biosynthesis were also identified (Table S15), including a pair of phenylalanine ammonia lyases (PALs) and four cinnamate-4-hydroxylases (C4Hs). Not flagged by the RNAseq screening, the corresponding *PAL* genes were named *NbPAL1* (Niben101Scf05617g00005) and *NbPAL2* (Niben101Scf05442g03015), while the *C4H* genes were named *NbC4H1* (Niben101Scf02316g01002), *NbC4H2* (Niben101Scf00262g05006), *NbC4H3* (Niben101Scf08196g01007), and *NbC4H4* (Niben101Scf01085g04007). Proteomics data further revealed increased levels of NbCCoAOMT2 and NbCCoAOMT3, as well as 4-coumarate-CoA ligase (4CL) Nb4CL1 (Niben101Scf03224g00012). Again, these results matched upregulation patterns of the corresponding genes in CoVLP samples compared to P19 samples (Table S9). While suggesting an enhanced accumulation of lignin in S-expressing leaf tissues, proteomics also supported the idea of increased terpene biosynthesis under this condition. For instance, several enzymes of the mevalonate pathway showed increased levels compared to P19 samples, including NbACAT1, NbHMGS1, and NbFPPS2 (Table S15). Also more abundant in CoVLP samples were six TPSs and NbCYP71b (Table S15), all believed to operate downstream of the mevalonate pathway. For each terpene biosynthesis enzyme identified, the corresponding gene had been flagged as being predominantly induced in CoVLP samples compared to P19 samples (Table S10; Figure 5b).

Inferred from the upregulation of *AO* and *BBE* genes (Table S11; Figure 6c,d), oxidative stress in the apoplast was proposed to be a prominent feature of the host plant response to S protein expression. Accordingly, proteomics revealed increased abundance of NbAO1 and NbAO2, as well as upregulated levels of the three putative carbohydrate oxidases NbBBE1b, NbBBE2, and NbBBE3a (Table S15). Proteomics further suggested an enhanced alteration of sugar metabolism in the apoplast, as shown by higher levels of NbCWI1, NbCWI2, and NbSTP13b in CoVLP samples (Table S15). As revealed by RNAseq, corresponding genes were accordingly induced at much higher levels in CoVLP samples compared to P19 samples (Table S11; Figure S4).

The mining of our proteomics dataset further revealed CoVLP samples to be enriched in PR proteins, including the SAR marker NbCHI and its close homolog NbPR3a (Table S15). Proteomics further confirmed enhanced abundance of NbPR1f and NbPR1g (Table S15), a pair of PR1 proteins harbouring a CTE (Hamel *et al*., 2024a). As expression profiles of the corresponding genes matched protein accumulation patterns (Table S12; Figure 7c,d), proteomics data further highlighted the strong activation of SAR in CoVLP samples compared to P19 samples. These results were also consistent with the idea that PR1 proteins with a CTE better reflect SAR than SA signalling (Hamel *et al*., 2024a).

Compared to P19 samples, many PIs were also found to be more abundant in CoVLP samples (Table S15). These included four KTIs, NbPI1, NbPR4a, and NbPR4c, in line with specific or higher induction profiles of the corresponding genes in CoVLP samples (Table S14; Figure 8b). During S protein expression, strong oxylipin signalling was also evidenced by an increased abundance of oxylipin biosynthesis enzymes such as NbPAT1, NbLOX1a, NbLOX1b, NbDOX1, NbCYP74a, NbCYP74c, NbEH1, and NbEH2. Again, this matched a stronger induction of the corresponding genes in CoVLP samples (Table S14; Figure 8c, Figure 9). Overall, our proteomics data largely mirrored alterations of the leaf transcriptomes, further suggesting oxylipins other than JA to be produced at higher levels upon S protein expression.

As for the downregulated proteins, the most noticeable finding was that many HSPs were less abundant in CoVLP samples than in P19 samples (Table S15). Again, this correlated with expression profiles of the corresponding *HSP* genes, which were highly induced in P19 samples, but not in CoVLP samples (Table S13; Figure S5). As enhanced HSP expression reflects SA signalling, lower HSP levels in CoVLP samples supported the idea that S protein expression in leaf tissues causes the dampening of SA responses otherwise induced by agroinfiltration and P19 expression.

### Influenza VLP- and CoVLP-induced responses broadly overlap

Using biomass of leaves P9 and P10 harvested at 6 DPI, we previously studied molecular responses induced by the expression of H5, a recombinant version of influenza HA that leads to the formation of enveloped VLPs *in planta* (Hamel *et al*., 2024a). While H5 and the S protein display very limited homology, processes mediating influenza VLP and CoVLP formation share many features, including protein secretion through the plant cell endomembrane system and the budding of enveloped nanoparticles from the PM. In defining molecular changes induced by S protein expression, it became obvious that plant responses to influenza VLPs and CoVLPs were highly similar, with both types of enveloped nanoparticles not only leading to the altered expression of similar gene families, but also to the altered expression of the same genes within these families.

To better portray the extent of the overlap between influenza VLP- and CoVLP-induced responses, we used the transcriptome datasets of H5 and CoVLP leaf samples at 6 DPI, and performed pairwise comparisons with the P19 samples respectively used as controls for the expression of influenza VLPs (Hamel *et al*., 2024a) or CoVLPs (this study). As few of the downregulated responses were specific to the production of either nanoparticle type, we focused on the upregulated gene sets. Moreover, expression thresholds used for this comparison were set to match those previously used to study expression of the influenza VLPs (Hamel *et al*., 2024a) or the CoVLPs (this study) (Log_2_FC ≥ 2 and padj < 0.1). Compared to their respective P19 control, H5 samples displayed 925 upregulated genes, and CoVLP samples 1,213 upregulated genes (Figure 10a). Unsorted lists of upregulated genes are available in the Table S16. A Venn diagram inferred from the lists of upregulated genes then identified 361 genes specific to H5 samples, 649 genes specific to CoVLP samples, and 564 genes shared by the two conditions (Table S16; Figure 10a). In other words, ∼61% of the genes induced by H5 expression were also induced by the S protein, and ∼47% of the genes induced by S protein expression were also induced by H5. Among shared genes, those encoding late NAC and WRKY TFs were present, as well as those of proteins involved in activation of the oxidative stress, lignin polymerization, terpene biosynthesis, SAR responses, oxylipin synthesis, and oxylipin responses (Table S16; Figure 10a). Importantly, several genes scored as being specific to one or the other condition at 6 DPI were in fact induced in both conditions, but with different expression kinetics. For instance, *NbAO1* was identified as a H5-specific gene (Table S16) but it was clearly induced at earlier time points upon S protein expression (Figure 6d). Also critical was the fact that most genes with very high upregulation levels were shared by both conditions (Table S16). Despite major differences in size and sequence identity, H5 and the S protein thus induced gene sets that largely overlapped, possibly because similar disturbance effects on the endomembrane system were induced upon forced secretion of the complex viral proteins, or since similar alteration of the PM composition and architecture were induced upon budding of the nanoparticles.

**Figure 10.**
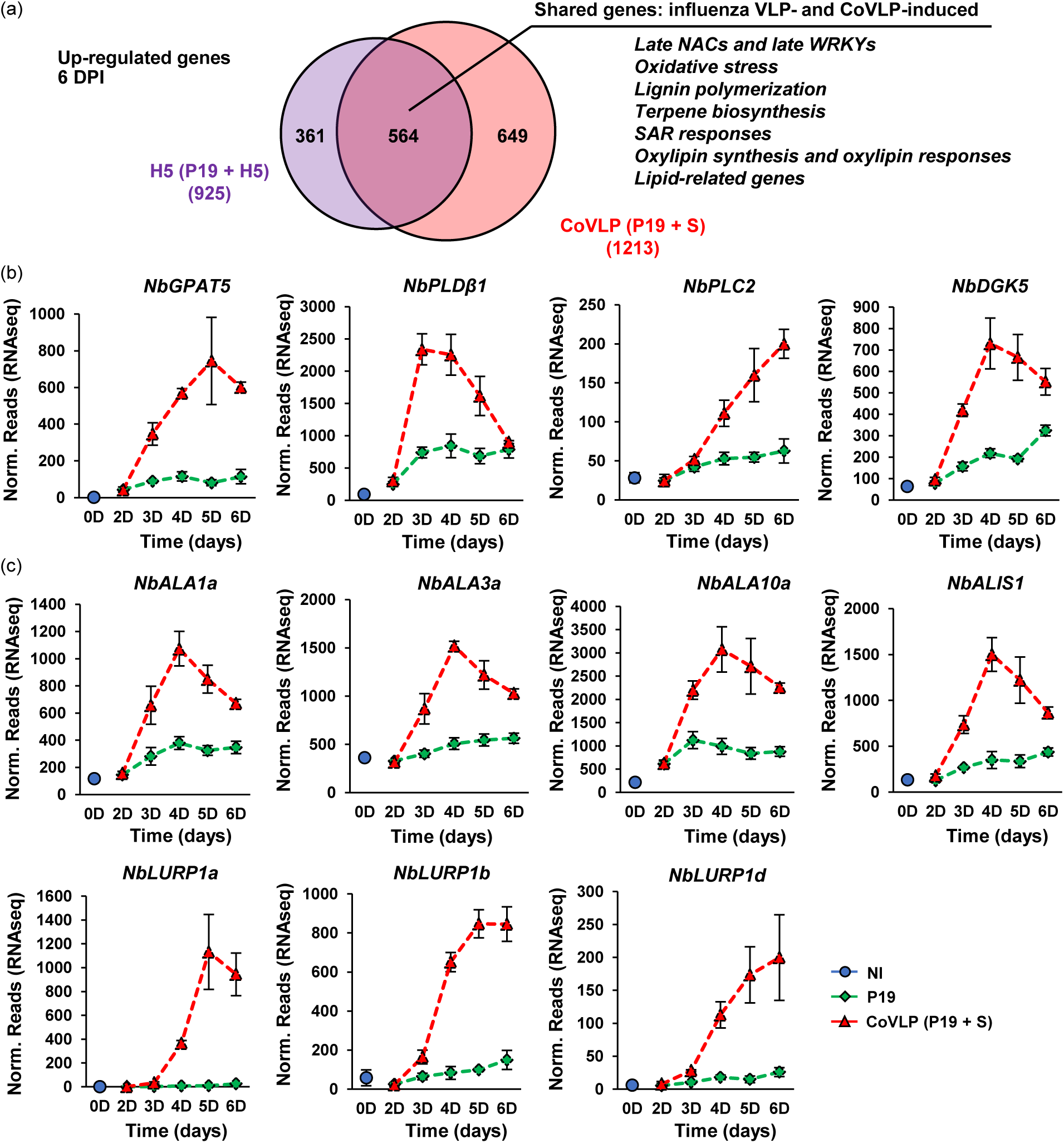
Overlap between enveloped nanoparticle responses and upregulation of lipid-related genes. (a) Venn diagram depicting the overlap between influenza VLP- and CoVLP-induced genes at 6 DPI. For H5 samples, genes were obtained from pairwise comparison with P19 samples from a previous study (Hamel *et al*., 2024a). For CoVLP samples, genes were obtained from pairwise comparison with P19 samples from this study. Expression thresholds are as follows: Log_2_FC ≥ 2 and adjusted p-value (padj) < 0.1. Circle size is proportional to the number of upregulated genes. Genes specific to H5 samples are shown in purple, those specific to CoVLP samples in red. Diagram intersect shows genes common to both conditions, with shared pathways highlighted on the right. Total numbers of upregulated genes are shown in parenthesis. Also displayed are the expression profiles of genes involved in lipid metabolism (b), or lipid distribution within membranes (c). For each time point in days (D) post-infiltration, RNAseq results are expressed in normalized (norm.) numbers of reads ± sd. Conditions are as follows: NI: non-infiltrated samples (blue); P19: agroinfiltrated samples expressing P19 only (green); CoVLP: agroinfiltrated samples co-expressing P19 and recombinant S protein (red).

To support the latter hypotheses, the expression profiles of genes involved in membrane lipid metabolism and distribution were examined upon CoVLP expression. Interestingly, many of those genes were more strongly induced in CoVLP samples than in P19 samples (Table S17), including the *glycerol-3- phosphate acyltransferase* (*GPAT*) gene *NbGPAT5* (Niben101Scf10067g02017), the *phospholipase D* (*PLD*) gene *NbPLDβ1* (Niben101Scf02465g00004), the *phospholipase C* (*PLC*) gene *NbPLC2* (Niben101Scf02221g00009), and the *diacylglycerol kinase* (*DGK*) gene *NbDGK5* (Niben101Scf06654g02004) (Figure 10b). While enhanced expression of these genes likely promotes defense signalling (Canonne *et al*., 2011), it could also lead to an increased production of phosphatidic acid, lysophosphatidic acid, and diacylglycerol. Interestingly, these lipids all exhibit conical shapes and their localized accumulation within membranes promotes bending rather than the formation of planar lipid bilayers (Roth, 2008). In line with this, genes involved in the redistribution of membrane lipids were also predominantly induced in CoVLP samples (Table S17), including homologs of Arabidopsis *aminophospholipid ATPase* (*ALA*) genes that encode lipid flippases (Figure 10c) (López-Marqués *et al*., 2014). Also induced in CoVLP samples was an *ALA-INTERACTING SUBUNIT* (*ALIS*) gene here termed *NbALIS1* (Niben101Scf02639g03015) as well as homologs of the Arabidopsis gene *LATE UPREGULATED IN RESPONSE TO HYALOPERONOSPORA PARASITICA 1* (*AtLURP1*), which most likely functions as a lipid scramblase (Figure 10c) (Bateman *et al*., 2009). Since these lipid-associated genes were also induced upon the expression of influenza VLPs (Table S16; Figure 10a), we previously hypothesized these transcriptional changes to represent feedback responses allowing proper modulation of membrane composition and architecture during the secretion of HA and/or the budding of influenza VLPs (Hamel *et al*., 2024a). Apparently, these changes are also necessary for S protein secretion and/or budding of the CoVLPs.

### Co-expression of KTIs reduces CoVLP-induced defense and leaf symptoms

Expression of CoVLPs triggered the upregulation of *PI* genes, including several *KTIs* (Table S14). A systematic search of the *N. benthamiana* genome indicated the KTI family in this plant to include 10 distinct members (Figure 11a), consistent with a previous survey of the PI repertoire in *N. benthamiana* (Grosse-Holz *et al*., 2018a). Our RNAseq data revealed eight of these 10 *NbKTI* genes to exhibit at least some expression in leaves P9 and P10 that were sampled. On the opposite, *NbKTI6* (Niben101Scf01971g01005) and *NbKTI9* (Niben101Scf02353g06034) showed no measurable expression in these specific organs (see asterisks in Figure 11a). Strongly and specifically induced in *N benthamiana* leaves expressing influenza VLPs (Hamel *et al*., 2024a), *NbKTI1* (Niben101Scf06424g02024), *NbKTI2* (Niben101Scf06825g00018), *NbKTI3* (Niben101Scf06424g00003), and *NbKTI4* (Niben101Scf06424g01007) were also strongly and specifically upregulated upon CoVLP expression (Figure 11b). By contrast, *NbKTI5* (Niben101Scf01971g01006) and *NbKTI7* (Niben101Scf08855g00008) exhibited basal expression in NI leaves prior to infiltration, but were both strongly repressed shortly after agroinfiltration, regardless of the foreign proteins expressed (Figure 11b). During the expression phase, these contrasted patterns among the *NbKTI* family members, confirmed by RTqPCR measurements (Figure S8a), suggested distinct roles for *NbKTI5* and *NbKTI7*. Interestingly, *NbKTI5* was previously proposed to be a close homolog of Arabidopsis *AtKTI1* (At1g73330) (Grosse-Holz *et al*., 2018b), a gene whose transcripts also get depleted upon *Agrobacterium* infection (Lee *et al*., 2009).

**Figure 11.**
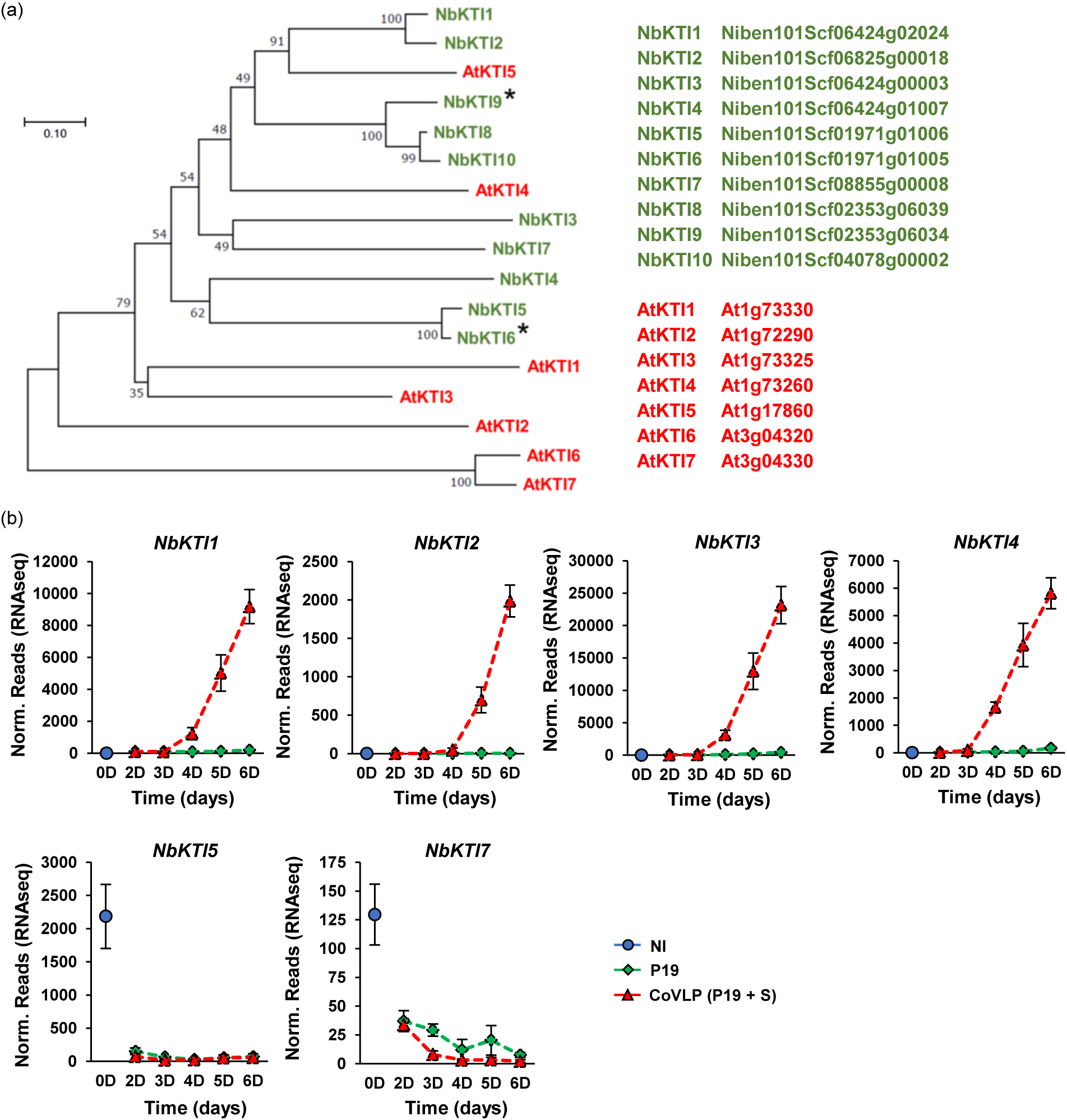
Phylogeny of KTIs and *NbKTI* gene expression. (a) The genome of *N. benthamiana* was searched using amino acid sequences of AtKTI1 and AtKTI5 as queries. Full-length sequences of retrieved proteins were aligned with ClustalW, using AtKTI6 and AtKTI7 as outgroups. The resulting alignment was submitted to the MEGA5 software, and a neighbor-joining tree derived from 5,000 replicates was generated. Bootstrap values are indicated on the node of each branch. Proteins of *A. thaliana* (At) are shown in red, those of *N. benthamiana* (Nb) in green. Identification numbers of corresponding genes are shown on the right. Asterisks (*) denote *NbKTIs* with no significant expression in P9 and P10 leaves harvested. (b) Expression profiles of up- and downregulated *NbKTIs*. For each time point in days (D) post-infiltration, RNAseq results are expressed in normalized (norm.) numbers of reads ± sd. Conditions are as follows: NI: non-infiltrated samples (blue); P19: agroinfiltrated samples expressing P19 only (green); CoVLP: agroinfiltrated samples co-expressing P19 and recombinant S protein (red).

A quick analysis of the NbKTI5 amino acid sequence indicated this protein to contain 239 amino acids, for a predicted molecular weight of 26.6 kDa and a calculated p*I* of 8.9. Sequence alignments revealed a conserved peptide, VLDINGEPLHVGEEYHI, characteristic of the PI family I03 (also known as the Kunitz family) (Figure S8b) (Park *et al*., 2000). *In silico* analysis suggested the nascent protein to include a 24-amino acids N-terminal SP for cellular secretion. The N-terminus of NbKTI5 also includes a pentapeptidic propeptide, NPIVL, presumably acting as a vacuolar sorting signal. Computer analysis finally revealed a putative NQT N-glycosylation site at position 141 (Figure S8b). These structural features are strictly conserved in close KTI homologs from other Solanaceae. Slightly smaller than NbKTI5, NbKTI7 includes 221 amino acids for a predicted molecular weight of 24.1 kDa and a calculated p*I* of 8.6. This protein also harbours a predicted N-terminal SP of 30 amino acids, the conserved peptide VLDIDGHELQIGSKYTI typical of Kunitz inhibitors, and three putative N-glycosylation sites at positions 60 (NGT), 106 (NLS), and 166 (NES). No vacuolar sorting signal was identified but the other features of NbKTI7 are strictly conserved in close KTI homologs from other Solanaceae (Figure S8b).

Because several proteases become activated during plant immunity and execution of PCD (Stael *et al*., 2019), we reasoned that the depletion of *NbKTI5* and *NbKTI7* transcripts could likely be involved in the activation of plant defense, and/or the onset of leaf necrosis during CoVLP expression. Such a scenario would be somewhat similar to AtKTI1 in Arabidopsis, a Kunitz inhibitor shown to antagonize PCD during immune responses (Li *et al*., 2008). To test this hypothesis, *NbKTI5* and *NbKTI7* were cloned and individually co-expressed with the CoVLP construct. After six days of expression, leaves P9 and P10 were selectively harvested, as for the RNAseq sampling. Compared to leaves agroinfiltrated with an empty vector (EV) control lacking a *KTI* transgene, leaves agroinfiltrated to co-express *NbKTI5* or *NbKTI7* showed a clearly reduced intensity of the CoVLP-induced leaf symptoms (Figure 12a). Beneficial effects on biomass quality not only consisted in obvious reduction of leaf necrosis, but also in reduced curling of leaf blade edges otherwise provoked by CoVLP expression.

**Figure 12.**
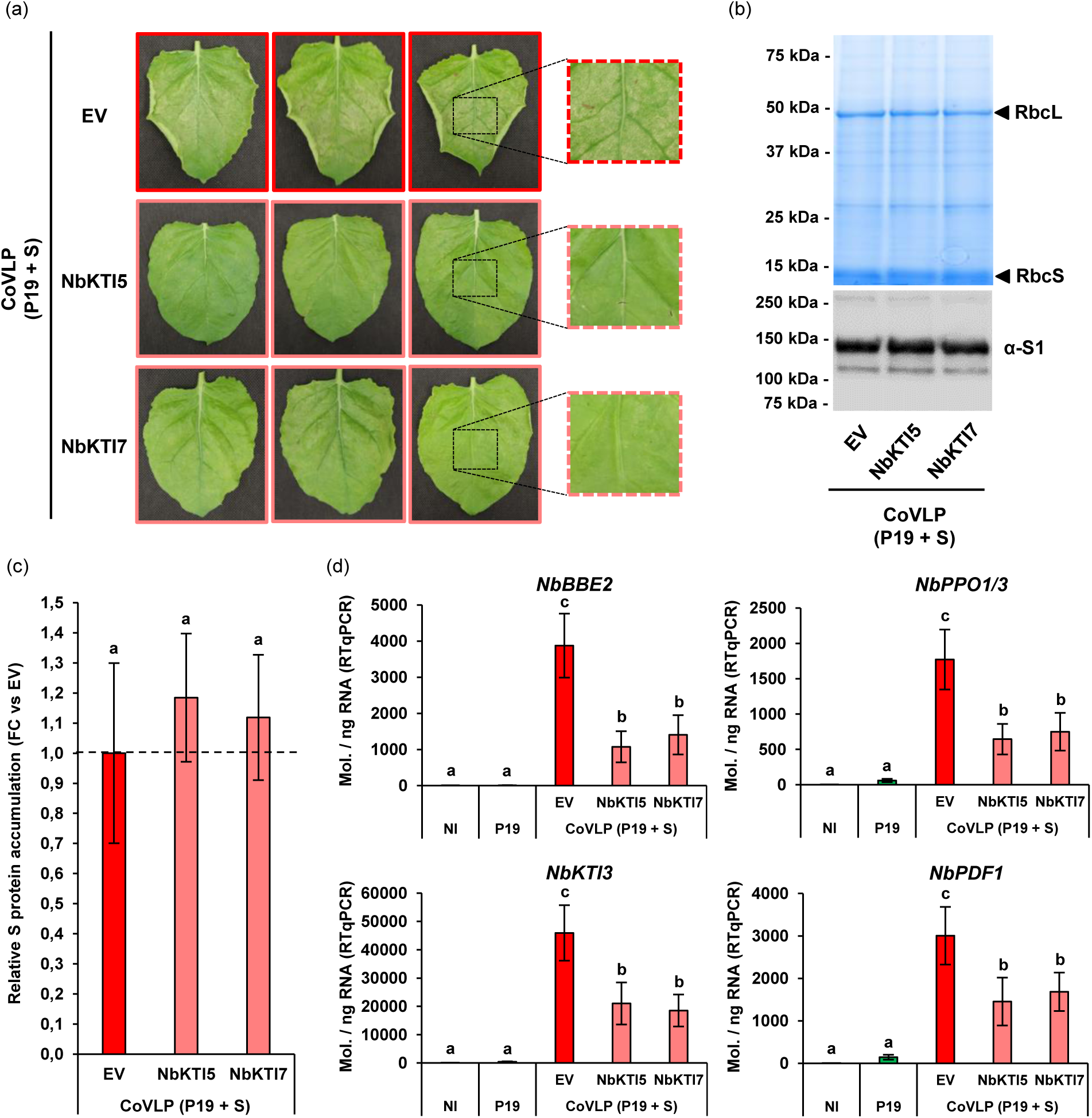
Co-expression of NbKTI5 or NbKTI7 reduces CoVLP-induced defense and leaf symptoms. Impact of helper protein candidates was assessed by co-expressing CoVLPs with either NbKTI5, NbKTI7, or an empty vector (EV) control. (a) Stress symptoms observed on representative leaves of each condition at 6 DPI. Based on primary leaf numbering (Figure S1a), pictures either highlight P9 or P10 leaves. On the right, magnified leaf sections highlight differences in leaf symptoms at the end of sampling. (b) Total protein extracts after SDS-PAGE and Coomassie blue-staining (upper panel). Arrows highlight RuBisCO small (RbsS) and large (RbsL) subunits. The western blot (lower panel) depicts S protein accumulation. (c) Relative S protein accumulation as measured by standardized capillary western immunoassays. S protein accumulation of CoVLP + EV samples was arbitrarily set at one-fold (dashed line). (d) Expression of CoVLP-induced genes *NbBBE2*, *NbPPO1/3*, *NbKTI3*, and *NbPDF1* as measured by RTqPCR. NI and P19 samples were used as controls. Results are expressed in numbers of transcript molecules per ng of RNA ± se. Groups that do not share the same letter are statistically different.

To ensure that reduced leaf symptoms were not simply associated with a lower accumulation of S protein in leaf tissues, P and S leaves from whole plant shoots were harvested altogether (Goulet *et al*., 2019), and their total soluble proteins extracted to monitor S protein levels in the presence or absence of KTIs. After immunoblotting, band signals showed S protein contents in KTI-expressing biomass to be roughly the same as in the EV control samples (Figure 12b). Using the same protein extracts, standardized capillary western immunoassays were conducted to formally quantify S protein levels, using EV control extracts as a baseline arbitrarily set at one-fold (see dashed line in Figure 12c). After six repetitions (n=6) and consistent with the immunoblot (Figure 12b), S protein levels were not significantly affected by co-expression of the KTIs (Figure 12c). This confirmed that NbKTI5 and NbKTI7, while improving the overall quality of the leaf biomass, did not interfere with S protein accumulation *in planta*.

To test whether immune signalling was also reduced upon KTI co-expression, biomass from leaves P9 and P10 collected to assess symptom intensity (Figure 12a) was used to perform RTqPCR analyses for genes shown previously to be specifically upregulated upon S protein expression. For *NbBBE2*, *NbPPO1/3*, *NbKTI3*, and *NbPDF1*, no or very limited expression was detected in NI and P19 samples (Figure 12d), confirming the CoVLP-specific induction of these genes in selected leaf tissues. Similar expression patterns were detected for the four genes in CoVLP samples, with significantly lower transcript abundance in the presence of NbKTI5 or NbKTI7 compared to the EV control (Figure 12d). Lower defense gene expression was consistent with lower leaf symptoms associated with the co-expression of either KTIs (Figure 12a).

## Discussion

In characterizing the effects of foreign protein expression in *N. benthamiana* leaf cells, we found a strong convergence in datasets gathered using RNAseq, RTqPCR, and iTRAQ proteomics. Overall, our results thus provide a comprehensive overview of the complex, sequential, and interconnected plant responses induced following agroinfiltration and subsequent accumulation of the viral foreign proteins. Together, these responses deeply affected the physiological status of plant cells and the overall fitness of host plants by activating a myriad of immune responses. Some responses were induced in both P19 and CoVLP samples, but with gene activation levels generally much higher upon S protein expression. On the opposite, some responses were more specific to a defined condition, including the enhanced expression of *HSP* genes and their corresponding proteins in P19 samples (Figure S5; Table S15), or the induction of lipid- and oxylipin-related responses in CoVLP samples (Figure 8; Figure 9; Figure 10).

Among the many pathways induced upon S protein expression, the UPR is of particular interest in the context of molecular farming. In fact, forced expression of the S protein was expected to trigger the UPR as this secreted protein requires complex domain folding and extensive post-translational modifications, including disulfide bridge formation and N-glycosylation. Here, S protein expression induced *NbbZIP60* (Figure 3a), which encodes a membrane-tethered TF acting as a master switch of the UPR. Expression of the S protein also led to the upregulation of several *NAC* genes that similarly encode membrane-tethered TFs, highly homologous to UPR-related NACs in Arabidopsis (Figure 4a). While these *NAC* genes remain to be formally characterized in *N. benthamiana*, our results strongly suggest that they participate in a complex transcriptional network operating downstream of *NbbZIP60*, similar to their closest homologs in Arabidopsis (Sun *et al*., 2013; Yang *et al*., 2014, 2023). During expression of the secreted S protein, we hypothesize the coordinated expression of these TFs to be required for optimal activation of the first UPR signalling branch.

In plants, the UPR comprises a second signalling branch, also controlled by membrane-tethered bZIPs (Howell, 2021). In *N. benthamiana*, these *bZIPs* were recently proposed to be *NbbZIP17* (Niben101Scf32851g00038), *NbbZIP28a* (Niben101Scf03647g01004), and *NbbZIP28b* (Niben101Scf00077g08013) (Hamel *et al*., 2024b). At the expression thresholds here employed, these TF genes were not flagged by RNAseq, but the mining of gathered data revealed these *bZIPs* to be more rapidly and more strongly induced in CoVLP samples compared to P19 samples (Figure S9). These findings thus suggest that both UPR signalling branches were activated upon S protein expression, leading to a robust upregulation of UPR genes that for instance encode ERQC components (Table S7; Figure 3b) involved in the folding and in the maturation of secreted proteins in the ER.

The early, but transient nature of UPR gene induction suggests that enhanced ERQC and ERAD functions were sufficient to cope with increased needs for protein secretion, or with ER stress provoked by agroinfiltration and forced expression of the S protein. Despite decreased UPR gene expression at late time points (Figure 3), proteomics revealed the levels of many UPR proteins to remain more abundant in CoVLP samples at 6 DPI (Table S15). Cellular benefits from early and transient UPR gene induction thus appears to be more sustained than what transcriptional profiles would suggest. We previously showed that the expression of influenza HA in *N. benthamiana* leaves also results in early and transient activation of the UPR (Hamel *et al*., 2024b). As for the HA protein, we here suggest that the activation of the UPR contributes to efficient folding and maturation of the S protein, which in turn contributes to the efficacy of COVID-19 vaccine production *in planta*.

While many lignin-related genes were induced upon S protein expression, monolignol biosynthesis and lignin polymerization genes showed distinct expression kinetics (Figure 5a). In Arabidopsis, the expression of monolignol biosynthesis genes was similarly shown to be uncoupled from the expression of lignin polymerization genes, with AtSND2 promoting expression of the latter group (Hussey *et al*., 2011). Here, S protein expression resulted in late upregulation of *NbSND2* and *NbSND3* (Figure 4b), two closely related *NACs* that potentially regulate the expression of lignin polymerization genes in *N. benthamiana*. In Arabidopsis, expression of monolignol biosynthesis genes relies on the other hand on MYB TFs, including AtMYB15 (Kim *et al*., 2020) and AtMYB43 (Geng *et al*., 2020). Consistent with early upregulation of monolignol biosynthesis genes upon S protein expression (Figure 5a), RNAseq here revealed an early upregulation of *MYBs* that encode close homologs of AtMYB15 and AtMYB43 (Table S8), including closely related *NbMYB14a* (Niben101Scf01013g01003), *NbMYB14b* (Niben101Scf03064g05001), and *NbMYB15a* (Niben101Scf06017g02002), as well as the more distant homolog *NbMYB43* (Niben101Scf02854g05001) (Figure S3b). In the context of molecular farming, defining transcriptional networks that control the expression of lignin-related genes is of great significance, since a reinforcement of the plant cell wall may impact downstream process efficacy, especially when the desired products are to be secreted.

In Arabidopsis, volatile terpene metabolites have been shown to enhance the expression of a specific subset of SAR-related genes (Riedlmeier *et al*., 2017). As this gene subset closely mirrors SAR-related genes induced upon influenza HA expression, we recently hypothesized that terpene signalling promotes SAR in this particular molecular farming context (Hamel *et al*., 2024a). If true, one would expect terpene biosynthesis genes to be induced early after agroinfiltration, so downstream upregulation of SAR-related genes can then intervene. While the use of a single sampling point at 6 DPI prevented further examination of this hypothesis during HA expression, time course sampling here confirmed the early upregulation of terpene biosynthesis genes in response to S protein expression (Figure 5b). While not formally confirming that terpene signalling indeed acts upstream of SAR during influenza VLP or CoVLP expression, our results are at least consistent with this hypothesis. In the context of molecular farming, plants are generally packed in closed incubation chambers after agroinfiltration and inter-plant communications would thus be expected to play a significant role in defense activation. This is especially relevant since the expression phase usually lasts for several days. To further challenge this model, characterization of VOC emissions during foreign protein expression will be necessary, and eventually highly meaningful as more knowledge on this question might result in improved molecular farming practices.

The activation of SAR is often linked to SA signalling (Durrant and Dong, 2004). When expressing influenza VLPs or CoVLPs, the positioning of terpene signalling upstream of SAR would also reconcile the fact that SAR-related genes were strongly induced despite the dampening of SA responses (Figure 7a,b; Figure S5) (Hamel *et al*., 2024a). Time course sampling indeed suggested that crosstalk between SA and oxylipin signalling is a major feature of these expression systems. Early repression of JA biosynthesis genes was detected in both P19 and CoVLP samples (Figure 8), a phenomenon likely triggered by SA signalling associated to agroinfiltration and P19 expression. As the S protein started to accumulate, the expression of some JA biosynthesis genes was reinduced, a process not observed in P19 samples (Figure 8). Moreover, S protein expression resulted in the enhanced expression of other oxylipin regulatory genes (Table S14; Figure 9), and in a strong upregulation of classical oxylipin response genes (Table S14; Figure 6b; Figure 8a). From these results, several conclusions can be drawn. First, JA biosynthesis genes were not all regulated in the same way, perhaps because distinct transcriptional networks controlled their expression. Second, CoVLP-induced signalling was strong enough to partially overcome early repression from one of the JA biosynthesis gene subsets. Third, S protein expression resulted in strong oxylipin signalling that apparently led to late antagonism of SA-dependant responses. Indeed, the induction (or the reinduction) of oxylipin-related genes coincided with the late dampening of SA responses, as revealed by the expression of *PR1* (Figure 7a), *PR2* (Figure 7b) and *HSP* (Figure S5) genes. These intricate signalling interplays highlight the plasticity of the plant immune system, which can be reoriented in a timely fashion according to the foreign proteins expressed and despite early signalling initiated after agroinfiltration.

In *N. benthamiana*, stress-induced production of SA relies on the enhanced expression of *isochorismate synthase* (*ICS*) genes (Catinot *et al*., 2008). Following agroinfiltration, RNAseq showed *NbICS1* (Niben101Scf00593g04010) to be rapidly repressed in both P19 and CoVLP samples (Tables S12; Figure S10a), suggesting this *ICS* gene not to be involved in the early activation of SA signalling. The genome of *N. benthamiana* comprises a second *ICS* gene, which was here termed *NbICS2* (Niben101Scf05166g06006). Although *NbICS2* was not flagged by RNAseq because it did not meet the required expression thresholds, this gene was robustly induced from 2 to 5 DPI in P19 samples (Figure S10a). At 2 DPI, *NbICS2* was also upregulated in CoVLP samples, although showing decreased expression as the S protein started to accumulate (Figure S10a). The transcriptional profile of *NbICS2* suggests this gene to be involved in the early activation of SA signalling and in associated repression of JA biosynthesis genes after agroinfiltration. Decreased expression of this gene in CoVLP samples, after an initial upregulation, also correlated with late antagonism of SA response. In light of these expression patterns, we propose that *NbICS2* is a key regulatory node controlling hormonal crosstalk during *Agrobacterium*-mediated expression of the foreign proteins here tested.

In Arabidopsis, decreased SA responses have been associated with *ICS* gene repression, but also with increased storage of SA following the upregulation of *SA GLUCOSYL TRANSFERASE 1* (*AtSAGT1*) (Zheng *et al*., 2012). This gene encodes an enzyme that converts SA into its inactive conjugated forms (Dean and Delaney, 2008). In *N. benthamiana*, the closest homolog of *AtSAGT1* is *NbSAGT1* (Niben101Scf05415g00003). RNAseq here showed *NbSAGT1* to be strongly induced in CoVLP samples (Figure S10b), matching our proteomics data showing the enhanced accumulation of the corresponding protein in CoVLP samples at 6 DPI (Table S15). Again, these results suggest S protein expression to dampen the SA pathway, possibly through the induction of late oxylipin signalling.

Much attention has been given to JA in plants responding to necrotrophic pathogens, wounding, or herbivory. While this compound appears to play a role in responses to influenza VLP (Hamel *et al*., 2024a) and CoVLP (Figure 8) expression, our data suggests the observed oxylipin responses to not solely (or mainly) depend on JA signalling. Other bioactive oxylipins likely produced during foreign protein expression are yet to be formally identified, but our RNAseq data provides circumstantial evidence for OPDA signalling in response to influenza VLP (Hamel *et al*., 2024a) and CoVLP (Figure S6) expression. The upregulation of other oxylipin regulatory genes further suggests oxidized polyunsaturated fatty acids, divinyl ethers, epoxy-oxylipins, and oxylipin-diols to be actively signalling in these conditions (Table S14; Figure 9) (Hamel *et al*., 2024a). Considering the high upregulation levels of oxylipin response genes, we hypothesize that a blend of oxylipin signals contribute to the activation of plant defenses and perhaps to the onset of PCD following influenza VLP (Hamel *et al*., 2024a) and CoVLP (Figure 1a) expression. To optimize production of plant-made vaccines against influenza and COVID-19, a formal identification of bioactive oxylipins will be instrumental considering that both types of nanoparticles induce similar oxylipin signalling and responses (Figure 10a).

Cellular processes that drive VLP and CoVLP assembly share many features, even though the recombinant proteins at the surface of these nanoparticles diverge considerably. In fact, the extensive overlap between influenza VLP- and CoVLP-induced responses suggests that the induced immune responses in leaf tissues are mostly due to the production of nanoparticles *per se*. At the cellular level, these effects can perhaps be triggered by the continuous ‘highjacking’ of PM lipids during nanoparticle budding, a process eventually perceived as ‘micro-wounds’ that would then induce oxylipin signalling. Activation of plant immunity might also result from a perturbation of the plant cell endomembrane system during forced secretion of the viral surface proteins. This scenario would be consistent with the upregulation of genes associated with lipid metabolism and distribution in cellular membranes (Figure 10b,c) (Hamel *et al*., 2024a). Despite many unsolved questions, extensive overlap observed in the host plant responses to influenza VLP and CoVLP expression could be an advantage from a practical standpoint, as strategies developed to improve the manufacturing of one product would likely benefit the manufacturing of the other product.

Here, a strategy based on the co-expression of NbKTIs helped to improve quality of the biomass expressing CoVLPs, without interfering with accumulation of the S protein (Figure 12). Additional work will be required to explain how these NbKTIs downregulate immune responses and PCD in CoVLP-expressing leaves, but the protease inhibitory spectra reported for close Solanaceae homologs (Rowan *et al*., 1990; Arnaiz *et al*., 2018) suggest that these KTIs behave as inhibitors of cysteine proteases (CPs) to maintain cellular homeostasis in the absence of stress. According to current models to explain the effects of PIs in plant cell environments (Benchabane *et al*., 2010), the transient co-expression of NbKTI5 (or NbKTI7) would compensate for the loss of endogenous *NbKTI* transcripts (Figure 11b) and help maintain an elevated PI/CP balance unfavorable to the onset of CP-inducible immunity and PCD-related responses. This model would be consistent with the upregulation of several *CP* genes, enhanced levels of the encoded CPs, and enhanced CP activity recently detected in CoVLP samples compared to NI or even P19 samples (Hamel *et al*., 2025). At 6 DPI, the beneficial effects of NbKTIs would also be consistent with the inhibition of CPs, activity of which largely overlaps with distribution of the S protein in young and actively growing leaf tissues (compare Figure S1b with Figure S11). While the functional characterization of NbKTI5 and NbKTI7 continues, it will be interesting to see whether these PIs also control plant responses leading to PCD, when observed upon the accumulation of other recombinant proteins commonly produced in plants, including influenza HA and therapeutic antibodies.

## Conclusion

Extensive datasets such as those generated in this study continue to expand our understanding of plant molecular responses to foreign protein expression in *N benthamiana*. While these responses likely vary according to the recombinant proteins expressed, the data provided is still highly meaningful as leaf agroinfiltration in *N. benthamiana* now represents a valuable and widely employed avenue to help solving global health issues such as the spreading of influenza- and COVID-19-causing viruses. Agroinfiltration undeniably represents an artificial pathosystem, in which a concentrated inoculum of root pathogen is forced into leaf tissues. In spite of the multiple stress layers imposed by this system, plants exhibit outstanding adaptability that yet allows them to efficiently produce a wide variety of recombinant products. By deciphering molecular responses associated with agroinfiltration and the subsequent accumulation of foreign proteins, our work will likely contribute to the definition of novel strategies aimed at improving the performance of molecular farming systems. Based on the data collected here, alternative improvement strategies we currently envision include the editing of the host plant genome and the heterologous co-expression of additional helper proteins to favor protein folding, cell detoxification or any other processes enhancing recombinant protein production in plants.

## Materials and methods

### Seed germination and plant growth

Seeds of wild-type *N. benthamiana* were spread on pre-wetted peat mix plugs (Ellepot) and placed in a germination chamber for 2 d, where conditions were as follows: 28°C/28°C day/night temperature, 16 h photoperiod, relative humidity of 90%, and light intensity of 7 µmol m^-2^ s^-1^. Germinated plantlets were then transferred in a growth chamber for 15 d, where conditions were as follows: mean temperature of 28°C over 24 h, 16 h photoperiod, mean relative humidity of 66% over 24 h, concentration of ambient carbon dioxide (CO_2_) of 800 ppm injected only during the photo-phase, and light intensity of 150 µmol m^-2^ s^-1^. Over this period, watering and fertilization were provided as needed. After 2 weeks, peat mix plugs were transferred into 4-inch pots containing pre-wetted peat-based soil mix (Agro- Mix). Freshly transferred plantlets were then moved to a greenhouse, where conditions were as follows: mean temperature of 25°C over 24 h, 16 h photoperiod, mean relative humidity of 66% over 24 h, 800 to 1,000 ppm ambient CO_2_ injected only during the photo-phase, and natural light conditions supplemented with artificial high pressure sodium lights for an overall light intensity of 160 µmol m^-2^ s^-1^. In the greenhouse, watering and fertilization were provided as needed. The plants were allowed to grow for ∼20 additional days, until ready for agroinfiltration.

### Binary vector constructs

To express CoVLPs, sequence of the full-length SARS-CoV-2 S protein was retrieved (strain hCoV-19/USA/CA2/2020). The isolated gene sequence consisted of nucleotides 21,563 to 25,384 from EPI_ISL_406036 in the GISAID database (https://www.gisaid.org/). Using PCR-based methods, the native S protein was modified to increase its stability by introducing three points mutations at the S1/S2 protease cleavage site (R667G, R668S, and R670S). The single substitutions K971P and V972P were also introduced to stabilize the S protein in its pre-fusion conformation (Wrapp *et al*., 2020). To favor protein entry in the secretory pathway of plant cells, the native SP of the S protein was replaced by the SP of a *Medicago sativa* (alfalfa) PDI. To optimize CoVLP assembly and budding, TMD and cytosolic tail (CT) of the native S protein were replaced by those of the HA protein of pandemic influenza virus strain H5 Indonesia (H5/A/Indonesia/05/2005) (D’Aoust *et al*., 2008). Once assembled, the chimeric *S* gene was reamplified by PCR and introduced, using the In-Fusion cloning system (Clontech), in the T-DNA region of a customized pCAMBIA0380 binary vector previously linearized with restriction enzymes *SacII* and *StuI*. Expression of the *S* transgene was driven by a *2X35S* promoter from the cauliflower mosaic virus (CaMV). The expression cassette also comprised proprietary 5’- and 3’-untranslated regions (UTRs) developed to maximize mRNA stability and protein translation, as well as the *Agrobacterium nopaline synthase* (*NOS*) gene terminator. To prevent silencing of transgene expression, T-DNA region of the CoVLP construct also comprised the tomato bushy stunt virus (TBSV) suppressor of RNA silencing gene *P19*, under the control of *plastocyanin* promoter and terminator sequences. For P19 samples, a binary vector allowing expression of the P19 silencing suppressor only was used.

For gene constructs allowing co-expression of NbKTI5 and NbKTI7, full-length nucleotide sequences of the corresponding *NbKTI* genes were retrieved from the *N. benthamiana* genome assembly database (https://solgenomics.net/). Wild-type sequences were codon-optimized and the resulting templates used to design synthetic gene blocks (gBlocks) (Integrated DNA Technologies). gBlocks were amplified by PCR and as described above, their amplicons individually introduced in the T-DNA region of a customized pCAMBIA0380 binary vector previously linearized with *SacII* and *StuI*. KTI expression was driven by a *2X35S* promoter from the CaMV. The expression cassette also comprised proprietary 5’- and 3’- UTRs, as well as the *NOS* terminator. An empty binary vector was used as control for the co-expression experiments.

### Agrobacterium cultures and leaf infiltration

Binary vectors were transformed by heat shock in *A. tumefaciens* strain AGL1. The transformed bacteria were plated on Luria-Bertani (LB) medium, with the transformation selection marker kanamycin at 50 µg/ml. Transformed colonies were allowed to grow for 2 d at 28°C. Frozen glycerol stocks were prepared from isolated colonies and kept at -80°C for long term storage. When ready, frozen bacterial stocks were thawed at room temperature before transfer in pre-culture shake flasks containing LB medium with kanamycin (50 µg/ml). Bacterial pre-cultures were grown for 18 h at 28°C with shaking at 200 rpm. Pre-cultures were next transferred to larger shake flasks containing kanamycin selection medium and the bacteria were allowed to grow for an extra 18 h at 28°C with shaking at 200 rpm. Using a spectrophotometer (Implen), bacterial inoculums were prepared by diluting appropriate volumes of the bacterial cell cultures in resuspension buffer (5 mM MES, pH 5.6, 10 mM MgCl_2_). For CoVLP expression, a final OD_600_ of 0.6 was used. For co-expression of helper proteins, separate bacterial cell cultures were prepared as described above. Prior to infiltration, appropriate volumes of the CoVLP and helper protein bacterial cell cultures were mixed to reach final OD_600_ of 0.6 and 0.2, respectively. Vacuum infiltration was performed by placing whole plant shoots upside down in an airtight stainless-steel tank containing the appropriate bacterial suspension. To draw air out of the leaves, vacuum pressure was applied for 1 min, before pressure release to force the bacterial inoculum into the leaves.

### Transient protein expression and biomass harvesting

Recombinant protein accumulation was allowed to proceed in agroinfiltrated leaves for several days, as indicated. For all experiments, the expression phase took place in plant growth chambers, where conditions were as follows: 20°C/20°C day/night temperature, 16 h photoperiod, relative humidity of 80%, and light intensity of 150 µmol m^-2^ s^-1^. Watering was performed every other day, with no fertilizer supplied during expression. For the sampling of leaf sections used to map S protein distribution and CP activity, P leaves and S leaf bundles were separated manually. Starting from the bottom of plants, P leaves were numbered based on their respective position along the main stem (P1 to P12). Primary leaf numbering excluded cotyledons, as well as small and not fully expanded leaves located close to the apex. Based on this numbering, a first set of four P leaf sections was created by pooling pairs of adjacent leaves (P12 + P11, P10 + P9, and so on). For the fifth P leaf section, leaves P4, P3, P2, and P1 were harvested together because P1 and P2 leaves were very small. Similarly, S leaf bundles were numbered based on the position of their respective axillary stem along the plant main stem, again starting from the bottom of plants (S1 to S10). To collect S leaf sections, biomass from adjacent leaf bundles was combined (S10 + S9, S8 + S7, and so on), avoiding axillary stems and leaf petioles as much as possible. The biomass from three randomly selected plants was collected and placed in pre-frozen 50 mL Falcon tubes, before flash freezing in liquid nitrogen. For sampling of the biomass for RNAseq, RTqPCR, and proteomics, P leaves of similar developmental stages were selected based on the leaf numbering described above. In line with the distribution of the S protein in the plant canopy (Figure S1), leaves P9 and P10 were selectively harvested without petiole. Freshly cut leaves were placed in pre-frozen 50 mL Falcon tubes, before flash freezing in liquid nitrogen. Each sample was made of six leaves collected on three randomly selected plants. To assess plant productivity following co-expression of KTIs, P and S leaves from whole plant shoots were manually detached, placed in aluminum foil, and flash frozen in liquid nitrogen. Each sample was made from the pooled leaf biomass of three randomly selected plants. For all sample sizes, frozen biomass was stored at -80°C until ready for analysis. Foliar tissues were then ground and homogenized to powder in liquid nitrogen using pre-chilled mortars and pestles. For all experiments, the average results presented were obtained from at least three biological replicates.

### Protein extraction and quantification

For protein extraction, 1 g of frozen leaf biomass powder was taken out of the - 80°C freezer and placed on ice. Two mL of extraction buffer (50 mM Tris, pH 8.0, 500 mM NaCl) was added, followed by 20 µL of 100 mM phenylmethanesulfonyl fluoride and 2 µL of 0.4 g/mL metabisulfite. Leaf samples were quickly crushed for 45 sec using a Polytron homogenizer (ULTRA-TURRAX^®^ T25 basic) at maximum speed. One mL of each sample was transferred to a prechilled Eppendorf tube and centrifuged at 10,000 x g for 10 min at 4°C. Supernatants were carefully recovered, transferred to new Eppendorf tubes, and kept on ice until protein quantification. Protein contents from crude extracts were determined according to the Bradford method, using bovine serum albumin as a protein standard.

### Western blotting and standardized capillary western immunoassays

For western blotting, total protein extracts were diluted in extraction buffer and mixed with 5X Laemmli sample loading buffer without reducing reagent. Dilution was performed to reach a final protein concentration of 0,5 µg/µL. Protein samples were denatured at 95°C for 5 min, followed by a quick spin using a microcentrifuge. Twenty µL of each denatured protein extract (10 µg) was next loaded on Criterion^TM^ XT Precast polyacrylamide gels 4-12% Bis-Tris and separated at 110 volts for 105 min. The proteins were then electrotransferred onto a polyvinylidene difluoride membrane at 100 volts using transfer buffer (25 mM Tris, pH 8.3, 192 mM glycine, 10% (v/v) methanol). After 30 min, the membranes were placed in a blocking solution comprising Tris-Buffered Saline (TBS-T; 50 mM Tris, pH 7.5, 150 mM NaCl, 0,1% (v/v) Tween-20), supplemented with 5% (w/v) nonfat dried milk. Membranes were blocked overnight at 4°C with gentle shaking and the blocking solution then removed. The membranes were incubated for 60 min at room temperature with gentle shaking in 1X TBS-T, 2% (w/v) nonfat dried milk solution containing primary antibodies. After four washes in 1X TBS-T, secondary antibodies were incubated for 60 min at room temperature with gentle shaking in 1X TBS-T, 2% (w/v) nonfat dried milk solution. After four extra washes in 1X TBS- T, the Luminata^TM^ Western HRP Chemiluminescence Substrate (Thermo Fisher Scientific) was added on the membranes and the protein complexes visualized under the chemiluminescence mode of an Imager 600 device (Amersham). The primary monoclonal antibody anti-SARS-CoV-2 Spike S1 (Sino Biological) was used for S protein detection at a dilution of 1/10,000. Goat anti-rabbit antibodies (JIR) were used as secondary antibody at a dilution of 1/10,000.

For relative quantification of the S protein, standardized capillary western immunoassays were performed using WES or JESS devices (ProteinSimple). For each microplate, a duplicated standard curve made of purified S protein was used. Standard curves comprised four points (2.0, 0.8, 0.32 and 0.128 ng/µL) and a duplicated S protein control of known concentration to validate the assay. To reach sample concentrations within the range of standard curves, protein extracts were diluted in TBS (pH 8.0) containing 0.1% (v/v) Tween-80, at dilutions ranging from 1/10 and 1/120 depending on the day of harvest. Standard curves, purified controls and CoVLP samples were denatured at 98°C for 10 min, followed by quick spin in a microcentrifuge. Five µL of denatured extract was loaded on pre-filled microplates (ProteinSimple), to which was added 10 µL of primary antibody, 10 µL of secondary antibody, 10 µL of antibody diluent, and 15 µL of chemiluminescence mix per microplate column. Anti-SARS-CoV-2 Spike S2 polyclonal antibodies (Novus Biologicals) were used as primary antibodies to detect the S protein and goat anti-rabbit antibodies (JIR) as secondary antibodies. Filled microplates were centrifuged for 5 min at 1,000 x g, before loading on the instrument. A new 12-230 kDa 25-capillaries cartridge was used for each experiment. For data capture, a protocol from the Compass for SW software was used, with the following specifications: separation matrix loaded for 200 sec, sample separation at 375 volts for 25 min, and incubation with antibody diluent, primary antibody, and secondary antibody for 5, 30, and 30 min, respectively. Chemiluminescence was detected using the high dynamic range protocol included in the software. Captured data was then transferred to Microsoft Excel for final analysis.

### RNA extraction and handling

For total RNA extraction, 100 mg of frozen leaf biomass powder was used as starting material for the RNeasy kit (Qiagen). Residual DNA was removed using the RNase-free DNase Set (Qiagen). For each extract, total RNA concentration was determined using a spectrophotometer (Implen) and RNA integrity evaluated using a 2100 BioAnalyzer (Agilent). For long-term storage, total RNA extracts were stabilized by adding the RNAseOUT recombinant ribonuclease inhibitor (Thermo Fisher Scientific), before freezing at -80°C until further analysis.

### RNAseq analysis

Total RNA extracts (see above) were used for RNAseq analyses. Three biological replicates of each tested condition and time point were sequenced. Prior to cDNA library construction, the mRNA fraction of total RNA extracts was enriched using Poly(A) columns. After fragmentation, stranded mRNA nucleotide libraries were prepared by fusing TruSeq RNA sequencing adapters (Illumina), followed by PCR amplification. cDNA libraries were sequenced using a NovaSeq 6000 device (Illumina), with runs of 2 × 150 bases. Sequenced reads were trimmed using the BBDuk software, available online through the BBtools suite (v. 38.79) (sourceforge.net/projects/bbmap/). Reads were aligned against the nuclear genome of *N. benthamiana* (https://solgenomics.net/), using the HISAT2 software (v. 2.2.0) (Kim *et al*., 2015). Duplicates were removed using the MarkDuplicates software available in the Genome Analysis Toolkit (v. 4.1.6.0) (McKenna *et al*., 2010). Based on aligned reads, fragments per kilobase of transcript per million mapped reads (FPKM) values were calculated using the Cuffdiff2 software (v. 2.2.1) (Trapnell *et al*., 2013). Once quantified, sequencing data was used to perform pairwise comparisons between conditions, as indicated. Previously published gene models were used as a basis for differential gene expression analyses (Sierro *et al*., 2014). Variance-stabilized read values were used to profile individual gene expression throughout the time series.

### RTqPCR analyses

For each sample, 1 µg of total RNA was reverse transcribed into cDNA using the QuantiTect Reverse Transcription Kit (Qiagen). Transcript quantification was performed in 96-well plates using the ABI PRISM 7500 Fast real-time PCR system and custom data analysis software (Thermo Fisher Scientific). Each reaction contained the equivalent of 5 ng cDNA template, 0.5 µM of forward and reverse primers, and 1X QuantiTect SYBR Green Master Mix (Qiagen), for a total reaction volume of 10 µL. Reactions were conducted under the SYBR Green amplification mode, under the following cycling conditions: 15 min incubation at 95°C, followed by 40 amplification cycles at 95°C for 5 sec, 60°C for 30 sec, and 65°C for 90 sec. Reactions in the absence of cDNA template were conducted as controls and melting curve analyses were performed to confirm amplification specificity and lack of primer dimer formation. Fluorescence levels and cycle threshold (Ct) values were next exported to the Microsoft Excel software for final analysis. To correct for biological variability and technical variations, the expression of six housekeeping genes (*NbACT1*, *NbVATP*, *NbSAND*, *NbUBQ1*, *NbEF1-α*, and *NbGAPDH1*) was monitored to normalize RTqPCR data (Vandesompele *et al*., 2002). Stable expression of housekeeping genes was confirmed by RTqPCR (Figure S12a) and using RNAseq data (Figure S12b). Normalized numbers of transcript molecules per ng of RNA were deduced using the 2^-ΔΔCt^ method (Livak and Schmittgen, 2001) and standard curves derived from known concentrations of phage lambda DNA. Standard deviation related to the within-treatment biological variation was calculated according to the error propagation rules. Primers used for RTqPCR analyses are available in Table S18.

### Large-scale proteomics

To study changes in protein abundance, total protein extracts derived from leaves P9 and P10 were used for iTRAQ labeling. Three biological replicates from each condition were analysed. Leaf biomass was first reduced to powder in liquid nitrogen using a cryoPREP Automated Dry Pulverizer (Covaris). Using a commercial extraction buffer (Bio-Rad), proteins were then extracted with a TissueLyser II (Qiagen). Extracted proteins were precipitated with acetone and resuspended in 1 M urea, 0.5 M TEAB containing 0.1% (w/v) SDS. Protein concentrations were next determined according to the Bradford method and the samples were normalized to 50 µg of protein in a volume of 30 µL. A 30 µL volume of PreOmics LYSE-NHS 2X buffer was added to denature, reduce, and alkylate the proteins. The resulting mixture was incubated for 10 min at 60°C, followed by centrifugation at 300 x g for 1 min. To digest proteins, 50 µL of PreOmics DIGEST solution was next added to the supernatant, followed by incubation for 3 h at 37°C. The reaction was stopped by adding 250 µL of PreOmics STOP solution (pH below 3), and the reaction mixture was purified following the plant kit protocol from PreOmics. Sample peptides were vacuum dried in a SpeedVac (Thermo Fisher Scientific) and injected into a Nano-LC-Q-Exactive HF mass spectrometer (Thermo Fisher Scientific). Direct data-independent (DIA) quantification was performed against a *N. benthamiana* protein database using the Spectronaut software (Biognosys). Common peptide contaminants such as keratin were taken out of the analysis.

### Measurement of the CP activity

The activity of CPs was monitored using the synthetic fluorogenic peptide *Z*-Phe-Arg 7-amido-4-methylcoumarin (*Z*-Phe-Arg-MCA) (Pepnet), a preferred substrate for CPs of the cathepsin L family. The CP activity of each leaf section was deduced from comparisons with standard curves derived from the activity of recombinant human cathepsin L (Abcam), using the enzymatic unit (U) definition provided by the manufacturer. More details on assay buffer, standard curve preparation, sample dilutions, and data capture are provided in our accompanying manuscript (Hamel *et al*., 2025).

### Phylogenetic analyses

The genome of *N. benthamiana* was searched using full-length amino acid sequences of Arabidopsis AtKTI1 and AtKTI5 as queries to identify members of the KTI family. Retrieved full-length protein sequences were aligned with ClustalW, using the following alignment parameters: for pairwise alignments, 10.0 for gap opening and 0.1 for gap extension; for multiple alignments, 10.0 for gap opening and 0.2 for gap extension. The resulting alignment was submitted to the MEGA5 software and a neighbor-joining tree derived from 5,000 replicates was generated. Bootstrap values are indicated on the node of each branch.

### Statistical analyses

Statistical analyses on standardized capillary western immunoassays and RTqPCR results were performed on Graph Pad Prism 10.1.0 to assess the effects of KTI co-expression. Groups (treatments) were analysed using one-way ANOVA followed by a post-hoc Tukey’s multiple comparison test with an alpha threshold of 0.05. Groups are labeled with a compact letter display. Groups that do not share the same letter are statistically different.

### Accession numbers

The RNAseq dataset used in this study has been deposited in the European Nucleotide Archive (ENA) database (https://www.ebi.ac.uk/ena/browser/home), under accession number PRJEB64341.

## Acknowledgments and funding

This work would not have been possible without the support of the Biomass Production and Research & Innovation teams of Medicago. Contributions included, but were not limited to, gene construct cloning, inoculum preparation, plant cultivation, and leaf agroinfiltration. The authors also wish to acknowledge Cecilia Cheval, James Nicolas Duncan Battey, Lucien Bovet, and Simon Goepfert (current or former employees of Philip Morris International) for assistance with study design, sequencing data acquisition, and helpful discussions. Funding for this work was provided by Medicago Inc.

## Author contributions

L.P.H., D.M., and M.A.D. designed the research, supervised the project, and analysed the data. L.P.H. and M.A.C. drafted the manuscript and assembled the figures. P.O.L. managed production of the genetic constructs. L.P.H., F.P.G., R.T., M.E.P., and M.A.C. produced biomass and extracts used for transcriptomics and proteomics analyses. F.P.G., R.T., M.E.P., and M.A.C. performed RTqPCR, recombinant protein quantification, and western blots. L.P.H., M.A.C., and F.P.G. managed RNAseq data and performed gene expression profiling. M.E.P. performed CoVLP purification and TEM imaging. M.C.G., A.L., M.E.P., and M.A.C. performed quantification of the CP activity. L.P.H. and M.C.G. performed phylogenetic analysis and protein sequence alignments. M.E.P. and L.P.H. conducted KTI co-expression experiments. M.A.C., M.C.G., and F.P.G. performed statistical analyses. All authors read, helped to edit, and approved final version of the manuscript.

## Data availability statement

All data discussed in this study can be found in the manuscript or the Supplementary Materials online.

## Supporting information

Additional supporting information may be found online in the Supporting Information section at the end of the article.

**Figure S1.**
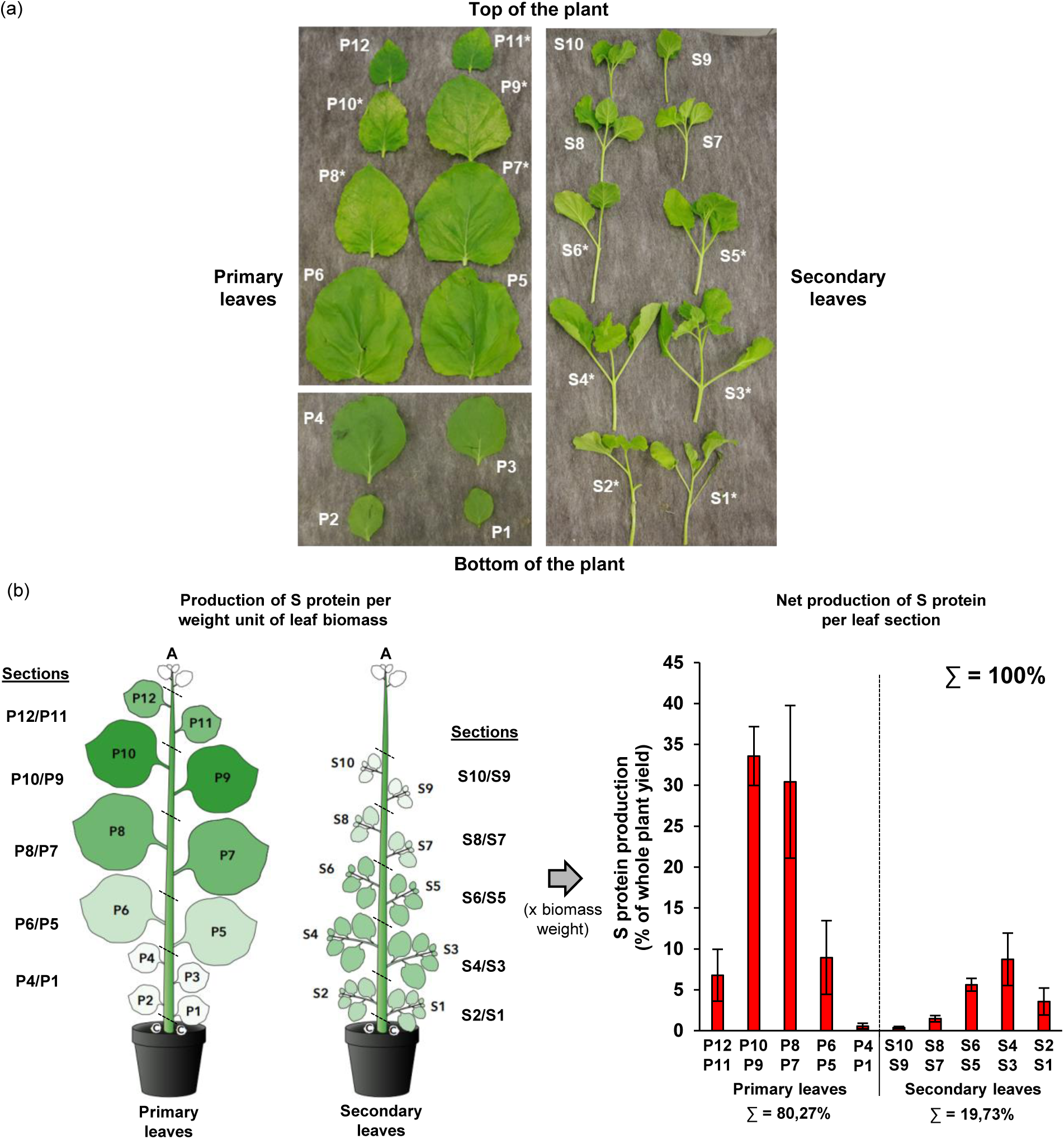
Distribution of the S protein in the plant canopy. (a) Stress symptoms observed on leaves of a representative CoVLP plant at 6 DPI. As symptoms were not evenly distributed, primary (P) leaves were numbered P1 to P12 based on their position along the main stem, starting from the bottom of plants. Similarly, secondary (S) leaf bundles were numbered S1 to S10 based on the position of their axillary stem along the plant main stem, again starting from the bottom of plants. Asterisks (*) denote leaves where greyish necrotic flecking was visible, although symptoms were more severe on leaves P8, P9, and P10. (b) Schematic view of the plant canopy and distribution of the S protein within five primary and five secondary leaf sections, as indicated. Left panel: standardized capillary western immunoassays were used to define productivity of each leaf section. The green colored gradient reflects S protein accumulation per weight unit of leaf biomass. For each section, displayed leaf surface is proportional to the weight of harvested biomass. Abbreviations: apex (A), cotyledon (C). Right panel: to define final yield contribution of each leaf section, S protein accumulation per weight unit of leaf biomass was multiplied by leaf biomass weight in each section. Results are expressed as a percentage (%) of the overall S protein yield (whole plant canopy), a total arbitrarily set at 100%.

**Figure S2.**
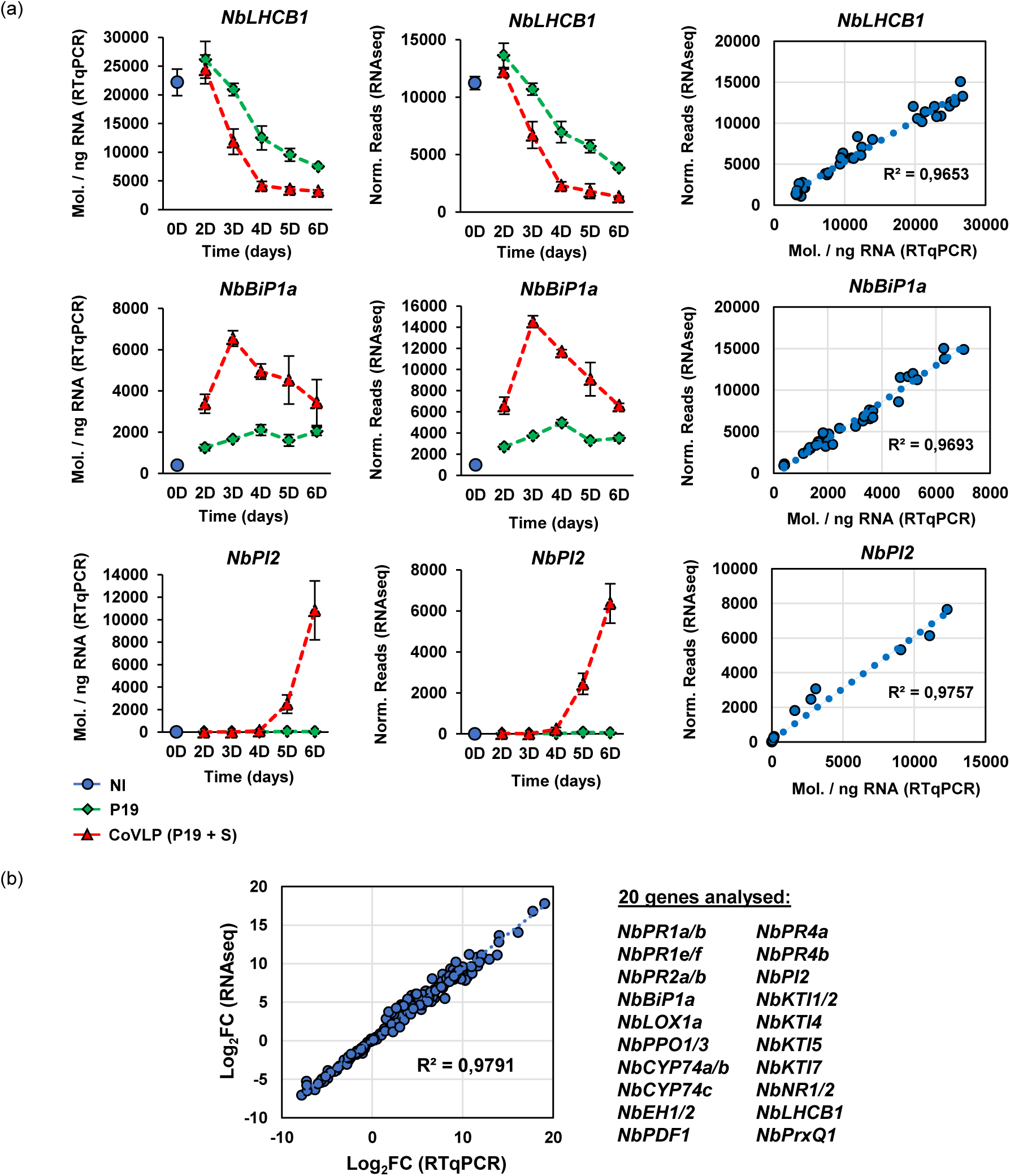
Correlation between RNAseq and RTqPCR data. To validate RNAseq data, RTqPCR was performed on 20 plant genes with different functions, expression levels, or transcriptional behaviors in the tested conditions. Analysed candidates included photosynthesis-related gene *NbLHCB1*, ERQC-related gene *NbBiP1a*, and oxylipin response gene *NbPI2* (a). For each time point in days (D) post-infiltration, RTqPCR results are expressed in numbers of transcript molecules per ng of RNA ± se (left panels). For RNAseq data (middle panels), results are expressed in normalized (norm.) numbers of reads ± sd. For each gene analysed, LR and deduced R^2^ highlight correlation between the independent methods (right panels). (b) Overall LR and R^2^ value were also deduced from expression data of the 20 genes analysed by RTqPCR (genes listed on the right). To account for variable expression levels among the tested genes, results are expressed as Log_2_FC values compared to the expression level in NI samples at 0 DPI. Conditions are as follows: NI: non-infiltrated samples (blue); P19: agroinfiltrated samples expressing P19 only (green); CoVLP: agroinfiltrated samples co-expressing P19 and recombinant S protein (red).

**Figure S3.**
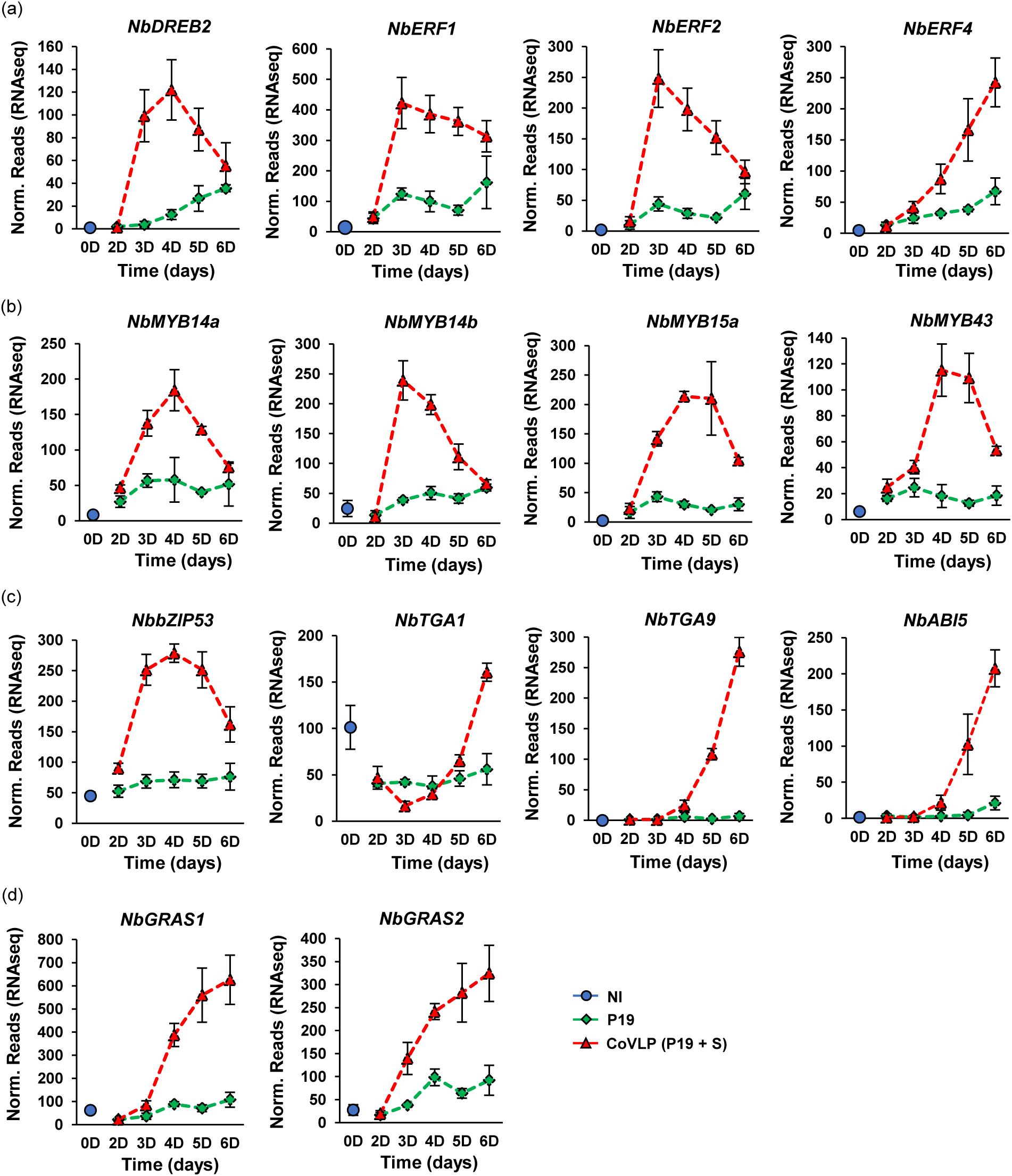
Upregulation of other TF genes. Expression profiles of genes encoding TFs of the AP2/ERF (a), MYB (b), bZIP (c), or GRAS (d) families. For each time point in days (D) post-infiltration, RNAseq results are expressed in normalized (norm.) numbers of reads ± sd. Conditions are as follows: NI: non-infiltrated samples (blue); P19: agroinfiltrated samples expressing P19 only (green); CoVLP: agroinfiltrated samples co-expressing P19 and recombinant S protein (red).

**Figure S4.**
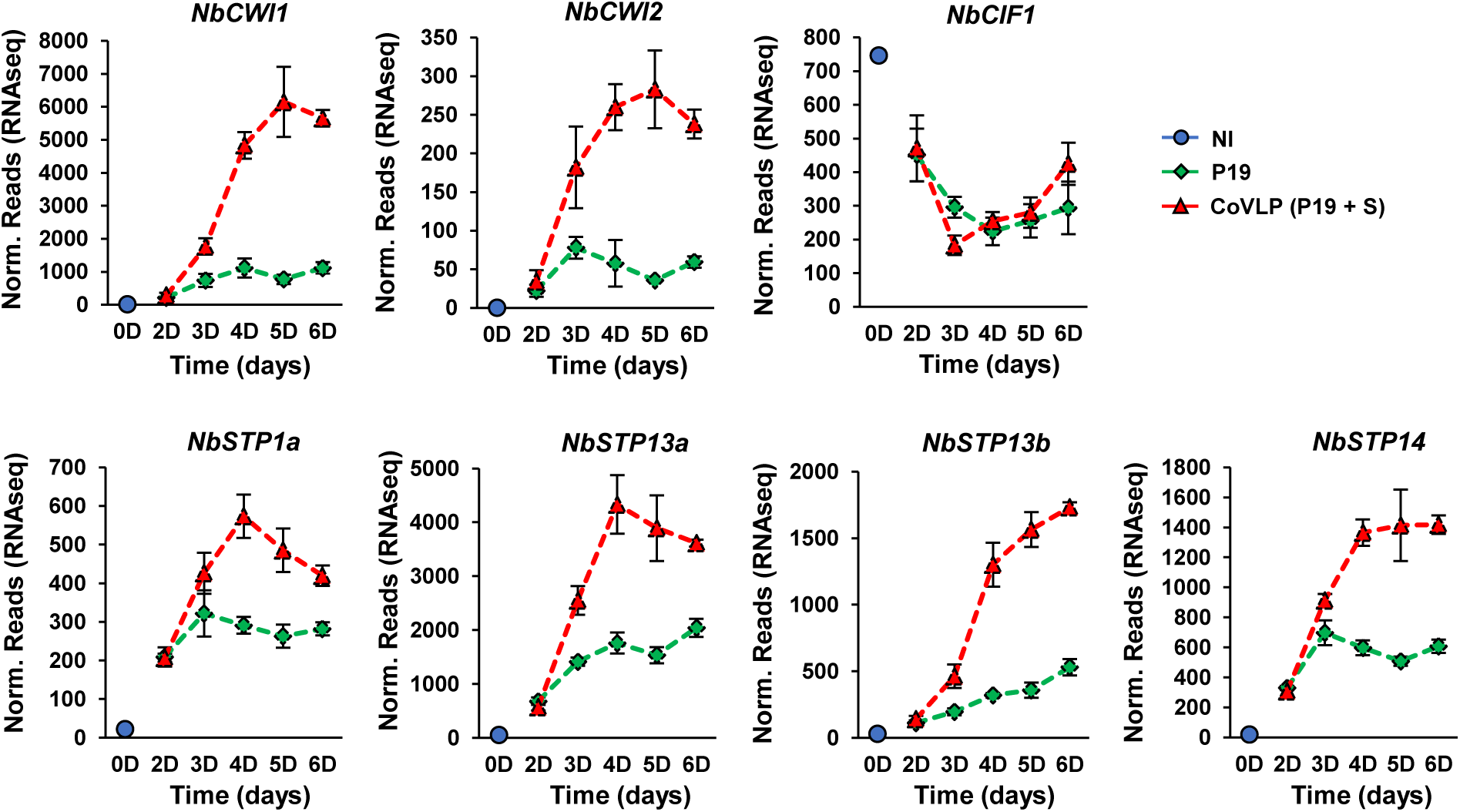
Expression of sugar-related genes. Expression profiles of CWI-encoding genes *NbCWI1* and *NbCWI2*. Expression profiles of genes encoding CWI inhibitor NbCIF1 and stress-related STPs are also shown. For each time point in days (D) post-infiltration, RNAseq results are expressed in normalized (norm.) numbers of reads ± sd. Conditions are as follows: NI: non-infiltrated samples (blue); P19: agroinfiltrated samples expressing P19 only (green); CoVLP: agroinfiltrated samples co-expressing P19 and recombinant S protein (red).

**Figure S5.**
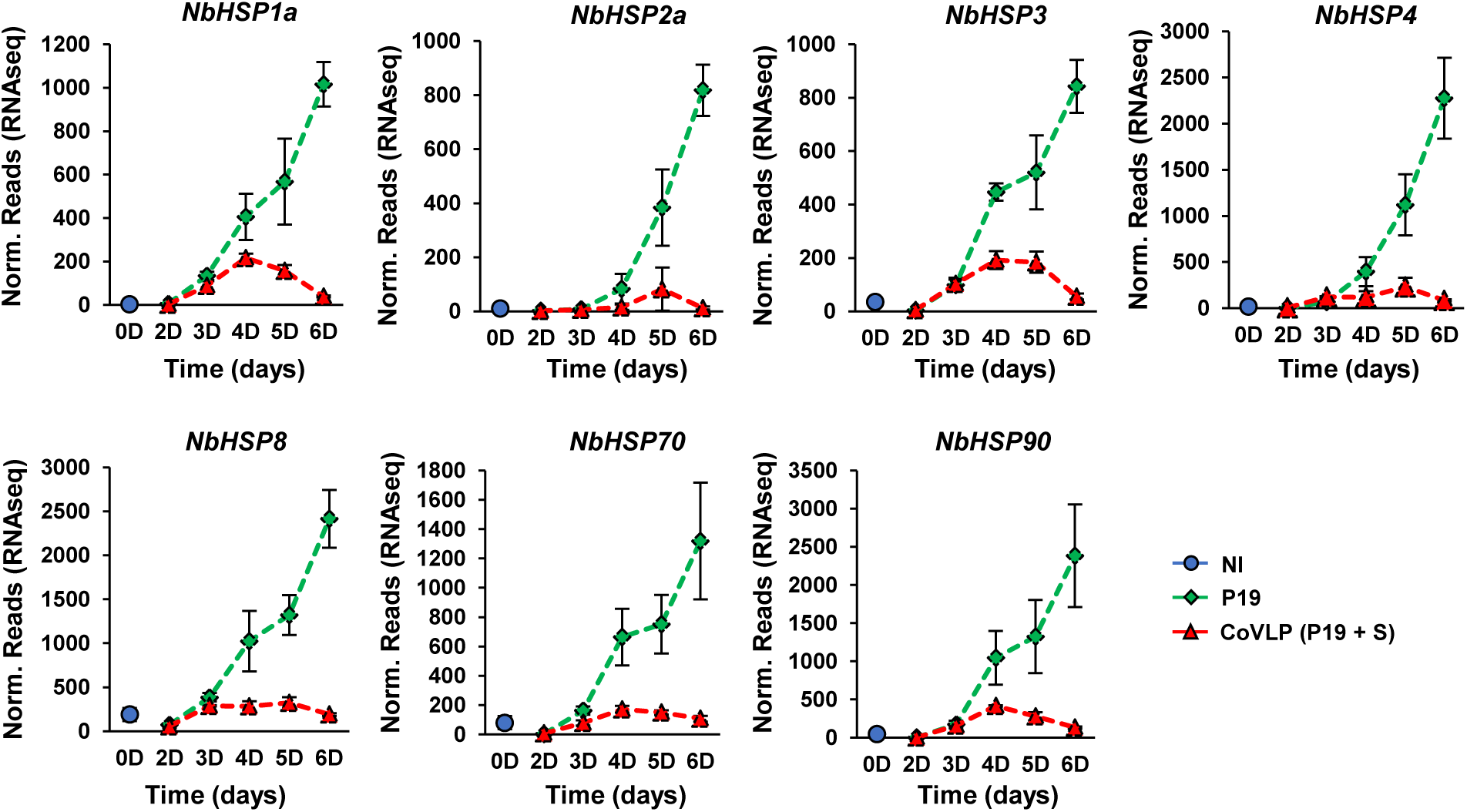
P19-mediated upregulation of *HSP* genes. Expression profiles of genes encoding various subtypes of cytosolic HSPs, namely small HSPs, HSP70 and HSP90. For each time point in days (D) post-infiltration, RNAseq results are expressed in normalized (norm.) numbers of reads ± sd. Conditions are as follows: NI: non-infiltrated samples (blue); P19: agroinfiltrated samples expressing P19 only (green); CoVLP: agroinfiltrated samples co-expressing P19 and recombinant S protein (red).

**Figure S6.**
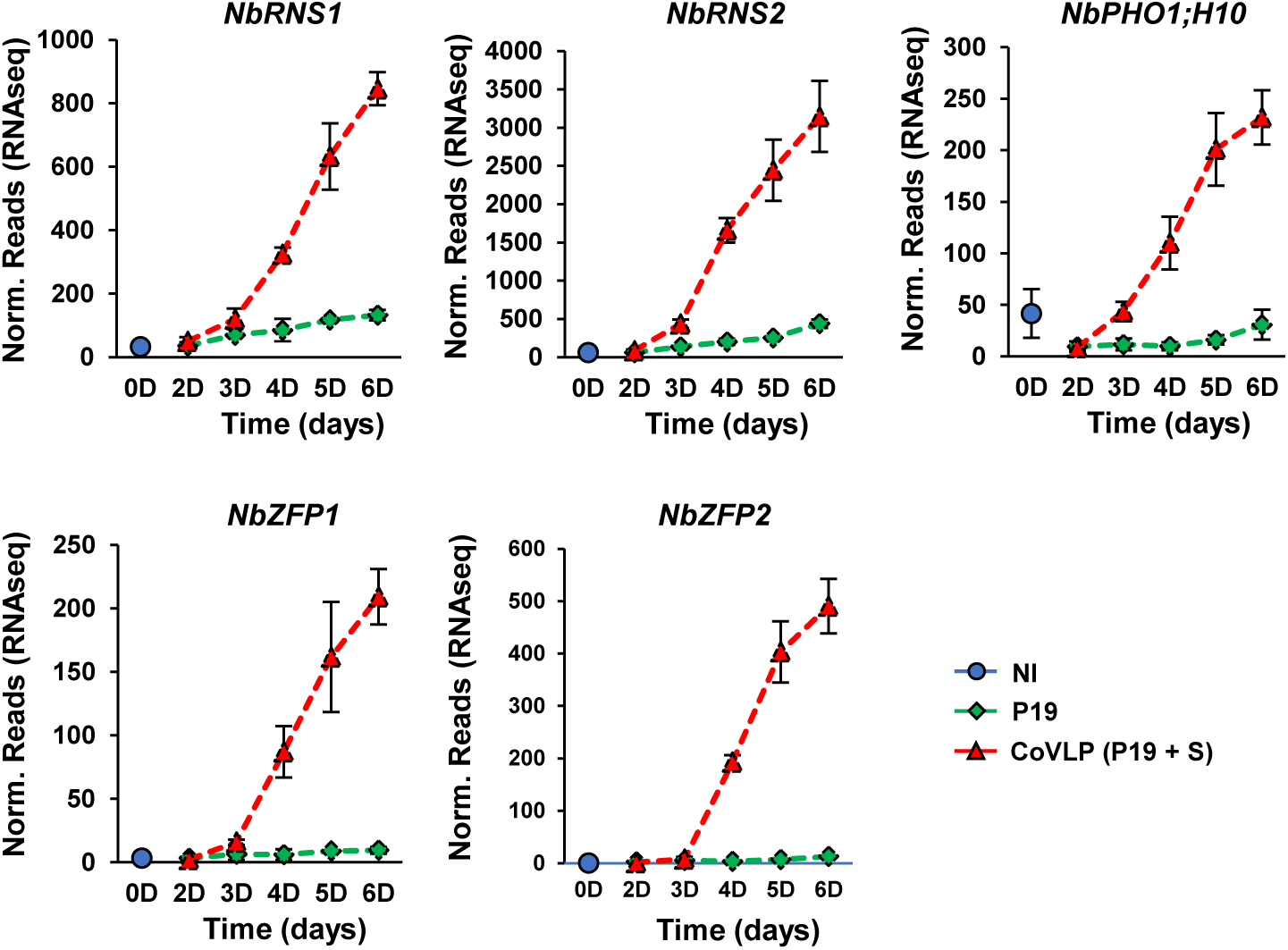
Upregulation of OPDA-specific genes. Expression profiles of genes specifically induced by OPDA, namely ribonuclease genes *NbRNS1* and *NbRNS2*, phosphate transporter gene *NbPHO1;H10*, as well as ZFP genes *NbZFP1* and *NbZFP2*. For each time point in days (D) post-infiltration, RNAseq results are expressed in normalized (norm.) numbers of reads ± sd. Conditions are as follows: NI: non-infiltrated samples (blue); P19: agroinfiltrated samples expressing P19 only (green); CoVLP: agroinfiltrated samples co-expressing P19 and recombinant S protein (red).

**Figure S7.**
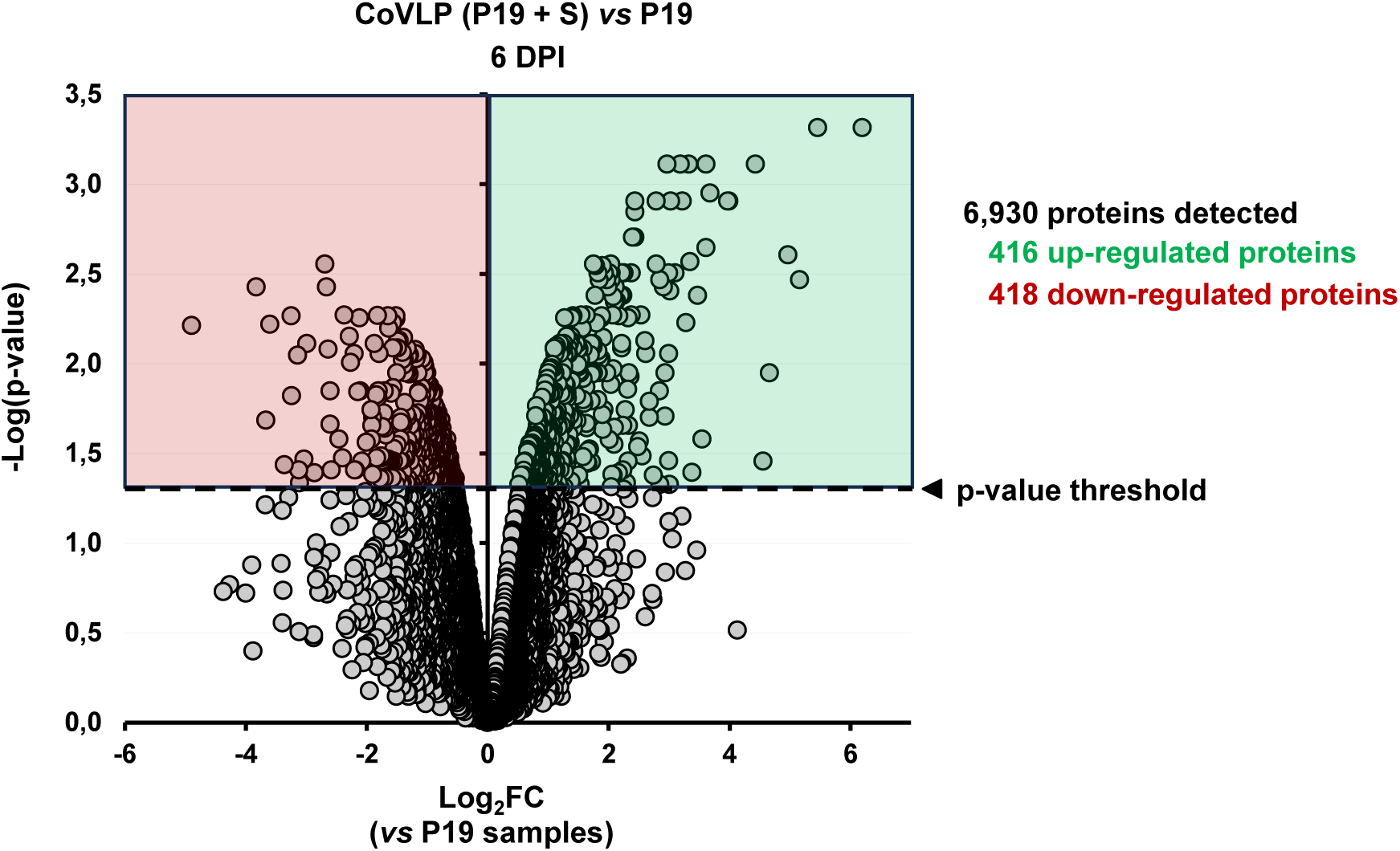
Overview of the proteomics results. Global translational changes as depicted by a volcano plot of all sequenced proteins at 6 DPI. Pairwise comparison was performed for CoVLP samples, using P19 samples as a control. Differentially expressed proteins were defined as all proteins whose differential expression adjusted p-value (padj) was below 0.05. Upregulated proteins are shown in green, downregulated proteins in red. A dashed line highlights the p-value threshold.

**Figure S8.**
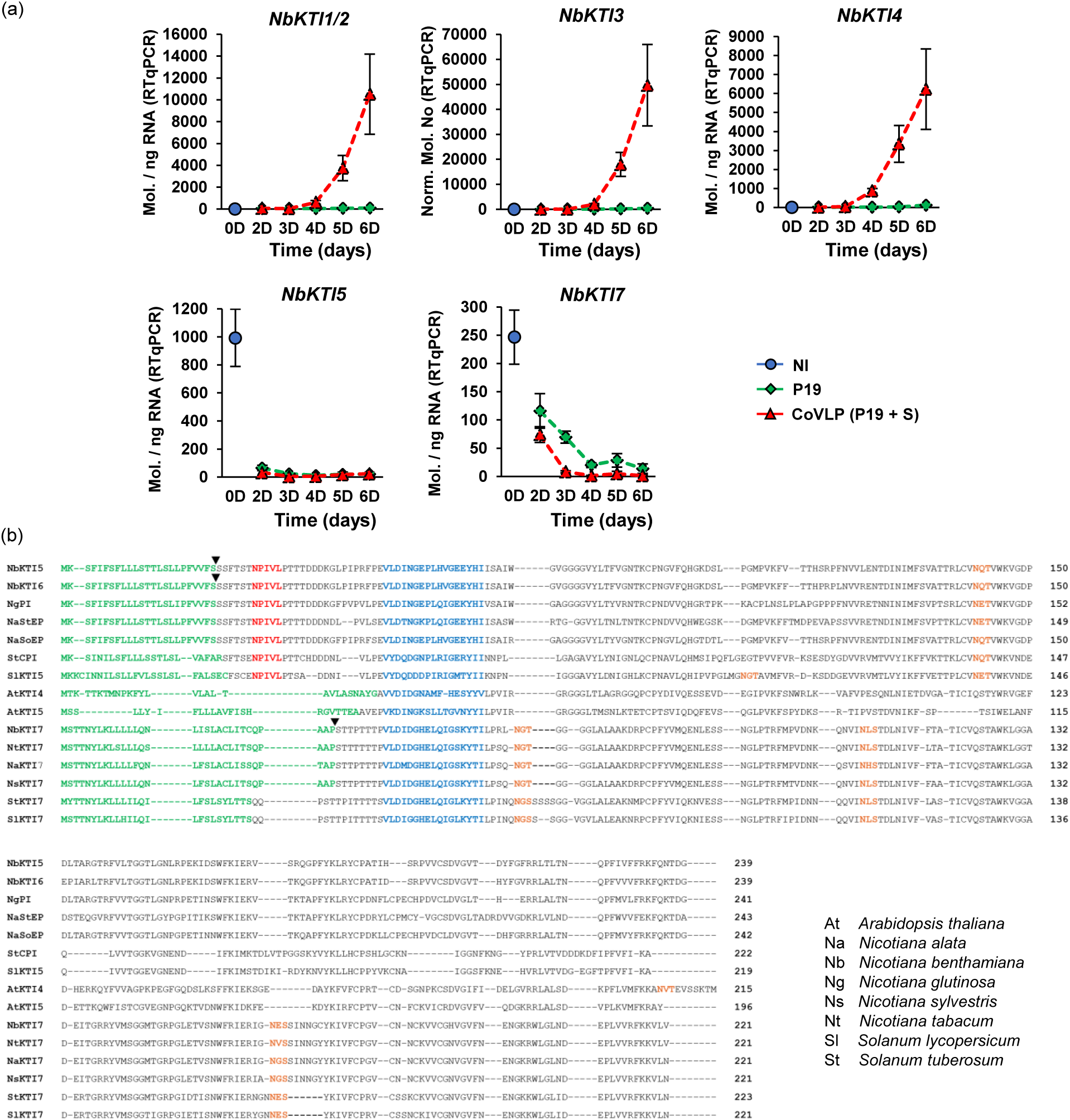
Confirmation of *NbKTI* expression profiles and protein alignment of NbKTI5 and NbKTI7 homologs. (a) Expression profiles of up- and downregulated *NbKTIs* as measured by RTqPCR. For each time point in days (D) post-infiltration, results are expressed in numbers of transcript molecules per ng of RNA ± se. Conditions are as follows: NI: non-infiltrated samples (blue); P19: agroinfiltrated samples expressing P19 only (green); CoVLP: agroinfiltrated samples co-expressing P19 and recombinant S protein (red). (b) Protein sequence alignment of full-length KTIs from various plant species (two-letter species acronyms shown at the bottom right). Amino acids are depicted using their one-letter code and positions within each protein are provided at the right of each sequence. Predicted SPs for protein secretion are highlighted in green, with black arrows highlighting probable SP cleavage sites of *N. benthamiana* KTIs. Vacuolar sorting signals are highlighted in red, while conserved peptides specific to Kunitz inhibitors (PI of family I03) are shown in blue. Predicted N-glycosylation sites are depicted in orange. Accession numbers: NgPI (AF208020); NaStEP (EU253563); NaSoEP (EU253564); StCPI (P20347); SlKTI5 (XP_004235410); NtKTI7 (XP_009601782); NaKTI7 (XP_019266547); NsKTI7 (XP_009758946); StKTI7 (XP_015167657); SlKTI7 (XP_004237191).

**Figure S9.**
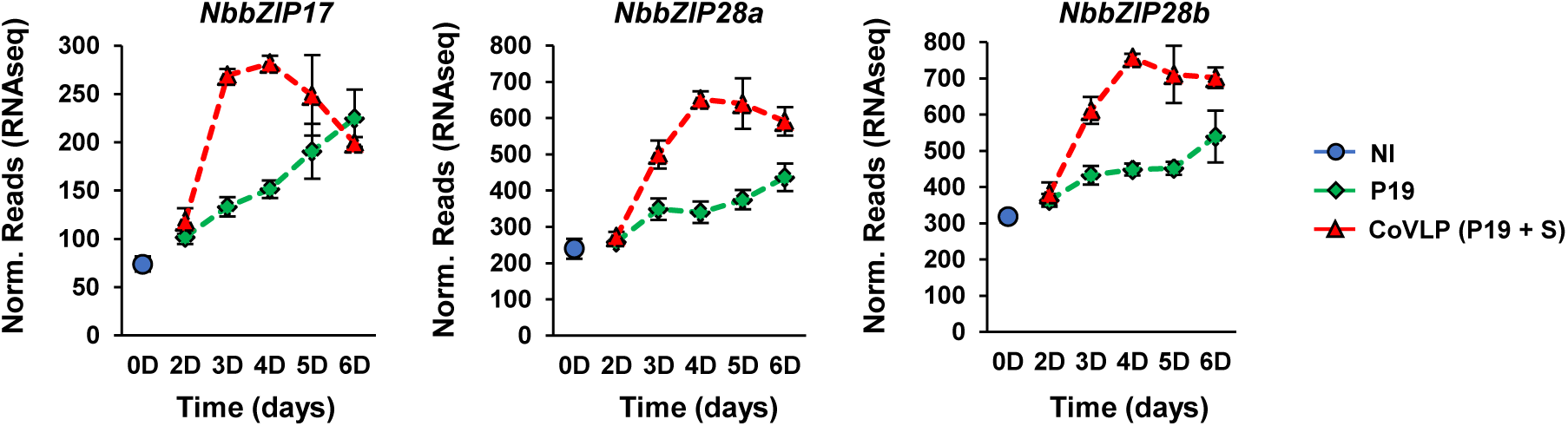
Upregulation of other UPR-related *bZIPs*. Expression profiles of TF genes encoding membrane-tethered bZIPs that are involved in the second branch of the plant UPR. For each time point in days (D) post-infiltration, RNAseq results are expressed in normalized (norm.) numbers of reads ± sd. Conditions are as follows: NI: non-infiltrated samples (blue); P19: agroinfiltrated samples expressing P19 only (green); CoVLP: agroinfiltrated samples co-expressing P19 and recombinant S protein (red).

**Figure S10.**
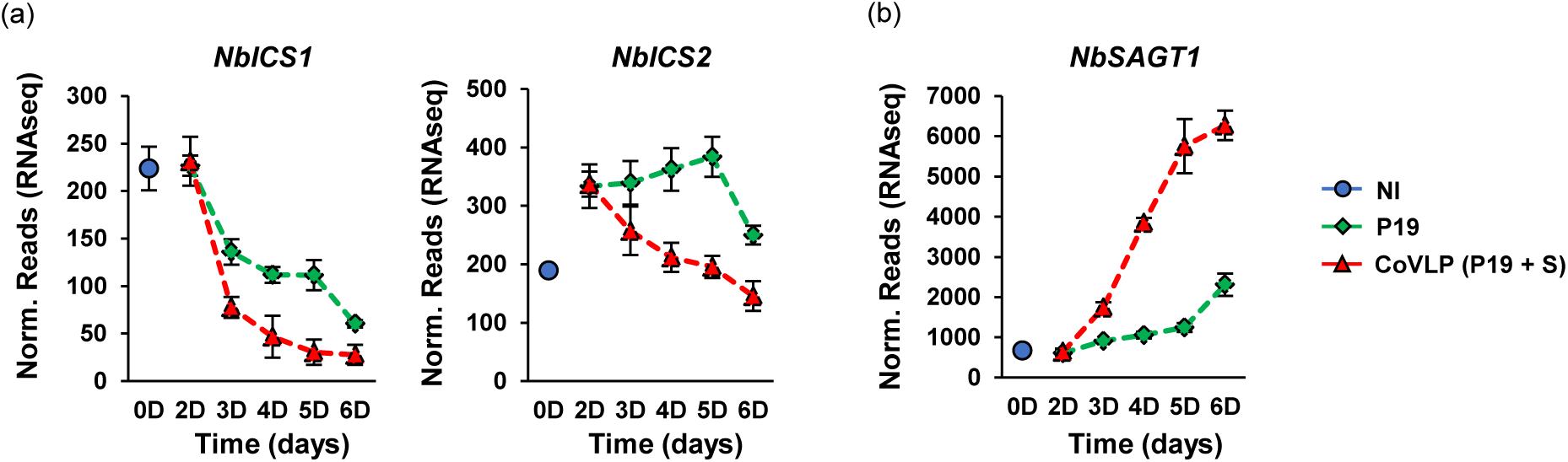
Expression of SA regulatory genes. (a) Expression profiles of *ICS* genes encoding enzymes involved in SA biosynthesis. Also shown is the expression profile of *NbSAGT1*, which encodes an enzyme converting SA into its inactive conjugated forms (b). For each time point in days (D) post-infiltration, RNAseq results are expressed in normalized (norm.) numbers of reads ± sd. Conditions are as follows: NI: non-infiltrated samples (blue); P19: agroinfiltrated samples expressing P19 only (green); CoVLP: agroinfiltrated samples co-expressing P19 and recombinant S protein (red).

**Figure S11.**
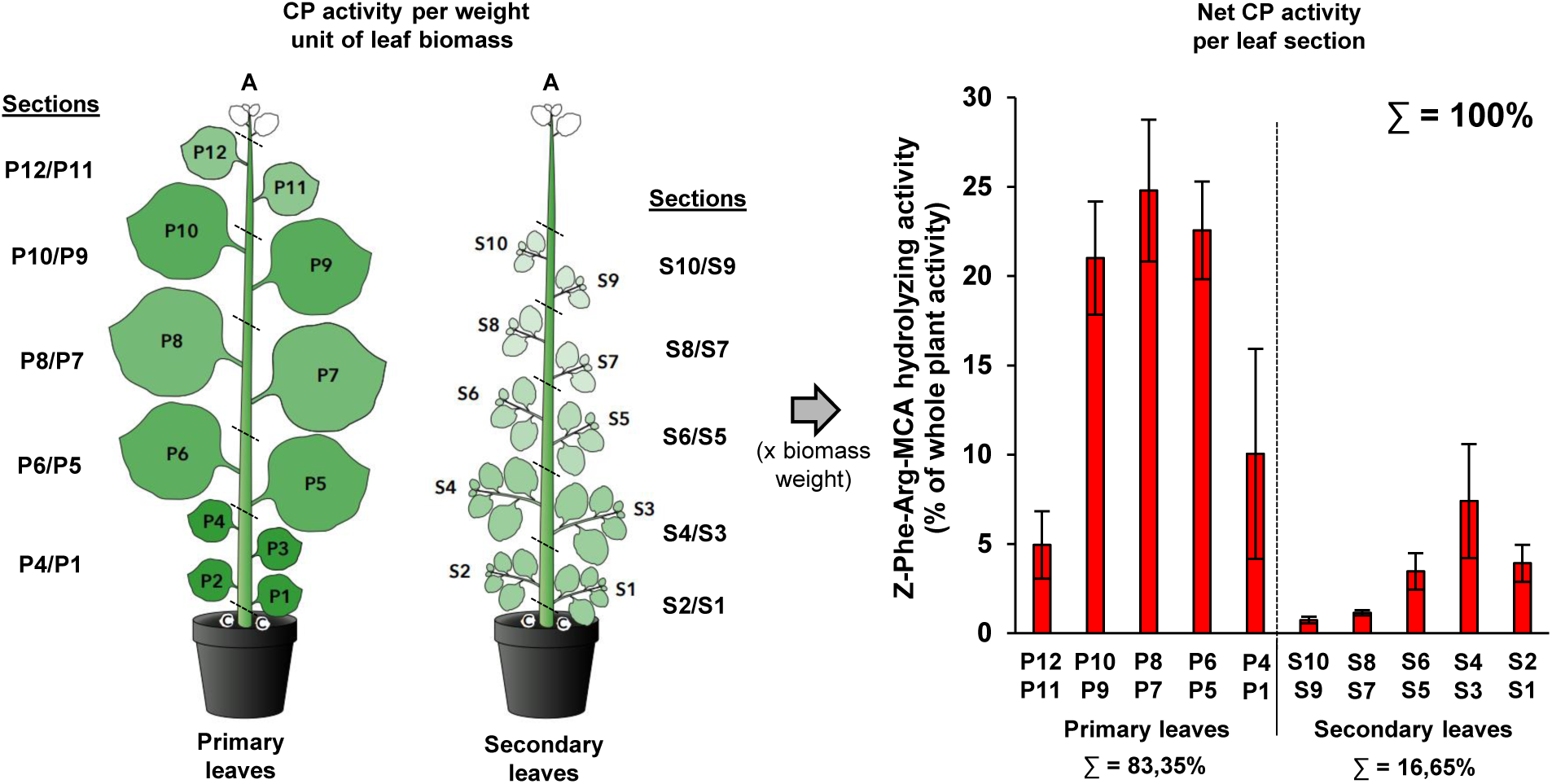
Distribution of CP activity in the plant canopy. Schematic view of the plant canopy and distribution of the CP activity within five primary (P) and five secondary (S) leaf sections, as indicated. Left panel: hydrolysis of the synthetic fluorogenic substrate *Z*-Phe-Arg-MCA and standard curves made with known concentrations of recombinant human cathepsin L were used to define CP activity of each leaf section. The green colored gradient reflects substrate hydrolyzing activity per weight unit of leaf biomass. For each section, displayed leaf surface is proportional to the weight of harvested biomass. Abbreviations: apex (A), cotyledon (C). Right panel: to define net CP activity of each leaf section, substrate hydrolyzing activity per weight unit of leaf biomass was multiplied by leaf biomass weight in each section. Results are expressed as a percentage (%) of the overall CP activity (whole plant canopy), a total arbitrarily set at 100%.

**Figure S12.**
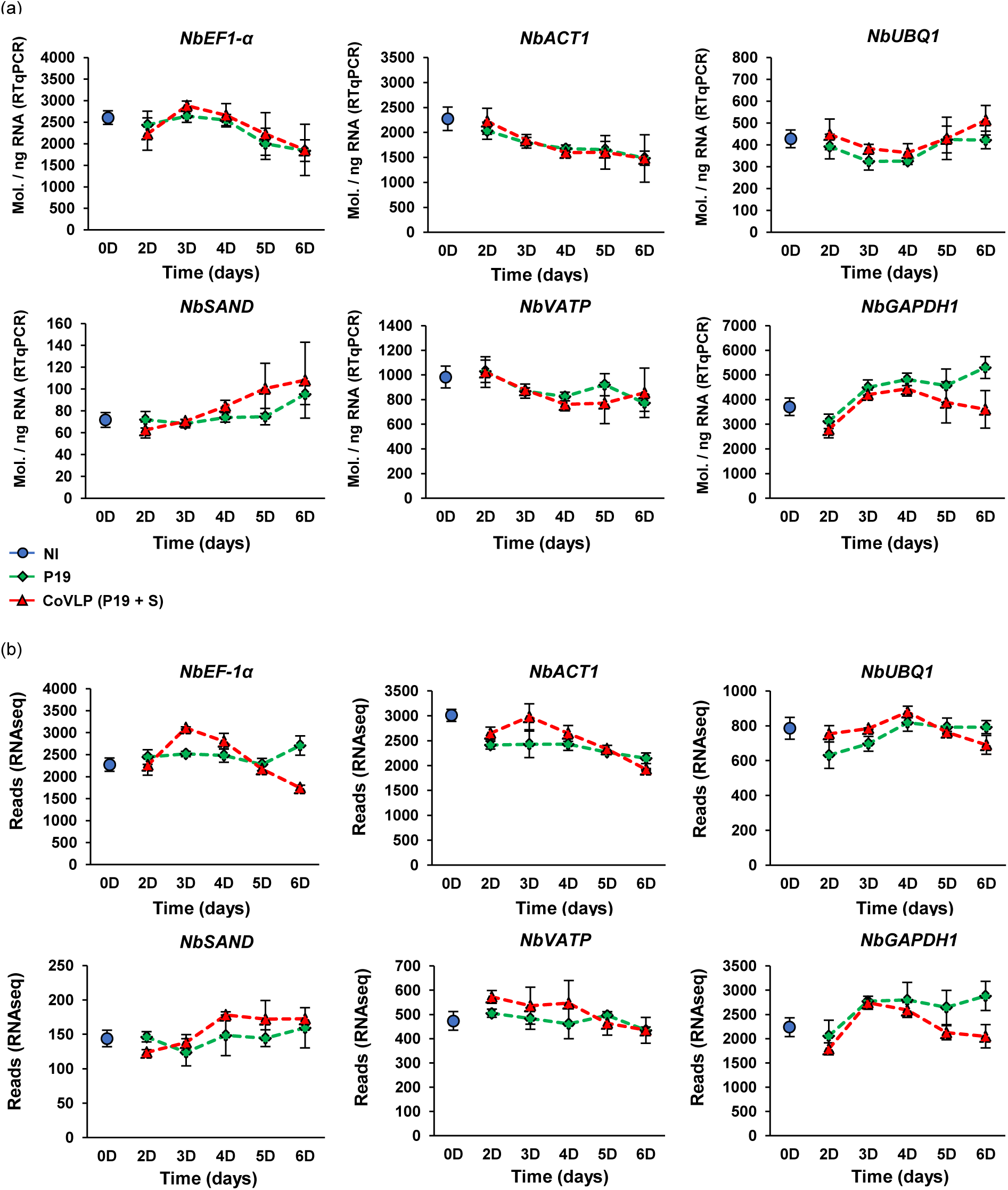
Expression of reference genes. Expression profiles of reference genes used to normalize RTqPCR results. Stable expression was verified using RTqPCR measurements (a), or RNAseq data (b). For each time point in days (D) post-infiltration, RTqPCR results are expressed in numbers of transcript molecules per ng of RNA ± se. RNAseq results are expressed in raw number of reads ± sd. Conditions are as follows: NI: non-infiltrated samples (blue); P19: agroinfiltrated samples expressing P19 only (green); CoVLP: agroinfiltrated samples co-expressing P19 and recombinant S protein (red).

**Table S1.** Unsorted lists of genes with significantly altered expression in P19 and CoVLP samples at 2 DPI.

**Table S2.** Unsorted lists of genes with significantly altered expression in P19 and CoVLP samples at 3 DPI.

**Table S3.** Unsorted lists of genes with significantly altered expression in P19 and CoVLP samples at 6 DPI.

**Table S4.** Unsorted lists of significantly regulated genes specific to P19 and CoVLP samples, or common to both conditions at 2 DPI.

**Table S5.** Unsorted lists of significantly regulated genes specific to P19 and CoVLP samples, or common to both conditions at 3 DPI.

**Table S6.** Unsorted lists of significantly regulated genes specific to P19 and CoVLP samples, or common to both conditions at 6 DPI.

**Table S7.** UPR genes significantly regulated in P19 and CoVLP samples at various time points.

**Table S8.** TF genes significantly regulated in P19 and CoVLP samples at various time points.

**Table S9.** Lignin-related genes significantly regulated in P19 and CoVLP samples at various time points.

**Table S10.** Terpene-related genes significantly regulated in P19 and CoVLP samples at various time points.

**Table S11.** Oxidative stress- and sugar-related genes significantly regulated in P19 and CoVLP samples at various time points.

**Table S12.** SA- and SAR-related genes significantly regulated in P19 and CoVLP samples at various time points.

**Table S13.** *HSP* genes significantly regulated in P19 and CoVLP samples at various time points.

**Table S14.** Oxylipin-related genes significantly regulated in P19 and CoVLP samples at various time points.

**Table S15.** Unsorted lists of proteins with significantly altered expression in CoVLP samples at 6 DPI.

**Table S16.** Overlap between molecular responses induced by influenza VLPs and CoVLPs.

**Table S17.** Lipid-related genes significantly regulated in P19 and CoVLP samples at various time points.

**Table S18.** RTqPCR primers used in this study.

**Appendix S1.** Definitions of all abbreviations used in this study.

## References

Adachi, H., Nakano, T., Miyagawa, N., Ishihama, N., Yoshioka, M., Katou, Y., Yaeno, T. et al. (2015) WRKY transcription factors phosphorylated by MAPK regulate a plant immune NADPH oxidase in *Nicotiana benthamiana*. Plant Cell 27, 2645–2663.

Anon. (2022) Editorial - Crop pharming. *Nat. Plants* **8**, 191.

Arnaiz, A., Talavera-Mateo, L., Gonzalez-Melendi, P., Martinez, M., Diaz, I. and Santamaria, M.E. (2018). Arabidopsis Kunitz trypsin inhibitors in defense against spider mites. Front. Plant Sci. 9, 986.

Asai, T., Tena, G., Plotnikova, J., Willmann, M.R., Chiu, W.L., Gomez-Gomez. L., Boller, T., et al. (2002) MAP kinase signalling cascade in *Arabidopsis* innate immunity. Nature 415, 977–983.

Baden, L.R., El Sahly, H.M., Essink, B., Kotloff, K., Frey, S., Novak, R., Diemert, D. et al. (2021) Efficacy and safety of the mRNA-1273 SARS-CoV-2 Vaccine. N. Engl. J. Med. 384, 403–416.

Bakshi, M. and Oelmüller, R. (2014) WRKY transcription factors: Jack of many trades in plants. Plant Signal. Behav. 9, e27700.

Bally, J., Jung, H., Mortimer, C., Naim, F., Philips, J.G., Hellens, R., Bombarely, A. et al. (2018) The rise and rise of *Nicotiana benthamiana*: A plant for all reasons. Annu. Rev. Phytopathol. 56, 405–426.

Bateman, A., Finn, R.D., Sims, P.J., Wiedmer, T., Biegert, A. and Soding, J. (2009) Phospholipid scramblases and Tubby-like proteins belong to a new superfamily of membrane tethered transcription factors. Bioinformatics 25, 159–162.

Benchabane, M., Schlüter, U., Vorster, J., Goulet, M.C. and Michaud, D. (2010) Plant cystatins. Biochimie 92, 1657–1666.

Boeckx, T., Webster, R., Winters, A.L., Webb, K.J., Gay, A. and Kingston-Smith, A.H. (2015) Polyphenol oxidase-mediated protection against oxidative stress is not associated with enhanced photosynthetic efficiency. Ann. Bot. 116, 529–540.

Bosch, M., Wright, L.P., Gershenzon, J., Wasternack, C., Hause, B., Schaller, A. and Stintzi, A. (2014) Jasmonic acid and its precursor 12-oxophytodienoic acid control different aspects of constitutive and induced herbivore defenses in tomato. Plant Physiol. 166, 396–410.

Cacas, J.L., Vailleau, F., Davoine, C., Ennar, N., Agnel, J.P., Tronchet, M., Ponchet, M. et al. (2005) The combined action of 9 lipoxygenase and galactolipase is sufficient to bring about programmed cell death during tobacco hypersensitive response. Plant Cell Environ. 28, 1367–1378.

Canonne, J., Froidure-Nicolas, S. and Rivas, S. (2011) Phospholipases in action during plant defense signaling. Plant Signal. Behav. 6, 13–18.

Carter, C.J. and Thornburg, R.W. (2004) Tobacco nectarin V is a flavin-containing berberine bridge enzyme-like protein with glucose oxidase activity. Plant Physiol. 134, 460–469.

Catinot, J., Buchala, A., Abou-Mansour, E. and Metraux, J.P. (2008) Salicylic acid production in response to biotic and abiotic stress depends on isochorismate in *Nicotiana benthamiana*. FEBS Lett. 582, 473–478.

Cecchini, N.M., Steffes, K., Schlappi, M.R., Gifford, A.N. and Greenberg, J.T. (2015) *Arabidopsis* AZI1 family proteins mediate signal mobilization for systemic defence priming. Nat Commun. 6, 7658.

Chung, Y.H., Church, D., Koellhoffer, E.C., Osota, E., Shukla, S., Rybicki, E.P., Pokorski, J.K. et al. (2022) Integrating plant molecular farming and materials research for next-generation vaccines. Nat. Rev. Mater. 7, 372–388.

Daniel, B., Konrad, B., Toplak, M., Lahham, M., Messenlehner, J., Winkler, A. and Macheroux, P. (2017) The family of berberine bridge enzyme-like enzymes: A treasure-trove of oxidative reactions. Arch. Biochem. Biophys. 632, 88–103.

D’Aoust, M.A., Lavoie, P.O., Couture, M.M., Trepanier, S., Guay, J.M., Dargis, M., Mongrand, S. et al. (2008) Influenza virus-like particles produced by transient expression in *Nicotiana benthamiana* induce a protective immune response against a lethal viral challenge in mice. Plant Biotechnol. J. 6, 930–940.

Dean, J.V. and Delaney, S.P. (2008) Metabolism of salicylic acid in wild-type, *ugt74f1* and *ugt74f2* glucosyltransferase mutants of *Arabidopsis thaliana*. Physiol. Plant. **132**, 417-425.

Demidchik, V. (2015) Mechanisms of oxidative stress in plants: From classical chemistry to cell biology. Environ. Exp. Bot. 109, 212–228.

Ding, P., Rekhter, D., Ding, Y., Feussner, K., Busta, L., Haroth, S., Xu, S. et al. (2016) Characterization of a pipecolic acid biosynthesis pathway required for systemic acquired resistance. Plant Cell 28, 2603–2615.

Durrant, W.E. and Dong, X. (2004) Systemic acquired resistance. Annu. Rev. Phytopathol. 42, 185–209.

Fammartino, A., Cardinale, F., Göbel, C., Mène-Saffrané, L., Fournier, J., Feussner, I. and Esquerré-Tugayé, M.T. (2007) Characterization of a divinyl ether biosynthetic pathway specifically associated with pathogenesis in tobacco. Plant Physiol. 143, 378–388.

Funk, C.D., Laferrière, C. and Ardakani, A. (2021) Target product profile analysis of COVID-19 vaccines in phase III clinical trials and beyond: An early 2021 perspective. Viruses 13, 418.

Geng, P., Zhang, S., Liu, J., Zhao, C., Wu, J., Cao, Y., Fu, C. et al. (2020) MYB20, MYB42, MYB43, and MYB85 regulate phenylalanine and lignin biosynthesis during secondary cell wall formation. Plant Physiol. **182**, 1272-1283.

Gobeil, P., Pillet, S., Séguin, A., Boulay, I., Mahmood, A., Vinh, D.C., Charland, N. et al. (2021) Interim report of a phase 2 randomized trial of a plant-produced virus-like particle vaccine for Covid-19 in healthy adults aged 18-64 and older adults aged 65 and older. medRxiv doi: 10.1101/2021.05.14.21257248.

Goulet, M.C., Gaudreau, L., Gagné, M., Maltais, A.M., Laliberté, A.C., Éthier, G., Bechtold, N. et al. (2019) Production of biopharmaceuticals in *Nicotiana benthamiana* - Axillary stem growth as a key determinant of total protein yield. Front. Plant Sci. 10, 735.

Grosse-Holz, F., Kelly, S., Blaskowski, S., Kaschani, F., Kaiser, M. and van der Hoorn, R.A.L. (2018a) The transcriptome, extracellular proteome and active secretome of agroinfiltrated *Nicotiana benthamiana* uncover a large, diverse protease repertoire. Plant Biotechnol. J. 16, 1068–1084.

Grosse-Holz, F., Madeira, L., Zahid, M.A., Songer, M., Kourelis, J., Fesenko, M., Ninck, S. et al. (2018b) Three unrelated protease inhibitors enhance accumulation of pharmaceutical recombinant proteins in *Nicotiana benthamiana*. Plant Biotechnol. J. 16, 1797–1810.

Hager, K.J., Pérez Marc, G., Gobeil, P., Diaz, R.S., Heizer, G., Llapur, C., Makarkov, A.I. et al. (2022) Efficacy and safety of a recombinant plant-based adjuvanted Covid-19 vaccine. N. Engl. J. Med. 386, 2084–2096.

Hamel, L.P., Tardif, R., Poirier-Gravel, F., Rasoolizadeh, A., Brosseau, C., Giroux, G., Lucier, J.F. et al. (2024a) Molecular responses of agroinfiltrated *Nicotiana benthamiana* leaves expressing suppressor of silencing P19 and influenza virus-like particles. Plant Biotechnol. J. 22, 1078–1100.

Hamel, L.P., Comeau, M.A., Tardif, R., Poirier-Gravel, F., Paré, M.E., Lavoie, P.O., Goulet, M.C. et al. (2024b) Heterologous expression of influenza hemagglutinin leads to early and transient activation of the unfolded protein response in *Nicotiana benthamiana*. Plant Biotechnol. J. 22, 1146–1163.

Hamel, L.P., Paré, M.E., Poirier-Gravel, F., Tardif, R., Comeau, M.A., Lavoie, P.O., Langlois, A. et al. (2025) Expression of a constitutively active nitrate reductase increases SARS-CoV-2 Spike protein production in *Nicotiana benthamiana* leaves that otherwise show traits of senescence. *Submitted for revision*.

Howell, S.H. (2021) Evolution of the unfolded protein response in plants. Plant Cell Environ. 44, 2625–2635.

Huang, F.C., Studart-Witkowski, C. and Schwab, W. (2010) Overexpression of *hydroperoxide lyase* gene in *Nicotiana benthamiana* using a viral vector system. Plant Biotechnol. J. 8, 783–795.

Hussey, S.G., Mizrachi, E., Spokevicius, A.V., Bossinger, G., Berger, D.K. and Myburg, A, A. (2011) *SND2*, a NAC transcription factor gene, regulates genes involved in secondary cell wall development in *Arabidopsis* fibres and increases fibre cell area in *Eucalyptus*. BMC Plant Biol. 11, 173.

Ishihama, N., Yamada, R., Yoshioka, M., Katou, S. and Yoshioka, H. (2011) Phosphorylation of the *Nicotiana benthamiana* WRKY8 transcription factor by MAPK functions in the defense response. Plant Cell. 23, 1153–1170.

Jackson, C.B., Farzan, M., Chen, B. and Choe, H. (2022) Mechanisms of SARS-CoV-2 entry into cells. Nat. Rev. Mol. Cell Biol. 23, 3–20.

Kang, J.H., Wang, L., Giri, A. and Baldwin, I.T. (2006) Silencing *threonine deaminase* and *JAR4* in *Nicotiana attenuata* impairs jasmonic acid-isoleucine-mediated defenses against *Manduca sexta*. Plant Cell 18, 3303–3320.

Kim, D., Langmead, B. and Salzberg, S.L. (2015) HISAT: a fast spliced aligner with low memory requirements. Nat. Methods. 12, 357–360.

Kim, S.Y., Kim, S.G., Kim, Y.S., Seo, P.J., Bae, M., Yoon, H.K. and Park, C.M. (2007) Exploring membrane-associated NAC transcription factors in *Arabidopsis*: implications for membrane biology in genome regulation. Nucleic Acids Res. 35, 203–213.

Kim, S.H., Lam, P.Y., Lee, M.H., Jeon, H.S., Tobimatsu, Y. and Park, O.K. (2020) The *Arabidopsis* R2R3 MYB transcription factor MYB15 Is a key regulator of lignin biosynthesis in effector-riggered immunity. Front. Plant Sci. 11, 583153.

Landry, N., Ward, B.J., Trepanier, S., Montomoli, E., Dargis, M., Lapini, G. and Vezina, L.P. (2010) Preclinical and clinical development of plant-made virus-like particle vaccine against avian H5N1 influenza. PLoS One 5, e15559.

Landry, N., Pillet, S., Favre, D., Poulin, J.F., Trépanier, S., Yassine-Diab, B. and Ward, B.J. (2014) Influenza virus-like particle vaccines made in *Nicotiana benthamiana* elicit durable, poly-functional and cross-reactive T cell responses to influenza HA antigens. Clin. Immunol. 154, 164–177.

Lee, C.W., Efetova, M., Engelmann, J.C., Kramell, R., Wasternack, C., Ludwig-Müller, J., Hedrich, R. et al. (2009) *Agrobacterium tumefaciens* promotes tumor induction by modulating pathogen defense in *Arabidopsis thaliana*. Plant Cell 21, 2948–2962.

Lemonnier, P., Gaillard, C., Veillet, F., Verbeke, J., Lemoine, R., Coutos-Thévenot, P. and La Camera, S. (2014) Expression of *Arabidopsis* sugar transport protein STP13 differentially affects glucose transport activity and basal resistance to *Botrytis cinerea*. Plant Mol. Biol. 85, 473–484.

Li, D., Sempowski, G.D., Saunders, K.O., Acharya, P. and Haynes, B.F. (2022) SARS-CoV-2 neutralizing antibodies for COVID-19 prevention and treatment. Annu. Rev. Med. 73, 1–16.

Li, J., Brader, G. and Palva, E.T. (2008) Kunitz trypsin inhibitor: an antagonist of cell death triggered by phytopathogens and fumonisin b1 in *Arabidopsis*. Mol. Plant 1, 482–495.

Li, W., Li, X., Chao, J., Zhang, Z., Wang, W., and Guo, Y. (2018) NAC family transcription factors in tobacco and their potential role in regulating leaf senescence. Front. Plant Sci. 9, 1900.

Livak, K.J. and Schmittgen, T.D. (2001) Analysis of relative gene expression data using real-time quantitative PCR and the 2^-ΔΔCt^ Method. Methods 25, 402–408.

López-Marqués, R.L., Theorin, L., Palmgren, M.G. and Pomorski, T.G. (2014) P4- ATPases: lipid flippases in cell membranes. Pflügers Arch. 466, 1227–1240.

Lorenzo, O., Chico, J.M., Sánchez-Serrano, J.J. and Solano, R. (2004) *JASMONATE-INSENSITIVE1* encodes a MYC transcription factor essential to discriminate between different jasmonate-regulated defense responses in *Arabidopsis*. Plant Cell 16, 1938–1950.

McKenna, A., Hanna, M., Banks, E., Sivachenko, A., Cibulskis, K., Kernytsky, A., Garimella, K. et al. (2010) The Genome Analysis Toolkit: a MapReduce framework for analyzing next-generation DNA sequencing data. Genome Res. 20, 1297–1303.

McLellan, H., Boevink, P.C., Armstrong, M.R., Pritchard, L., Gomez, S., Morales, J., Whisson, S.C. et al. (2013) An RxLR effector from *Phytophthora infestans* prevents re-localisation of two plant NAC transcription factors from the endoplasmic reticulum to the nucleus. PLoS Pathog. 9, e1003670.

Morisseau, C. (2013) Role of epoxide hydrolases in lipid metabolism. Biochimie 95, 91–95.

Ninkuu, V., Zhang, L., Yan, J., Fu, Z., Yang, T. and Zeng, H. (2021) Biochemistry of terpenes and recent advances in plant protection. Int. J. Mol. Sci. 22, 5710.

Niu, F., Cui, X., Zhao, P., Sun, M., Yang, B., Deyholos, M.K., Li, Y. et al. (2020) WRKY42 transcription factor positively regulates leaf senescence through modulating SA and ROS synthesis in *Arabidopsis thaliana*. Plant J. 104, 171–184.

Ooka, H., Satoh, K., Doi, K., Nagata, T., Otomo, Y., Murakami, K., Matsubara, K. et al. (2003) Comprehensive analysis of NAC family genes in *Oryza sativa* and *Arabidopsis thaliana*. DNA Res. 10, 239–247.

Park, K.S., Cheong, J.J., Lee, S.J., Suh, M.C. and Choi, D. (2000) A novel proteinase inhibitor gene transiently induced by tobacco mosaic virus infection. Biochim. Biophys. Acta 1492, 509–512.

Piccoli, L., Park, Y.J., Tortorici, M.A., Czudnochowski, N., Walls, A.C., Beltramello, M., Silacci-Fregni, C. et al. (2020) Mapping neutralizing and immunodominant sites on the SARS-CoV-2 spike receptor-binding domain by structure-guided high-resolution serology. Cell 183, 1024–1042.

Pignocchi, C. and Foyer, C.H. (2003) Apoplastic ascorbate metabolism and its role in the regulation of cell signalling. Curr. Opin. Plant Biol. 6, 379–389.

Pignocchi, C., Kiddle, G., Hernandez, I., Foster, S.J., Asensi, A., Taybi, T., Barnes, J. et al. (2006) Ascorbate oxidase-dependent changes in the redox state of the apoplast modulate gene transcript accumulation leading to modified hormone signaling and orchestration of defense processes in tobacco. Plant Physiol. 141, 423–435.

Pillet, S., Aubin, É., Trépanier, S., Bussière, D., Dargis, M., Poulin, J.F., Yassine-Diab, B. et al. (2016) A plant-derived quadrivalent virus like particle influenza vaccine induces cross-reactive antibody and T cell response in healthy adults. Clin. Immunol. 168, 72–87.

Pillet, S., Couillard, J., Trépanier, S., Poulin, J.F., Yassine-Diab, B., Guy, B., Ward, B.J. et al. (2019) Immunogenicity and safety of a quadrivalent plant-derived virus like particle influenza vaccine candidate-two randomized phase II clinical trials in 18 to 49 and ≥50 years old adults. PLoS One. 14, e0216533.

Polack, F.P., Thomas, S.J., Kitchin, N., Absalon, J., Gurtman, A., Lockhart, S., Perez, J.L. et al. (2020) Safety and efficacy of the BNT162b2 mRNA Covid-19 vaccine. N. Engl. J. Med. 383, 2603–2615.

Ramos, R.N., Martin, G.B., Pombo, M.A. and Rosli, H.G. (2021) WRKY22 and WRKY25 transcription factors are positive regulators of defense responses in *Nicotiana benthamiana*. Plant Mol. Biol. 105, 65–82.

Rausch, T. and Greiner, S. (2004) Plant protein inhibitors of invertases. Biochim. Biophys. Acta 1696, 253–261.

Read, A. and Schröder, M. (2021) The unfolded protein response: an overview. Biology (Basel) 10, 384.

Ribot, C., Zimmerli, C., Farmer, E.E., Reymond, P. and Poirier, Y. (2008) Induction of the *Arabidopsis PHO1;H10* gene by 12-oxo-phytodienoic acid but not jasmonic acid via a CORONATINE INSENSITIVE1-dependent pathway. Plant Physiol. 147, 696–706.

Riedlmeier, M., Ghirardo, A., Wenig, M., Knappe, C., Koch, K., Georgii, E., Dey, S. et al. (2017) Monoterpenes support systemic acquired resistance within and between plants. Plant Cell 29, 1440–1459.

Robatzek, S. and Somssich, I.E. (2001) A new member of the *Arabidopsis* WRKY transcription factor family, *AtWRKY6*, is associated with both senescence- and defence-related processes. Plant J. **28**, 123-133.

Roth, M.G. (2008) Molecular mechanisms of PLD function in membrane traffic. Traffic 9, 1233–1239.

Rotshild, V., Hirsh-Raccah, B., Miskin, I., Muszkat, M. and Matok, I. (2021) Comparing the clinical efficacy of COVID-19 vaccines: a systematic review and network meta-analysis. Sci. Rep. 11, 22777.

Rowan, A.D., Brzin, J., Buttle, D.J. and Barrett, A.J. (1990) Inhibition of cysteine proteinases by a protein inhibitor from potato. FEBS Lett. 269, 328–330.

Sainsbury, F. and Lomonossoff, G.P. (2014) Transient expressions of synthetic biology in plants. Curr. Opin. Plant Biol. 19, 1–7.

Sangwan, S., Shameem, N., Yashveer, S., Tanwar, H., Parray, J.A., Jatav, H.S., Sharma, S. et al. (2022) Role of salicylic acid in combating heat stress in plants: insights into modulation of vital processes. Front. Biosci. (Landmark Ed) 27, 310.

Sette, A. and Crotty, S. (2021) Adaptive immunity to SARS-CoV-2 and COVID-19. Cell 184, 861–880.

Sierro, N., Battey, J.N.D., Ouadi, S., Bakaher, N., Bovet, L., Willig, A., Goepfert, S., et al. (2014) The tobacco genome sequence and its comparison with those of tomato and potato. Nat Commun. 5, 3833.

Silhavy, D., Molnár, A., Lucioli, A., Szittya, G., Hornyik, C., Tavazza, M. and Burgyán, J. (2002) A viral protein suppresses RNA silencing and binds silencing-generated, 21- to 25-nucleotide double-stranded RNAs. EMBO J. **21**, 3070-3080.

Song, J.T., Lu, H., McDowell, J.M. and Greenberg, J.T. (2004) A key role for *ALD1* in activation of local and systemic defenses in *Arabidopsis*. Plant J. 40, 200–212.

Stael, S., Van Breusegem, F., Gevaert, K. and Nowack, M.K. (2019) Plant proteases and programmed cell death. J. Exp. Bot. 70, 1991–1995.

Staswick, P.E. and Tiryaki, I. (2004) The oxylipin signal jasmonic acid is activated by an enzyme that conjugates it to isoleucine in *Arabidopsis*. Plant Cell 16, 2117–2127.

Stintzi, A., Weber, H., Reymond, P., Browse, J. and Farmer, E.E. (2001) Plant defense in the absence of jasmonic acid: the role of cyclopentenones. Proc. Natl. Acad. Sci. USA 98, 12837–12842.

Sun, L., Yang, Z.T., Song, Z.T., Wang, M.J., Sun, L., Lu, S.J. and Liu, J.X. (2013) The plant-specific transcription factor gene *NAC103* is induced by bZIP60 through a new *cis*-regulatory element to modulate the unfolded protein response in *Arabidopsis*. Plant J. 76, 274–286.

Taki, N., Sasaki-Sekimoto, Y., Obayashi, T., Kikuta, A., Kobayashi, K., Ainai, T., Yagi, K. et al. (2005) 12-oxo-phytodienoic acid triggers expression of a distinct set of genes and plays a role in wound-induced gene expression in *Arabidopsis*. Plant Physiol. 139, 1268–1283.

Thaler, J.S., Humphrey, P.T. and Whiteman, N.K. (2012) Evolution of jasmonate and salicylate signal crosstalk. Trends Plant Sci. 17, 260–270.

Torres, M.A., Dangl, J.L. and Jones, J.D. (2002) *Arabidopsis* gp91^phox^ homologues *AtrbohD* and *AtrbohF* are required for accumulation of reactive oxygen intermediates in the plant defense response. Proc. Natl. Acad. Sci. USA 99, 517–522.

Trapnell, C., Hendrickson, D.G., Sauvageau, M., Goff, L., Rinn, J.L. and Pachter, L. (2013) Differential analysis of gene regulation at transcript resolution with RNA-seq. Nat Biotechnol. 31, 46–53.

Vandesompele, J., De Preter, K., Pattyn, F., Poppe, B., Van Roy, N., De Paepe, A. and Speleman, F. (2002) Accurate normalization of real-time quantitative RT-PCR data by geometric averaging of multiple internal control genes. Genome Biol. 3, Research0034.1.

Veillet, F., Gaillard, C., Coutos-Thevenot, P. and La Camera, S. (2016) Targeting the *AtCWIN1* gene to explore the role of invertases in sucrose transport in roots and during *Botrytis cinerea* infection. Front. Plant Sci. 7, 1899.

Vicente, J., Cascón, T., Vicedo, B., García-Agustín, P., Hamberg, M. and Castresana, C. (2012) Role of 9-lipoxygenase and α-dioxygenase oxylipin pathways as modulators of local and systemic defense. Mol. Plant 5, 914–928.

Wang, K., Guo, Q., Froehlich, J.E., Hersh, H.L., Zienkiewicz, A., Howe, G.A. and Benning, C. (2018) Two abscisic acid-responsive plastid lipase genes involved in jasmonic acid biosynthesis in *Arabidopsis thaliana*. Plant Cell 30, 1006–1022.

Wang, L., Allmann, S., Wu, J. and Baldwin, I.T. (2008) Comparisons of LIPOXYGENASE3- and JASMONATE-RESISTANT4/6-silenced plants reveal that jasmonic acid and jasmonic acid-amino acid conjugates play different roles in herbivore resistance of *Nicotiana attenuata*. Plant Physiol. 146, 904–915.

Wang, Y., Chantreau, M., Sibout, R. and Hawkins, S. (2013) Plant cell wall lignification and monolignol metabolism. Front. Plant Sci. 4, 220.

Ward, B.J., Gobeil, P., Séguin, A., Atkins, J., Boulay, I., Charbonneau, P.Y., Couture, M. et al. (2021) Phase 1 randomized trial of a plant-derived virus-like particle vaccine for COVID-19. Nat. Med. 27, 1071–1078.

Ward, B.J., Makarkov, A., Séguin, A., Pillet, S., Trépanier, S., Dhaliwall, J., Libman, M.D. et al. (2020) Efficacy, immunogenicity, and safety of a plant-derived, quadrivalent, virus-like particle influenza vaccine in adults (18-64 years) and older adults (≥65 years): two multicentre, randomised phase 3 trials. Lancet 396, 1491–1503.

Wasternack, C. and Feussner, I. (2018) The oxylipin pathways: biochemistry and function. Annu. Rev. Plant Biol. 69, 363–386.

Wrapp, D., Wang, N., Corbett, K.S., Goldsmith, J.A., Hsieh, C.L., Abiona, O., Graham, B.S. et al. (2020) Cryo-EM structure of the 2019-nCoV spike in the prefusion conformation. Science 367, 1260–1263.

Wu, A., Allu, A.D., Garapati, P., Siddiqui, H., Dortay, H., Zanor, M.I., Asensi-Fabado, M.A. et al. (2012) *JUNGBRUNNEN1*, a reactive oxygen species-responsive NAC transcription factor, regulates longevity in *Arabidopsis*. Plant Cell 24, 482–506.

Yang, Z.T., Fan, S.X., Wang, J.J., An, Y., Guo, Z.Q., Li, K. and Liu, J.X. (2023) The plasma membrane-associated transcription factor NAC091 regulates unfolded protein response in *Arabidopsis thaliana*. Plant Sci. 334, 111777.

Yang, Z.T., Lu, S.J., Wang, M.J., Bi, D.L., Sun, L., Zhou, S.F., Song, Z.T. et al. (2014) A plasma membrane-tethered transcription factor, NAC062/ANAC062/NTL6, mediates the unfolded protein response in *Arabidopsis*. Plant J. **79**, 1033-1043.

Zhang, Y., Xu, S., Ding, P., Wang, D., Cheng, Y.T., He, J., Gao, M. et al. (2010) Control of salicylic acid synthesis and systemic acquired resistance by two members of a plant-specific family of transcription factors. Proc. Natl. Acad. Sci. USA 107, 18220–18225.

Zheng, X.Y., Spivey, N.W., Zeng, W., Liu, P.P., Fu, Z.Q., Klessig, D.F., He, S.Y. et al. (2012) Coronatine promotes *Pseudomonas syringae* virulence in plants by activating a signaling cascade that inhibits salicylic acid accumulation. Cell. Host Microbe 11, 587–596.

Zheng, Z., Qamar, S.A., Chen, Z. and Mengiste, T. (2006) *Arabidopsis* WRKY33 transcription factor is required for resistance to necrotrophic fungal pathogens. Plant J. 48, 592–605.

Zhong, R., Lee, C., Zhou, J., McCarthy, R.L. and Ye, Z.H. (2008) A battery of transcription factors involved in the regulation of secondary cell wall biosynthesis in *Arabidopsis*. Plant Cell 20, 2763–2782.

Zhou, P., Yang, X.L., Wang, X.G., Hu, B., Zhang, L., Zhang, W., Si, H.R. et al. (2020) A pneumonia outbreak associated with a new coronavirus of probable bat origin. Nature 579, 270–273.

Zhou, X., Jiang, Y. and Yu, D. (2011) WRKY22 transcription factor mediates dark-induced leaf senescence in *Arabidopsis*. Mol. Cells 31, 303–313.

Zhu, N., Zhang, D., Wang, W., Li, X., Yang, B., Song, J., Zhao, X. et al. (2020) A novel coronavirus from patients with pneumonia in China, 2019. N. Engl. J. Med. **382**, 727-733.

