## Appendix S1 for "Molecular changes in agroinfiltrated leaves of *Nicotiana benthamiana* expressing suppressor of silencing P19 and coronavirus-like particles"

**Definitions of all abbreviations** **used in this study:**

ABI5	Abscisic acid-Insensitive 5

ACAT	Acetoacetyl-CoA thiolase

ACC	1-aminocyclopropane-1-carboxylate

ACE2	Angiotensin-converting enzyme 2

ACO	ACC oxidase

ACS	ACC synthase

ACT1	Actin 1

ACX	Acyl-CoA oxidase

ALD1	AGD2-Like Defense 1

ALA	Aminophospholipid ATPase

ALIS	ALA-Interacting Subunit

AO	Ascorbate oxidase

AOC	Allene oxide cyclase

AOS	Allene oxide synthase

AP2/ERF	Apetala2/Ethylene Responsive Factor

AsA	Ascorbic acid

ASN	Asparagine synthetase

At	*Arabidopsis thaliana*

ATAF1	ARABIDOPSIS ACTIVATION FACTOR 1

ATAF2	ARABIDOPSIS ACTIVATION FACTOR 2

ATP	Adenosine triphosphate

AZI	Azelaic acid-induced

BBE	Berberine-bridge enzyme

bHLH	Basic helix-loop-helix

BiP	Binding immunoglobulin Protein

bZIP	Basic leucine zipper

CAD	Cinnamyl alcohol dehydrogenase

CaMV	Cauliflower mosaic virus

CCA1	CIRCADIAN CLOCK-ASSOCIATED 1

CCR	Cinnamoyl-CoA reductase

CCoAOMT	Caffeoyl-CoA O-methyltransferase

cDNA	complementary DNA

C3H	Coumarate-3-hydroxylase

C4H	Cinnamate-4-hydroxylase

CHI	Chitinase

CHLD	Magnesium-chelatase subunit D

CHLH	Magnesium-chelatase subunit H

CHLM	Magnesium-protoporphyrin IX methyltransferase

CIF	Cell wall inhibitor of β-fructosidase

4CL	4-coumarate-CoA ligase

CNX	Calnexin

CO_2_	Carbon dioxide

COMT	Caffeic acid 3-O-methyltransferase

COVID-19	Coronavirus disease 2019

CoVLP	Coronavirus-like particle

CP	Cysteine protease

CPI	Cysteine Protease Inhibitor

CRT	Calreticulin

CT	Cytosolic tail

Ct	Cycle threshold

CTE	C-terminal extension

CUP2	CUP SHAPED COTYLEDON 2

CWI	Cell wall invertase

CYP71	Cytochrome P450 of family 71

CYP74	Cytochrome P450 of family 74

D	Day

DER	Derlin

DES	Divinyl ether synthase

DGK	Diacylglycerol kinase

DIA	Direct data independent

DNA	Deoxyribonucleic acid

DOX	α-dioxygenase

DPI	Days post-infiltration

DREB	Dehydration-responsive element-binding protein

EAS	Epoxyalcohol synthase

EBI	European Bioinformatics Institute

EIN2	ETHYLENE-INSENSITIVE 2

EIN3	ETHYLENE-INSENSITIVE 3

EF1-α	Elongation Factor 1α

EH	Epoxide hydrolase

EMBL	European Molecular Biology Laboratory

ENA	European Nucleotide Archive

ER	Endoplasmic reticulum

ERAD	ER-associated degradation

ERQC	ER quality control

EV	Empty vector

FC	Fold change

FDR	False discovery rate

F5H	Ferulate-5-hydroxylase

FPKM	Fragments per kilobase of transcript per million mapped reads

FPPS	Farnesyl pyrophosphate synthase

GAPDH	Glyceraldehyde 3-phosphate dehydrogenase

GATK	Genome Analysis Toolkit

gBlock	Gene block

GEO	Gene Expression Omnibus

GGPPS	Geranyl geranyl diphosphate synthase

GLK	Golden 2-Like protein

GLN1	Cytosolic glutamine synthetase

GLN2	Chloroplastic glutamine synthetase

GLT	NADH-dependent glutamate synthase

GLU	Chloroplastic ferredoxin-dependent glutamate synthase

GLV	Green leaf volatile

GPAT	Glycerol-3-phosphate acyltransferase

GPPS	Geranyl diphosphate synthase

GRP94	Glucose-regulated protein 94

GUN4	GENOMES UNCOUPLED 4

HA	Haemagglutinin

HCT	Hydroxycinnamoyl-CoA transferase

HEMA1	Heme synthesis A 1 (glutamyl-tRNA reductase 1)

HEMA2	Heme synthesis A 2 (glutamyl-tRNA reductase 2)

HIN1	Harpin-induced 1

HMGR	3-hydroxy-3-methylglutaryl-CoA reductase

HMGS	3-hydroxy-3-methylglutaryl-CoA synthase

HPL	Hydroperoxide lyase

HSP	Heat shock protein

ICS	Isochorismate synthase

Ile	Isoleucine

IPCS	inositolphosphorylceramide synthase

IPPI	Isopentyl diphosphate isomerase

iTRAQ	Isobaric tags for relative and absolute quantitation

JA	Jasmonic acid

JA-Ile	Jasmonate-isoleucine

JAR4	Jasmonate-Resistant 4

JAR6	Jasmonate-Resistant 6

JUB1	JUNGBRUNNEN1

KTI	Kunitz trypsin inhibitor

LAC	Laccase

LB	Luria-Bertani

LHCB	Light-harvesting chlorophyll a/b binding protein

LKR1	Lys ketoglutarate reductase 1

Log_2_FC	Log_2_ of the expression FC value

9-LOX	9-lipoxygenase

13-LOX	13-lipoxygenase

LR	Linear regression

LURP1	Late up-regulated in response to *Hyaloperonospora parasitica* 1

MAPK	Mitogen-activated protein kinase

MCA	Methylcoumarin

MEP	Methyl erythritol phosphate

MVK	Mevalonate kinase

MLO1	Mildew Resistance Locus O 1

MLO6	Mildew Resistance Locus O 6

Na	*Nicotiana alata*

NAC	NAM, ATAF1, and CUC2

NADH	Nicotinamide adenine dinucleotide

NAM	NO APICAL MERISTEM

Nb	*Nicotiana benthamiana*

Ng	*Nicotiana glutinosa*

NI	Non-infiltrated

NiR	Nitrite reductase

NOS	Nopaline synthase

NR	Nitrate reductase

Nt	*Nicotiana tabacum*

NTP1	NAC Targeted by *Phytophthora* 1

OPDA	12-oxophytodienoic acid

OPR	OPDA reductase

ORE1	ORESARA 1

OS9	Osteosarcoma 9

P	Primary

padj	Adjusted p-value

PAL	Phenylalanine ammonia lyase

PaO	pheophorbide a oxygenase

PAT	Patatin

PC	Principal component

PCA	Principal component analysis

PCD	Plant cell death

PDF	Plant defensin

PDI	Protein disulfide isomerase

PI	Protease inhibitor

PLA	Phospholipase A

PLC	Phospholipase C

PLD	Phospholipase D

PLIP	PLASTID LIPASE

PM	Plasma membrane

PMD	Mevalonate diphosphate decarboxylase

PMK	Phosphomevalonate kinase

PPH	Pheophytinase

PPO	Polyphenol oxidase

PR1	Pathogenesis-Related 1

PR2	Pathogenesis-Related 2

PR3	Pathogenesis-Related 3

PR4	Pathogenesis-Related 4

PRX	Peroxidase

PrxQ	Peroxiredoxin Q

R^2^	Coefficient of determination

RbcL	RuBisCO large subunit

RbcS	RuBisCO small subunit

RBD	Receptor-binding domain

RBOH	Respiratory burst oxidase homolog

RD19	RESPONSIVE TO DEHYDRATION 19

RD21	RESPONSIVE TO DEHYDRATION 21

RNA	Ribonucleic acid

mRNA	Messenger RNA

ROS	Reactive oxygen species

RuBisCO	Ribulose-1,5-bisphosphate carboxylase/oxygenase

RTqPCR	Real time quantitative polymerase chain reaction

S	Spike (or secondary)

SA	Salicylic acid

SAG	Senescence-associated gene

SAG2	Senescence-associated gene 2

SAG12	Senescence-associated gene 12

SAGT1	SA glucosyl transferase 1

SAM	S-adenosylmethionine

SAMS	SAM synthase

SAND	Sp100, AIRE-1, NucP41/75, DEAF-1

SAR	Systemic acquired resistance

SARD1	SAR DEFICIENT 1

SARD4	SAR DEFICIENT 4

SARS-CoV-2	Severe Acute Respiratory Syndrome Coronavirus 2

SDS-PAGE	Sodium dodecyl sulfate-polyacrylamide gel electrophoresis

SGR1	STAYGREEN 1

Sl	*Solanum lycopersicum*

SND2	Secondary Wall-Associated NAC Domain 2

SND3	Secondary Wall-Associated NAC Domain 3

SoEP	Sexual Organs Expressed Protein

SOX1	Sarcosine oxidase 1

SP	Signal peptide

St	*Solanum tuberosum*

StEP	Stigma Expressed Protein

STP	Sugar transport protein

TBS	Tris-Buffered Saline

TBSV	Tomato bushy stunt virus

TD	Threonine deaminase

TEM	Transmission electron microscopy

TF	Transcription factor

TGA	TGACG-binding protein

TM	Transmembrane

TMD	Transmembrane domain

TPS	Terpene synthase

UBQ1	Ubiquitin 1

UPR	Unfolded protein response

UTR	Untranslated region

VATP	Vacuolar ATPase

VLP	Virus-like particle

VOC	Volatile organic compound

VSR	Viral suppressor of RNA silencing

WHO	World Health Organization

ZFP	Zinc finger protein
